# Contingent Convergence: The ability to detect convergent genomic evolution is dependent on population size and migration

**DOI:** 10.1101/592105

**Authors:** James R. Whiting, Bonnie A. Fraser

## Abstract

Outlier scans, in which the genome is scanned for signatures of selection, have become a prominent tool in studies of local adaptation, and more recently studies of genetic convergence in natural populations. However, such methods have the potential to be confounded by features of demographic history, such as population size and migration, which are considerably varied across natural populations. In this study, we use forward-simulations to investigate and illustrate how several measures of genetic differentiation commonly used in outlier scans (F_ST_, D_XY_ and Δπ) are influenced by demographic variation across multiple sampling generations. In a factorial design with 16 treatments, we manipulate the presence/absence of founding bottlenecks (N of founding individuals), protracted bottlenecks (proportional size of diverging population) and migration rate between two populations with ancestral and derived phenotypic optima. Our results illustrate known constraints of individual measures associated with reduced population size and a lack of migration; but notably we demonstrate how relationships between measures are similarly dependent on demography. We find that false-positive signals of convergent evolution (the same simulated outliers detected in independent treatments) are attainable as a product of similar demographic treatment, and that outliers across different measures (particularly F_ST_ and D_XY_) can occur with little influence of selection. Taken together, we show how underappreciated, yet quantifiable measures of demographic history can influence commonly employed methods for detecting selection.

## INTRODUCTION

Studies assessing adaptation and evolution across the genome are increasing in popularity with the availability of modern sequencing technologies. These studies often centre around quantifying patterns of variation in genome-wide SNPs, which can be used to highlight regions or genes having experienced selection relative to the neutral backdrop of the rest of the genome. These analyses, which we refer to here as outlier scans, have become a common tool in population genetics and have seen great success across diverse, natural systems in identifying candidate genes associated with the evolution of a range of adaptive traits (reviewed recently by (Ahrens et al. 2018)). More recently, the method of overlapping outlier scans across independent lineages has been employed to test whether the same regions are involved in independent adaptation events (i.e. genetic convergent evolution) (reviewed by (Fraser and Whiting 2019)). This study seeks to investigate how different outlier scan methods are influenced by demographic variation in natural populations, and how this may lead to overlapping false-positives.

Recent discussions have highlighted the propensity of outlier scans to yield false-positives, given that outliers caused by heterogeneous genomic landscapes are commonplace irrespective of selection (Ellegren and Wolf 2017). For example, background selection (BGS), whereby linkage between neutral and deleterious variants reduces local diversity (B. Charlesworth, Morgan, and Charlesworth 1993; Bank et al. 2014; Burri 2017), has been invoked to offer alternative explanations for patterns at first attributed to directional selection (Cruickshank and Hahn 2014). Furthermore, the influence of neutral processes such as genetic drift (B. Charlesworth 2009) can similarly produce elevated genetic divergence and false-positive signatures of selection.

The strength of these processes is dependent on demography. For example, the influence of neutral processes and BGS should be more pronounced in smaller populations (low N_e_) (D. Charlesworth, Charlesworth, and Morgan 1995; Cutter and Payseur 2013; B. Charlesworth 2009; Yeaman and Otto 2011); as has been demonstrated in humans (Torres, Szpiech, and Hernandez 2018) and *Drosophila* (Sella et al. 2009; B. Charlesworth 1996). This relationship, however, may not be simply linear, with additional simulation evidence suggesting the effects of BGS are strongest at intermediate-low N_e_, and weaker at very low or high N_e_ (Zeng 2013).

Lack of connectivity among populations may also elevate measures of genetic differentiation, if a large global population is composed of smaller, isolated, sub-divided populations that each experience the effects of reduced N_e_ (B. Charlesworth, Nordborg, and Charlesworth 1997; Hoban et al. 2016). In an attempt to mitigate the likelihood of false-positives, some have advocated using multiple measures of population differentiation and divergence to identify regions of the genome likely to be under selection (B. Charlesworth 1998; Cruickshank and Hahn 2014). However, how these measures are correlated with each other, and with selection under different demographic scenarios, has not been explored.

A diverse array of measures of genetic differentiation and divergence have been employed for outlier scans, but here we focus on two of the most common measures of relative differentiation (F_ST_ and changes in nucleotide diversity [Δπ]) and a measure of absolute divergence (D_XY_). The most commonly used measure in outlier scans is the fixation-index F_ST_ (Weir and Cockerham 1984; Hudson, Slatkin, and Maddison 1992), which measures the relative amount of within- and between-population variance. This measure is therefore maximised when genomic regions exhibit the lowest within- and highest between-population variance. F_ST_ outliers at the right-tail of the distribution are considered candidates for adaptation because they reflect regions with large differences in allele frequency or high substitution rate (large between-population variance) and/or low nucleotide variation (low within-population variance) relative to the rest of the genome. Changes in nucleotide diversity (Δπ) are another indicator of adaptation, as selection on a beneficial allele limits variation within a population resulting in selective sweeps (Smith and Haigh 1974). Comparisons of the ratio of π between diverging populations reveal regions under selection, as local π is reduced in one population in comparison to the other. Whilst similar to F_ST_, Δπ does not discriminate between which copy of a polymorphism is fixed, such that a substitution between populations is equivalent to a common non-polymorphic site. Δπ outliers therefore represent regions of the genome with reduced π in either population relative to the rest of the genome. As a measure of absolute genetic divergence, D_XY_ (Nei 1987) does not consider the relative frequencies of polymorphisms within populations (B. Charlesworth 1998; Cruickshank and Hahn 2014). D_XY_ can be quantified as the average number of pairwise differences between sequence comparisons between two populations. This measure is therefore influenced by ancestral π and the substitution rate, so D_XY_ outliers highlight regions that are highly variable ancestrally, or in either population (large π), or exhibit many substitutions and thus increased sequence divergence.

Because each measure of differentiation/divergence (hereafter referred to collectively as measures of divergence) quantifies genetic variation between populations in a slightly different way, each has a unique relationship with demography. Whilst we can predict how individual measures are influenced by demography, and subsequently neutral processes, we know little about how different relationships between measures of divergence and demography affect their combined usage. We expect then that the utility of using multiple measures of divergence to detect selection may vary with demography.

This complex interplay between divergence measures and demography may be further exacerbated in studies that compare scans from multiple populations to identify convergent genomic evolution. Here, researchers use measures of population divergence across independent pairs of evolutionary replicates with outlier loci compared across results. This strategy has been employed extensively across diverse taxa, including: birds (Cooper and Uy 2017), fish (Hohenlohe et al. 2010; Jones, Chan, et al. 2012; Rougemont et al. 2017; Fraser et al. 2015; Reid et al. 2016; Meier et al. 2018), insects (Soria-Carrasco et al. 2014; Van Belleghem et al. 2018), mammals (Waterhouse et al. 2018), molluscs (Ravinet et al. 2016; Westram et al. 2014) and plants (Trucchi et al. 2017; Roda et al. 2013). For the sake of consistency, it is common to infer outlier loci within each replicate pair through a common method, but if replicates differ in their demographic histories then the applicability/power of that common method will also vary accordingly. This begs the question, can demographic variation among replicates alone explain signals of, or lack of, convergence?

Here, we used forward-simulations to investigate the effects of different demographic histories on the relationships between measures of divergence with selection. We varied demography through manipulating the number of founding individuals (founding bottlenecks), the population size of the diverging population (protracted bottlenecks) and the presence/absence of migration. We simulated the effects of selection on 25kb windows, each with a gene designed from features taken from the guppy (*Poecilia reticulata*) genome assembly, a prominent model system for studies of convergent evolution (Fraser et al. 2015; Reznick and Endler 1982). Moreover, we measured genetic divergence at 12 set time points through F_ST_, D_XY_ and Δπ between diverging populations to examine temporal relationships. Our aim was to investigate the significance of demographic history when employing outlier scans and highlight demographic scenarios that are susceptible to false-positives. We also aimed to test the occurrence of common outliers across measures, and whether overlapping outliers are consistently good indicators of selection. We sought to answer the following questions: 1) How do demographic factors influence measures of divergence through time? 2) How well do measures of divergence identify regions of the genome under strong selection through time and across demographic factors? 3) How are the different measures of divergence related through time and across demographic factors? 4) Where do we detect the strongest signals of convergence (i.e. overlapping outliers using single or multiple measures), and are these consistent with selection?

## METHODS

The forward-simulation software SLiM 3.0 (Haller and Messer 2019) was used to simulate population divergence under contrasting demographic treatments in a fully factorial design between a phenotypically-ancestral population (AP) of N = 1000 and phenotypically-derived population (DP), with a mutation rate based on the guppy genome (4.89e^-8^) (Künstner et al. 2016) and scaled 100-fold over three raw mutation rates (4.89e^-5^, 4.89e^-6^, 4.89e^-7^) to ensure robustness of results across different, more realistic values of θ (4N_e_μ). These scaled mutation rates therefore represent effective population sizes of 10-, 100-, and 1000-times greater than the number of individuals in our simulations (N = 1000), in line with estimates from other species (B. Charlesworth 2009). The main text reflects results for the intermediate mutation rate 4.89e^-6^, with others presented in supplementary figures. SLiM employs a classic Wright-Fisher model to simulate populations, in which a population of diploid hermaphrodites proceeds through generations such that an individual’s contribution towards the next generation is proportional to its relative fitness.

Demographic treatments included reducing DP size relative to the AP size (N = 1000) (0.01, 0.1, 0.5, 1.0), migration as a proportion of individuals exchanged between populations (0.0, 0.002) and founding bottlenecks as the number of individuals sampled from the burn-in population to construct DP genomes (100, 1000). For example, a demographic history with founding bottleneck = 100 individuals, DP size = 0.01, and migration = 0.002 would represent the following scenario: 1) At generation 1, following a burn-in period of 10,000 generations to reach mutation-selection balance, populations split as 100 genomes are sampled from the burn-in population of 1000 to form DP, whilst AP is formed by sampling all 1000 burn-in genomes; 2) At the next generation (and for the remainder of the simulation), DP size reflects the protracted bottleneck treatment, in this case 10 (0.01 of 1000); 3) AP and DP experience migration in both directions for the remainder of the simulation. Thus, each of our treatments can be thought of as a manipulation of the following: Founding bottlenecks limit the amount of variation within the burn-in population that is available to found DP; Protracted bottlenecks limit the size of DP, reducing mutational input and moderating the strength of neutral evolution and efficacy of selection; and migration dictates the presence of migration between AP and DP (Figure 1). The total N of individuals (AP + DP) within simulations varies between 1010 and 2000 depending on DP size parameter, however this potential expansion does not impact our results (Supporting Information).

**Figure 1:**
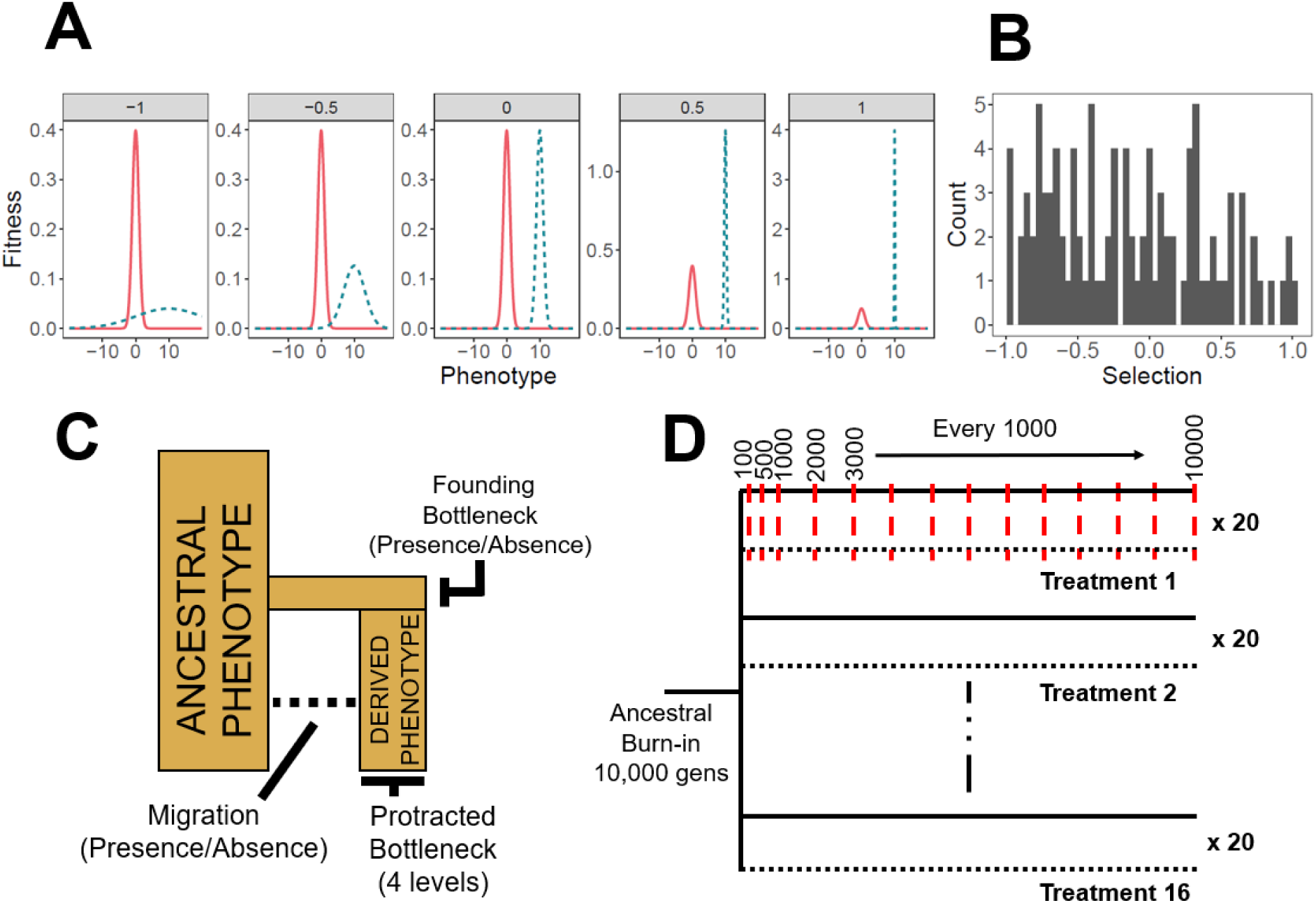
Experimental design for simulations. **A)** Examples of selection treatments experienced by genes from across the range, illustrated as the relationship between relative fitness of an individual and its phenotype. Each facet represents a different fitness landscape modified through editing the standard deviation of the normal distribution of fitness consequences in DP (blue, dashed). Facet labels constitute *S*, which were transformed as 10^-^*^S^* to give fitness function standard deviations (*S*_σ_) The x-axes represent phenotype as calculated through non-synonymous mutations and the y-axes represent relative fitness of individuals. The ancestral phenotype of AP (red, solid) is the same in all treatments (mean = 0, σ = 1), whilst DPs have a diverged phenotypic optimum (mean = 10, σ = *S*_σ_). **B)** Distribution of *S* values applied to genes (1 per gene) **C)** Demographic representation of treatment factors. **D)** Representation of simulation timeline for treatments, illustrating that all treatments share an ancestral burn-in population before splitting into 16 replicated “AP” (solid) and “DP” (dotted) population pairs. Red, dashed lines denote sampling generations at which F_ST_, D_XY_ and Δπ are calculated and averaged across the preceding 20 generations. The purpose of this averaging was to achieve a general sense of population differentiation at sampling points, such that values represent stable patterns rather than stochastic generation to generation variation.

Combining these treatment levels in a fully factorial experimental design generated a total of 16 different, independent demographic histories that all 25kb gene windows experienced. This factorial design allows us to examine the relative influence of founding bottlenecks, protracted bottlenecks and migration, but our study is limited to these features of demographic history and does not extend to population expansions, more complex cases of migration, or demographic fluctuations through time. In some cases, the influence of protracted bottlenecks renders the effects of founding bottlenecks unnecessary. However, when DP size is greater than 100, the inclusion of the founding bottleneck allows us to compare populations with equivalent mutational input but different standing variation.

SLiM runs were performed independently over 25kb genomic windows with a central gene that varied in length and exon content; with exon number, lengths, and intron lengths drawn at random from the guppy genome gene annotation file (gff). Recombination rate was set at a constant of 1e^-8^ across all windows.

Selection (*S*) in our model was represented as the fitness consequences incurred through distance from a phenotypic optimum in a one-dimensional fitness landscape. Selection in Wright-Fisher models is soft, in that low fitness individuals are not removed but have a lower likelihood of contributing to the next generation. Phenotypic optima were maintained over the course of simulations; thus, selection was constant throughout. This setup can be considered as analogous to two environments with contrasting optimum trait values for a single trait. Intensity of selection was manipulated by modifying the standard deviation (*S*_σ_) of the normal distribution curve from which the density distribution was calculated through the “dnorm” function in SLiM. Values for *S* were drawn from a continuous distribution between -1.00 and 1.00 and transformed such that *S*_σ_ = 10^-*S*^, yielding values between 0.1 and 10, with 0.1 representing the steepest fitness peak (*S* = 1) and 10 the shallowest (*S* = -1) (Figure 1A). Phenotypes were calculated per individual, per gene, as the additive phenotypic effects of exonic non-synonymous mutations, which appeared at a rate of 7/3 relative to synonymous mutations (assuming that most mutations in the third base of a codon do not alter the amino acid). Additive genetic variance was assessed due to its prevalence in complex traits in nature (Hill, Goddard, and Visscher 2008). Effect sizes for mutations with phenotypic effects were drawn from a Gaussian distribution with mean = 0 and σ = 1. The remaining synonymous exon mutations, and mutations in introns and outside of genes, had no effect on fitness.

For each simulation, populations were seeded with 1000 individuals and allowed to proceed for a burn-in period of 5*2N (10,000) generations to reach mutation-selection-migration balance. During this period, burn-in populations evolved towards the ancestral phenotypic optimum of 0, defined as a normal distribution with mean = 0 and σ = 1.0 (Figure 1A). Burn-in populations were then subjected to each demographic treatment to simulate the founding of multiple populations from a shared ancestral state. During this ‘divergence period’, AP continued to evolve around the ancestral optimum of 0, whilst DP’s phenotypic optimum was centred around 10, with fitness consequences defined according to *S*_σ_. Individuals also experienced fitness costs associated with phenotypic proximity to other individuals within the population as a proxy for competition and to ensure a realistic amount of phenotypic variation persisted within populations. Fitness costs due to competitive proximity were scaled to a maximum value of 1 with σ = 0.4 and occurred reciprocally between local individuals with phenotypes with a difference of ≤ 1.2 (3 * 0.4). Each gene was then run across all demographic treatment levels in parallel. Simulations were sampled at 100, 500, 1000, and then every 1000 generations up to 10,000 (Figure 1C). F_ST_ was calculated across the window as one minus the proportion of subpopulation heterozygosity (*H_S_*) relative to total heterozygosity (*H_T_*) (according to Hudson, Slatkin and Maddison (1992)). D_XY_ was calculated as the sum of nucleotide differences (*d_ij_*) between the *i*^th^ haplotype from AP and the *j*^th^ haplotype from DP (according to Nei (1987)). Mean heterozygosity (π) for each population was calculated as a single measure across the 25kb window. At each sampling point, each measure was calculated and averaged across the preceding 20 generations. Averaging was performed such that measures would not be dramatically biased by events occurring within individual generations. The change in mean heterozygosity between AP and DP (Δπ) was calculated as the log_10_-transformed ratio of π_AP_ to π_DP_, such that reduced diversity in the DP population increases the value of Δπ. Statistics were calculated over all monomorphic and polymorphic sites of 25kb windows. This has been designed to replicate genome scans that use sliding-approaches, with each 25kb region representing an independent window.

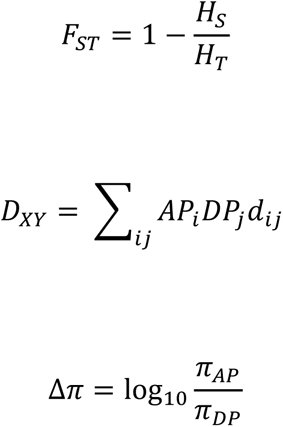

In total, 100 unique 25kb windows were simulated across 16 demographic treatments across three mutation rates, with results for the intermediate mutation rate presented in the main text. To account for stochastic noise in the simulation, each 25kb window was iterated 20 times for each demographic treatment. Simulations with divergent AP and DP phenotypes are referred to as “Pheno_Div_” simulations. Additional simulations were performed in which both populations shared the ancestral phenotype (“Pheno_Null_” simulations) and in which all sites were neutral with no selection imposed (“Neutral” simulations). The former of these was used to assess whether patterns associated with selection were driven by phenotypic divergence or variable stabilising selection within populations, whilst the latter was used to disentangle effects of selection and neutral processes such as drift. A common library of 100 25kb windows were used for all simulations. Outliers for simulations were taken as upper 95% quantiles.

All data analysis was performed in R (3.5) (R Core Team 2016). To assess relationships between divergence measures and selection, data were grouped by sampling generation within each treatment group (N = 16) and Pearson’s correlation coefficients were calculated between measures of divergence and selection (*S*) at each sampling point for all windows (N = 100). Correlation coefficients were then grouped within specific treatment levels (e.g. DP size = 0.5, or migration = 0.002) and averaged to give a coefficient reflecting each specific treatment level. These correlation coefficients were calculated for each iteration (N = 20) and averaged over to give final values.

To assess the effects of treatments on detecting outliers, we compared distributions and 95% cut-offs within each treatment for each measure of divergence for Pheno_Div_, Pheno_Null_, and Neutral simulations. We limited this analysis to early (100, 500), intermediate (3000) and late (10,000) sampling generations. Here, data from all iterations of each gene were pooled. To calculate false positive (FPR) and false negative rates (FNR), we pooled Neutral simulation data within treatment groups (20 iterations of 100 genes, N = 2000), removed a random set of iterations (N = 100), and replaced it with an assortment of random iterations of each gene (e.g. Gene 1/Iteration 2, Gene 2/Iteration 14… Gene 100/Iteration 4) from Pheno_Div_ data. We calculated FPR as the proportion of data above the 95% quantile for each measure of divergence/differentiation that came from neutral simulations. For single measures, FPR = FNR as 5% of data in each permutation comes from Pheno_Div_. We combined outlier sets across all combinations of F_ST_, D_XY_ and Δπ and examined neutral proportions within outlier sets to determine FPR. FNR for combined outlier sets corresponded to the proportion of Pheno_Div_ data not recovered in the combined outlier sets. These permutations were performed 100 times with results averaged. Proportional overlap of outlier sets was also calculated and compared across demographic treatment groups to examine convergence of results across treatments. Overlap was calculated within each permutation, averaged over, and visualised using heatmaps with hierarchical clustering of axes.

To examine how simulated gene features influenced patterns of genetic divergence, we used a linear mixed modelling (LMM) approach with gene ID and treatment group as random factors. Gene features modelled as independent factors were: number of exons (Exon N), gene size, the proportion of gene that is coding (selection target %), selection applied to each gene (*S*), and the generation at which the optimum phenotype was reached (Pheno Gen).

## RESULTS

### Demography modifies measures of divergence

Founding bottlenecks, simulated by reducing the number of possible founding genomes to a random 10% of the ancestral population, had little measurable effect on F_ST_, D_XY_ or Δπ, producing minimal variance between treatments over all sampling points (Figure 2A).

**Figure 2:**
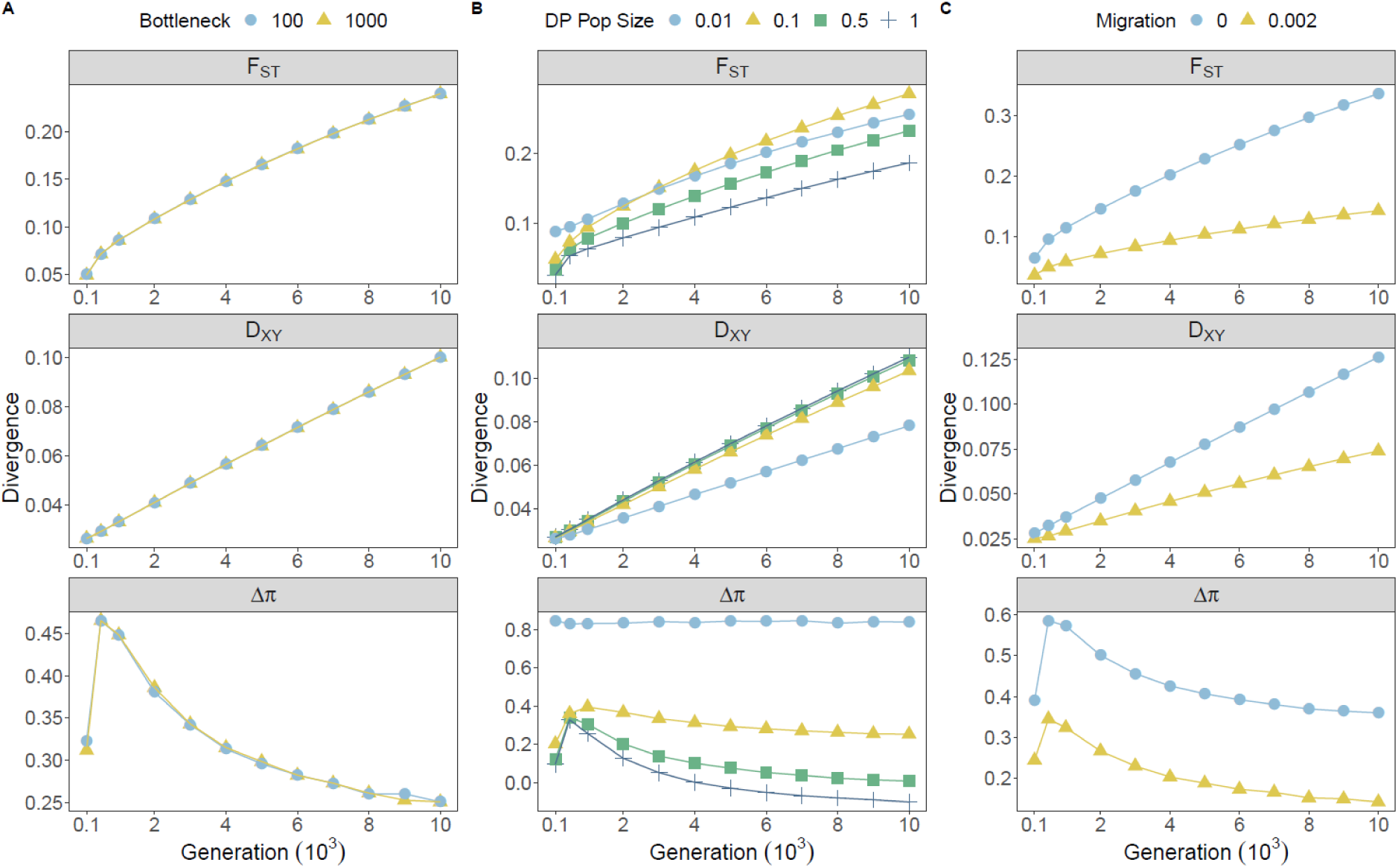
Effects of demographic treatments on measures of genetic divergence across all sampling generations. Point colour and shape denote treatment groups for founding bottlenecks (**A**), protracted bottlenecks (**B**) and migration (**C**). Each point represents values of divergence averaged across all genes within individual treatments groups, averaged within treatment levels, and averaged over 20 iterations.

Protracted bottlenecks, simulated by modifying the stable number of individuals within DP, had pronounced and variable effects on all measures. F_ST_ increased with reductions in DP size. This effect was generally consistent through time, although variance between treatments increased gradually through time (Figure 2B). A similar effect was observed for Δπ; however, this measure was particularly susceptible to inflation under the most extreme reductions in DP size, with substantially more elevated values observed between DP size proportions of 0.01 and 0.1 compared with 0.1 and either 0.5 or 1.0. Whilst this pattern was broadly consistent through sampling points, variance among the different DP sizes for Δπ was most pronounced at the initial (100 generations) and final sampling points (10,000 generations). Such observations are unsurprising given that both F_ST_ and Δπ increase with processes that reduce within-population genetic variance. Except for the most extreme reduction in DP size, D_XY_ was generally robust to protracted bottlenecks. Following this, in contrast to F_ST_ and Δπ, D_XY_ decreased when DP sizes were reduced (Figure 2B).

The inclusion of migration reduced absolute values of all measures of divergence. For F_ST_ and D_XY_, variance between migration treatments increased across the simulation, however D_XY_ was generally more consistent across migration treatments. Δπ was also reduced in the presence of migration, although the effect of migration on Δπ was generally consistent across all generations and did not increase through time, as was the case for F_ST_ and D_XY_.

Whilst F_ST_ and D_XY_ increased generally over time, Δπ peaked around generation 500 and declined thereafter. This peak corresponded to the median generation that DP replicates reached their optimum phenotype (median across all data = 519), suggesting this peak reflects selective sweeps. The majority of these patterns were apparent in the Pheno_Null_ (Figure S1) and neutral (Figure S2) simulations, however divergence for all measures was reduced and ultimately negligible by the removal of divergent phenotypes when migration was present.

### Demography moderates the association between measures of divergence and selection

Again, founding bottlenecks had a minimal effect on the correlations observed between strength of selection and measures of divergence with minimal variance observed between bottleneck treatments in all comparisons (Figure 3A).

**Figure 3:**
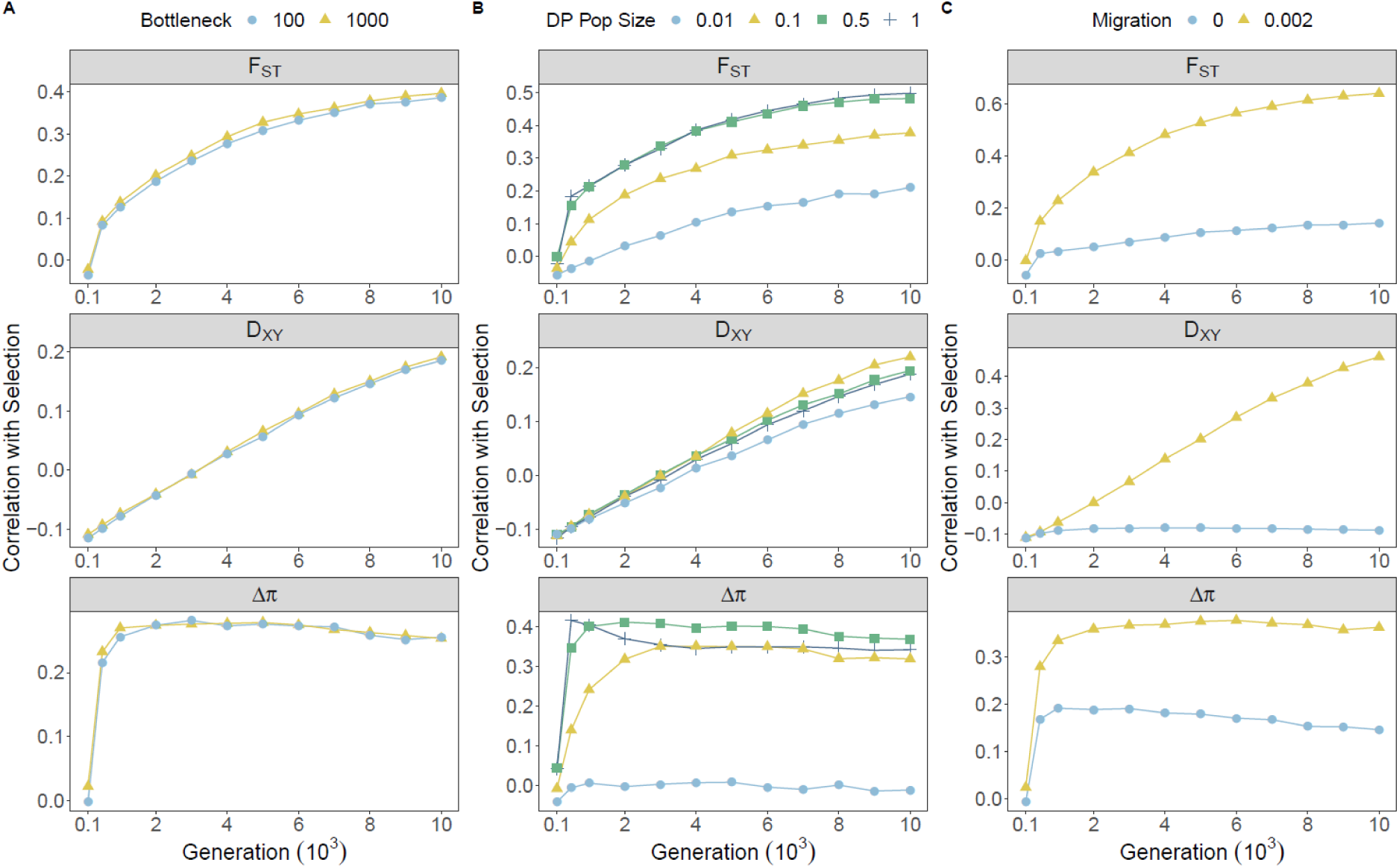
Effects of demographic treatments on the relationship between selection and measures of genetic divergence across all sampling generations. Point colour and shape denote treatment groups for founding bottlenecks (**A**), protracted bottlenecks (**B**) and migration (**C**). Each point represents correlation coefficients calculated across all genes within individual treatments groups, averaged within treatment levels, and averaged over 20 iterations.

Protracted bottlenecks had substantial effects on relative (F_ST_ and Δπ) measures and marginal effects on absolute (D_XY_) measures of divergence (Figure 3B). Relative measures were less correlated with selection when DP size was reduced (0.01 or 0.1) than with minimal (0.5) or no (1.0) reductions in DP size, which were generally similar. For F_ST_, the variance in correlation coefficients between DP size treatments was generally stable across time, whereas variance between treatments for Δπ peaked at 500 generations. For D_XY_, correlations with selection were broadly consistent across DP size reductions up until 10,000 generations, in which associations with selection were reduced by the strongest protracted bottlenecks, but generally robust (Figure 3B).

Expectedly, the absence of migration largely precluded the ability of measures of divergence to predict strength of selection, with notable variance observed between no migration (0.0) and minimal migration (0.002) treatments emerging rapidly for all divergence measures (Figure 3C). Both F_ST_ and D_XY_ variance between migration treatments was greatest at the 10,000 generations sampling point, whereas similar variance was observed between migration treatments for Δπ across simulations. This observation again highlights the significance of temporal differences between measures. Interestingly, correlations between Δπ and selection persisted, albeit weakly, in the absence of migration, which was not the case for F_ST_ and D_XY_.

Protracted bottlenecks had a minimal effect on how measures of divergence correlated with selection in Pheno_Null_ data (Figure S5). Both F_ST_ and D_XY_ became negatively correlated with selection over time without divergent selection, likely due to stronger selection on common alleles shared between AP and DP. Negative correlations were stronger for D_XY_, consistent with reductions in DP_π_ with stronger selection. This suggests associations between selection and D_XY_ in Pheno_Div_ simulations are likely more dependent on adaptive substitutions in order to overcome this effect. Δπ was generally positively associated with selection in Pheno_Null_ simulations regardless of migration, but was slightly reduced when migration was absent.

By examining the correlation coefficients of all 16 unique demographic histories, we can investigate the combined effect of migration and protracted bottlenecks and directly compare effectiveness of individual measures across time (Figure 4). F_ST_ consistently outperforms D_XY_ in terms of associating with selection under most demographic treatments, particularly when DP sizes are larger and migration is present. By 10,000 generations however, the relative dominance of F_ST_ appears to subside, with the trend through time suggesting a relative improvement in D_XY_ in treatments with gene flow (Figure 4). Δπ is similarly more informative than D_XY_ across sampling generations under most demographic treatments. Interestingly at sampling generation 10,000, protracted bottlenecks produce the biggest bias towards Δπ (Figure 4). The resilience of Δπ under no-migration treatments is also apparent in F_ST_ - Δπ comparisons, such that at 3,000 generations Δπ is more informative than F_ST_ in the absence of migration. Consistent with its rapid response to sweeps around 500 generation, Δπ outperforms F_ST_ under most demographic scenarios in early generations. By 10,000 generations, however, F_ST_ performs as well as Δπ without migration and outperforms Δπ with migration.

**Figure 4:**
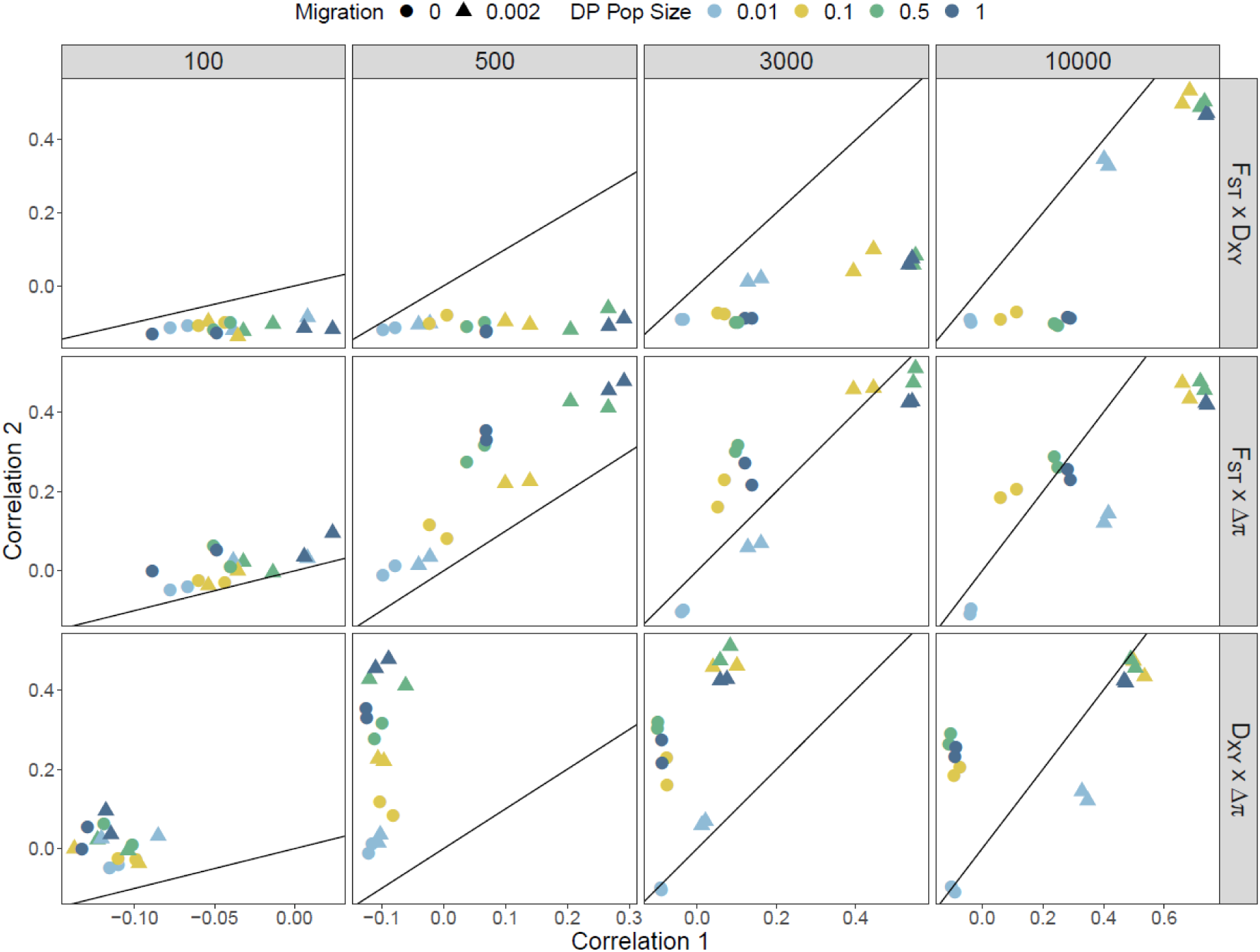
Pairwise comparisons between the correlation coefficients of selection with F_ST_, D_XY_ and Δπ across four sampling points. Each data point represents a unique demographic treatment with points coloured according to DP population size and shaped according to migration level. Correlation 1 refers to the first measure listed in the comparison and Correlation 2 to the second. The y=x line is plotted within each facet to illustrate biases towards one measure. Points below the line are biased towards Correlation 1, whilst points above the line are biased towards Correlation 2. Each point represents correlation coefficients calculated across all genes within individual treatments groups and averaged over 20 iterations.

### Demography moderates the shape and tail end of divergence distributions

By comparing distributions across simulations with divergent (Pheno_Div_) and stabilising (Pheno_Null_) selection with neutral runs, we can examine the effect of demographic treatments on the ability of each measure of divergence to discriminate between them (Figure S11-13; Figure 5). There are few differences between distributions of F_ST_ for the three simulation types when migration is absent between AP and DP replicates, highlighting an increased likelihood of false-positives. The exceptions occur around 500 generations (Figure S12) when sweeps are most common, and towards the end of simulations. Following probable sweeps, Pheno_Div_ F_ST_ is slightly elevated, and towards the end of simulations Pheno_Div_ F_ST_ is marginally lower than Pheno_Null_ F_ST_, which is marginally lower than neutral F_ST_ (Figure 5). With migration, we see few differences between Pheno_Null_ and neutral F_ST_. Pheno_Div_ F_ST_ distributions however become more positive and flattened, according to variable selection, with the majority of Pheno_Div_ F_ST_ above the 95% quantiles of Pheno_Div_ and neutral F_ST_ by 10,000 generations (Figure 5).

**Figure 5:**
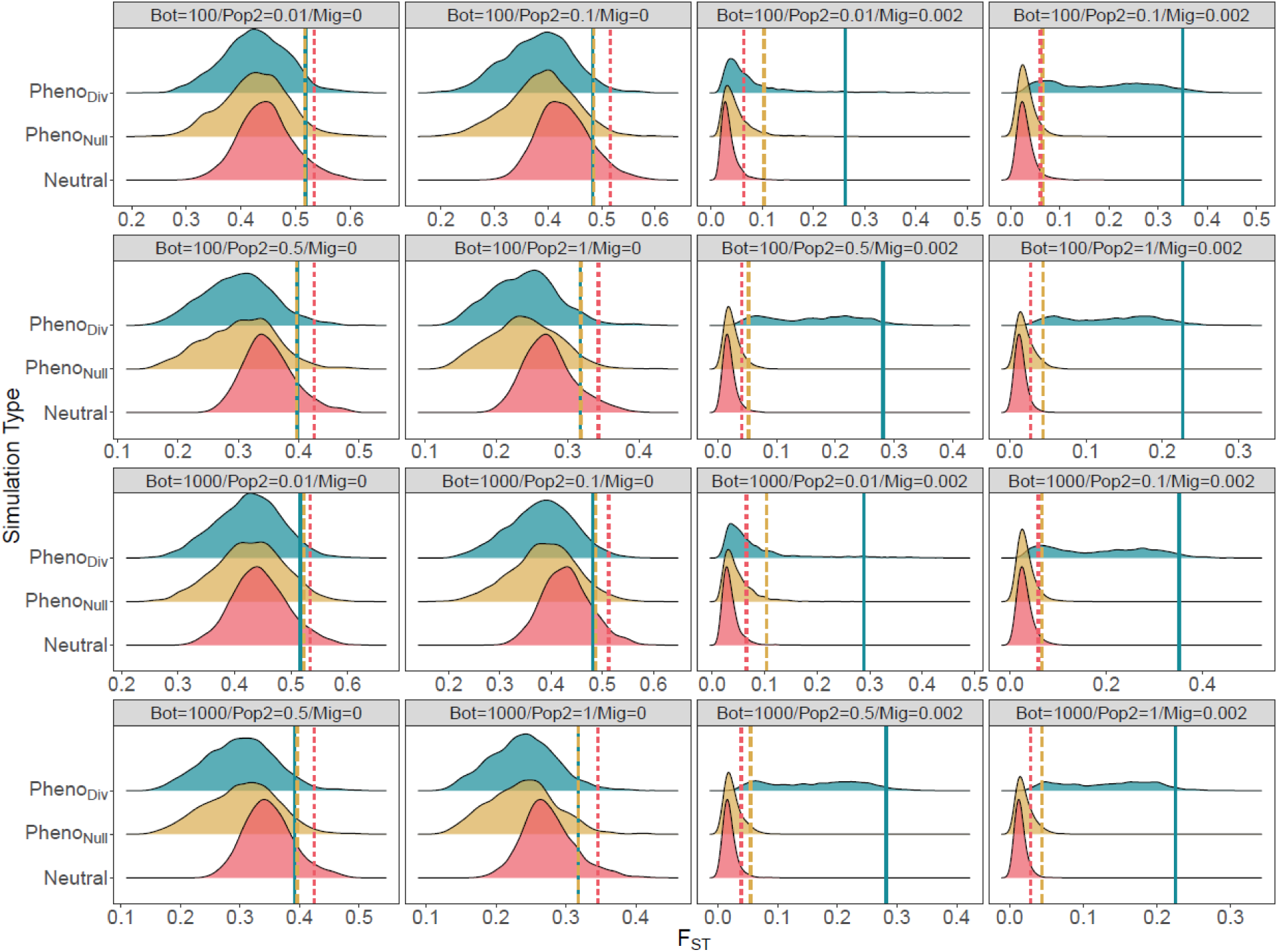
Distributions of F_ST_ under each of the 16 unique demographic treatments under three selection regimes: Pheno_Div_ (divergent selection), Pheno_Null_ (stabilising selection) and Neutral, after 10,000 generations. Upper 5% quantiles are highlighted for each distribution, with linetype corresponding to selection: Solid = Divergent, Dashed = Stabilising, Dotted = Neutral. Each distribution represents data pooled from 20 iterations of 100 gene windows (N = 2000).

Similar patterns were observed for D_XY_ distributions (Figure S14-17), with little to discriminate between in treatments without migration. However, without migration, neutral D_XY_ was generally reduced relative to Pheno_Null_ and Pheno_Div_ D_XY_. With gene flow, like F_ST_, Pheno_Div_ D_XY_ was readily distinguishable from Pheno_Null_ and neutral distributions, but Pheno_Null_ D_XY_ was also generally more positive than neutral D_XY_. These patterns also emerged more slowly than for F_ST_. In contrast, Pheno_Div_ Δπ (Figure S18-21) was elevated according to DP size, such that at 500 generations the majority of windows under divergent selection exhibited Δπ above neutral and Pheno_Null_ 95% cut-offs for all treatments with DP size ≥ 0.5.

We quantified false-positive rates (FPR) by permuting over merged data comprised of randomly sampled 5% Pheno_Div_ windows and 95% neutral windows and observing the upper 5% quantile (Table S1). By 100 generations, F_ST_ FPR ranged between 0.35 and 0.93, and was lower with increased DP size and lower in treatments without migration (Table S1). By 500 generations (Table 2), FPR rates were lower with migration and higher without, but only for treatments with DP sizes of 0.5 and 1.0. F_ST_ FPR remained high (> 0.85) for all treatments with smaller DP sizes. By 3,000 generations, migration was the most important demographic factor for F_ST_ FPR. With migration, FPR ranged from 0.14 – 0.70, and decreased with increasing DP size. Without migration, FPR were high (0.93-0.97), in line with the random proportion of neutral (0.95) data. By the end of simulations, F_ST_ FPR was as low as 0.08, and was no greater than 0.26 with migration and DP size ≥ 0.1. Without migration, FPR was > 0.97.

**Table 1:** Simulation parameters.

| Variable | Value | Description |
| --- | --- | --- |
| AP | 0 | Ancestral phenotypic optimum |
| AP-$S_\sigma$ | 1 | AP Selection ($\sigma$ of fitness around phenotypic optimum, after transforming) |
| AP<sub>N</sub> | 1000 | AP population size |
| DP | 10 | Derived phenotypic optimum |
| DP-$S_\sigma$ | 0.1-10.00 | DP Selection ($\sigma$ of fitness around phenotypic optimum, after transforming) |
| DP<sub>N</sub> | (10, 100, 500, 1000) | DP population size |
| BN<sub>Founding</sub> | (100, 1000) | Founding bottleneck (N burn-in individual genomes sampled to populate DP) |
| $m$ | (0, 0.002) | Migration (Percentage gene flow in both directions) |
| $\mu$ | $4.89e^{-6}$ ($4.89e^{-5}$, $4.89e^{-7}$) | Mutation rate ($bp^{-1}$) (additional mutation rates with 10-fold increase and reduction) |
| $\mu_\sigma$ | 1.00 | Mutation effect size ($\sigma$ of distribution of effect sizes) |
| $r$ | $1.00e^{-8}$ | Recombination rate ($bp^{-1}$) |
| $C_{max}$ | 1.00 | Maximum fitness cost through competition |
| $C_\sigma$ | 0.40 | Local phenotypic competition ($\sigma$ of fitness reductions between local individuals) |
| $C_{Dist}$ | 1.20 | Maximum phenotypic distance between competitive individuals |

**Table 2:**
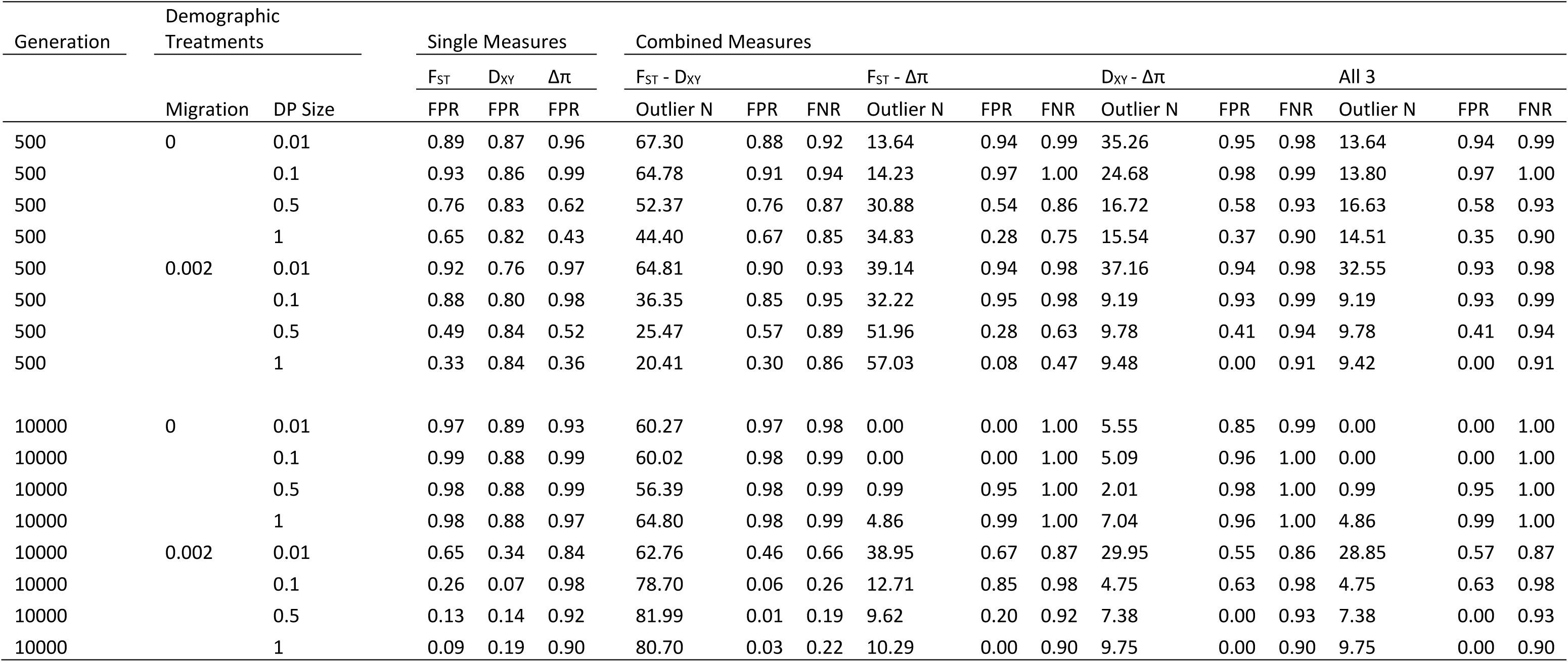
False-positive (FPR) and false-negative rates (FNR) calculated across all measures of divergence and their combined use. Estimates were calculated by combining 5% of data under divergent selection with 95% neutral data and taking upper 5% cut-offs. For single measures, outlier N is always 100 (5% of 2,000) and FNR = FPR. Data represent means calculated over 100 downsampled permutations.

Initial D_XY_ FPR were generally high irrespective of demography, ranging between 0.81 and 0.87. FPR were largely similar after 500 generations, but by 3,000 generations there was a distinction between treatments with (0.33 ≤ FPR ≤ 0.65) and without (0.87 ≤ FPR ≤ 0.90) migration. Interestingly, here FPR rates were lower (0.33 ≤ FPR ≤ 0.46) when DP were smaller (size = 0.01, 0.1) rather than larger (0.57 ≤ FPR ≤ 0.65). This pattern was also observed at the end of simulations, with FPR lower without migration (0.07 ≤ FPR ≤ 0.36) and lowest with DP size = 0.1.

Δπ FPR rates were generally higher across all sampling generations, with FPR not falling below 0.35. There was a clear distinction based on DP size in earlier (100 and 500) sampling points, with FPR lower with larger DP size. However, at later sampling generations (3,000 and 10,000), FPR were generally high (≥ 0.70) regardless of demography.

Taken together, there are clear effects of DP size and migration on distributions of genetic variation and upper quantiles of interest. DP size appears initially most important in the first few hundred generations for F_ST_ and Δπ, but these effects are later swamped by the effect of migration for F_ST_ and are simply eroded for Δπ. D_XY_ exhibits similar patterns to F_ST_, but these develop after many more generations, and whilst gene flow increases the informativeness of D_XY_ for detecting divergent selection, as it does for F_ST_, D_XY_ and F_ST_ experience opposing effects of increases to DP size.

### Demography moderates relationships between measures of divergence

Given F_ST_, D_XY_ and Δπ are all measures of population genetic divergence, there is an assumption that positive correlations should exist between them. We employed the same analysis as above for correlations with selection, but instead examined correlations between individual measures. Founding bottlenecks had minimal effects on the correlations observed between all measures of divergence (Figure 6A).

**Figure 6:**
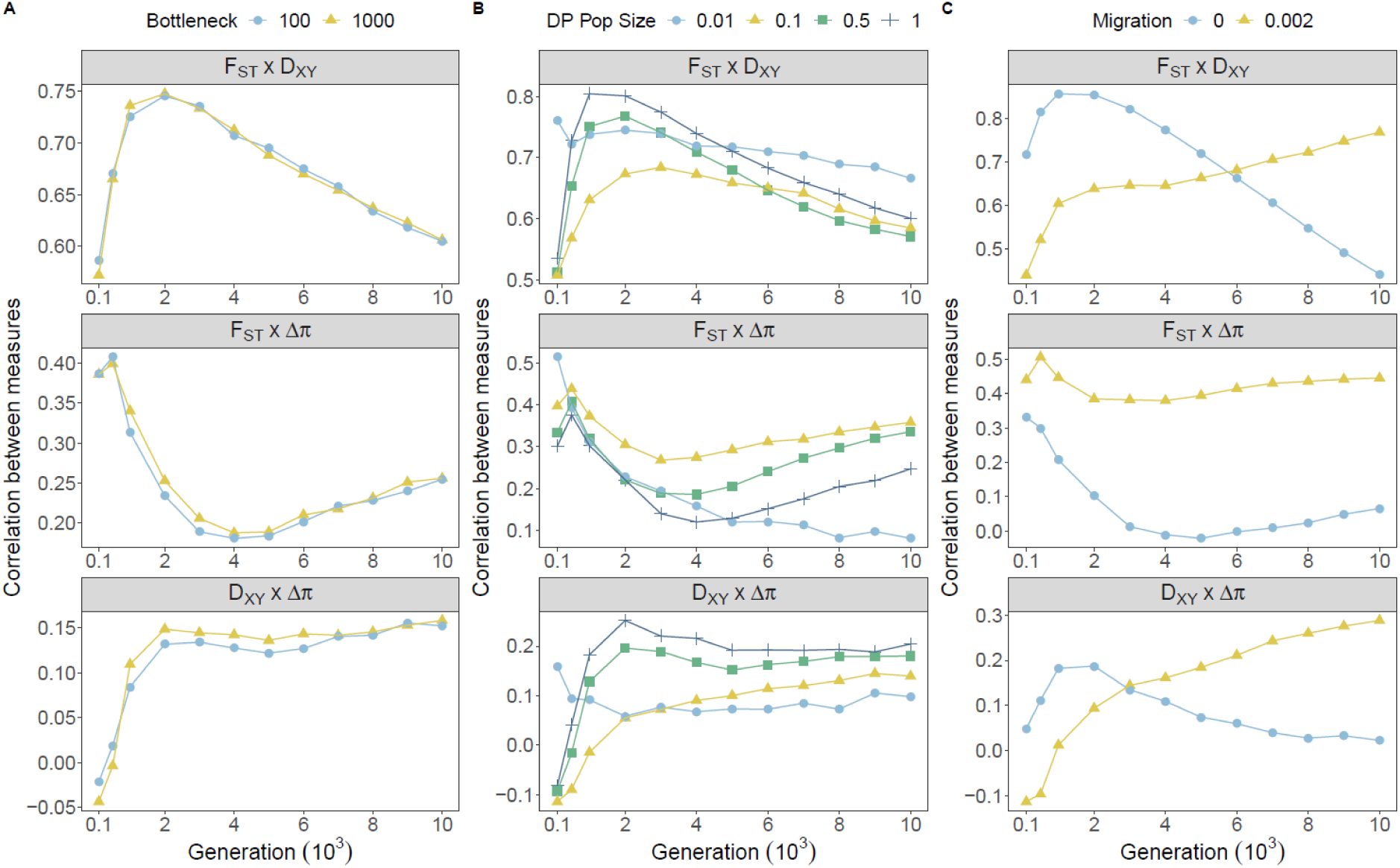
Effects of demographic treatments on the relationship between measures of genetic divergence across all sampling generations. Point colour and shape denote treatment groups for founding bottlenecks (**A**), protracted bottlenecks (**B**) and migration (**C**). Each point represents correlation coefficients calculated across all genes within individual treatments groups, averaged within treatment levels, and averaged over 20 iterations.

Positive correlations between F_ST_ and D_XY_ emerged rapidly irrespective of DP size, but smaller DP sizes generally increased the correlation (Figure 6B), with variance between treatments generally decreasing over time. Similarly, F_ST_ and Δπ were generally positively correlated, however reductions in DP size reduced correlation coefficients. By 4,000 generations, F_ST_ - Δπ correlations for DP sizes ≥ 0.1 stabilised, but correlations under extreme protracted bottlenecks continued to decline to a low of 0.08 (Figure 6B). D_XY_ - Δπ correlations were generally weaker than other comparisons across the course of simulations, but were minimally affected by protracted bottlenecks. Interestingly, correlations with D_XY_ and both F_ST_ and Δπ appear to peak for treatments with minimal population size reductions and without migration at 500-2,000 generations (Figure 6B), suggesting positive correlations with D_XY_ are in part driven by the generation in which sweeps are taking are place.

Migration induced substantial variance between correlations of divergence measures, with effects dependent on sampling point (Figure 6C). In the absence of migration, F_ST_ and D_XY_ were more strongly correlated for the first 6,000 generations than in treatments with gene flow. However, from here until 10,000 generations this pattern reversed and F_ST_ - D_XY_ correlations increased with gene flow and deteriorated in allopatry. F_ST_ - Δπ correlations were strong initially, but a lack of migration weakened the correlation over time until measures were uncorrelated by around 3,000 generations. In contrast, in the presence of migration, positive correlations between F_ST_ and Δπ were relatively stable (R^2^ = 0.51 – 0.38 over the whole simulation period). Interestingly, without migration, D_XY_ and Δπ were largely uncorrelated, but migration induced a positive correlation between D_XY_ and Δπ that emerged after 1,000 generations and continued to increase through time.

Contextualised by our previous demonstrations of associations with selection, these results highlight that selection can induce positive correlations between measures. By 10,000 generations, all pairwise comparisons of divergence measures become positive correlated with migration, which we know is when variation is most strongly associated with selection. Crucially however, this relationship only emerges after several thousand generations, before which we observe positive correlations in migration-absent treatments when associations with selection are weak for all measures (F_ST_ - D_XY_ in particular). Extreme reductions in DP size also increase positive correlations between F_ST_ and D_XY_ despite poor associations with selection in these treatments. These results highlight that positive correlations between statistics are also achievable in the absence of divergent selection. It is also interesting to note the decay of correlations with Δπ and both F_ST_ and D_XY_ in the absence of migration. These observations are most likely driven by the substitution rate. Substitutions that are not linked to selection (drift with ineffective selection) likely drive increased F_ST_ and D_XY_ but not Δπ.

In Pheno_Null_ simulations, F_ST_ and D_XY_ were positively correlated in all treatments, but stronger associations linked with demographic treatments with effective selection did not emerge (Figure S22). This was also true for neutral simulations (Figure S23), but F_ST_ - D_XY_ correlations were larger than for Pheno_Div_ and Pheno_Null_ when selection was ineffective, reaching a maximum of R^2^ = 0.89 without migration (Figure S23C). This demonstrates that the positive correlations in Pheno_Div_ simulations are driven in part by divergently adaptive allele frequency shifts and substitutions when selection is effective, but these correlations also emerge under stabilising selection or neutrality. Specifically, strong positive correlations with Δπ were dependent on divergent selection, and F_ST_ – Δπ correlations became negative over time in treatments without migration under neutrality. D_XY_ – Δπ correlations became negative in Pheno_Null_ data with migration and were unassociated without. Thus, these results highlight that the relationships between measures of divergence are highly dependent on demography, time, and selection experienced over a genomic region.

We then examined FPR and false negative rates (FNR) when combining outliers across F_ST_, D_XY_ and Δπ (Table S1; summaries from 500 and 10,000 generations in Table 2). Combined F_ST_ - D_XY_ outliers exhibited FPR rates that were reasonably high (0.30 - 0.93) and similar to F_ST_ alone at generations 100 and 500, suggesting little improvement in reducing FPR. During this period, FNRs were also high (≥ 0.84), suggesting most windows with divergent selection could not be detected on a neutral backdrop. It was not until 10,000 generations that reductions in FPR were evident for combined F_ST_ - D_XY_ outlier sets for treatments with migration, in some cases dropping to 0.01, although FNR were highly variable (0.19 ≤ FNR ≤ 0.66). High FPR (0.96 ≤ FPR ≤ 0.98) and high FNR (0.98 ≤ FPNR ≤ 0.99) were observed without migration, highlighting most common outliers between F_ST_ and D_XY_ to be neutral, and a failure to detect almost all divergent windows.

Combined F_ST_ - Δπ outliers did marginally outperform F_ST_ and Δπ outliers according to FPR with DP sizes of 0.5 or 1.0 and in earlier (100 and 500) sampling generations. Here, FPR dropped to a low of 0.08, but FNR were reasonable through this time (≥ 0.43). Beyond this (sampling generations 3,000 and 10,000), FPR did drop to 0, however these were generally alongside high FNR of up to 1.0, highlighting a failure to detect any common outliers at all. A good example of improvement over singular measures was observed at generation 500, with no founding bottleneck, equal sized populations, and migration. Here, an average of ∼53% of divergent windows were detected with an average FPR of 0.08. This is compared with: singular F_ST_, where FPR/FNR = 0.33; and singular Δπ where FPR/FNR = 0.36. By 10,000 generations, combined outlier sets of F_ST_ - Δπ performed poorer than singular F_ST_ with migration present, but generally returned low numbers of false positives, unlike F_ST_ - D_XY_.

Combined D_XY_ - Δπ outliers performed poorly across all treatments and all sampling generations, with FNR failing to fall below 0.86. There were, however, initial benefits in terms of reduced FPR in earlier sampling generations for treatments with larger DP sizes (0.5 and 1.0). This discordance between D_XY_ and Δπ and large FNR also limited the combined usage of all three measures together, with similarly high FNR precluding their combined usage.

These results therefore highlight that combining measures can help reduce FPR, but usually at the cost of increased FNR (expectedly), and only under certain demographies. Our observation that high FPR are prevalent among combined outlier sets from statistics, particularly in the absence of migration, suggests their usage must be dependent on a knowledge of the underlying demographic history of populations. These findings also highlight that the strong correlations that emerge between measures of divergence under scenarios with ineffective selection or even under neutrality do extend to the tail-ends of distributions.

### Demography drives signals of convergence irrespective of selection

Clusters of overlapping outliers developed steadily over time (Figure S26-28). By sampling generation 10,000, significant proportions of overlapping outliers were recovered across different demographic treatment groups (Figure 7). For F_ST_ and D_XY_, clustering of treatments was driven by the presence/absence of migration, with the highest proportions of convergent outliers observed between treatments with migration. Interestingly, for both measures, reasonable proportions of convergent outliers were recovered across no migration treatments, which we have demonstrated to have ineffective selection. Importantly, there was little overlap between these clusters, suggesting different convergent outliers within each cluster. Combining outliers from F_ST_ and D_XY_ did not alter the patterns observed in either, and importantly did not remove convergent outliers across no-migration demographies. There were minimal convergent outliers observed for Δπ outliers, however combining Δπ with F_ST_ and D_XY_ did appear somewhat effective in removing convergent outliers found between no-migration treatments. However, for both combination of Δπ with F_ST_ and D_XY_, the highest proportional overlap was observed for treatments with the smallest DP size.

**Figure 7:**
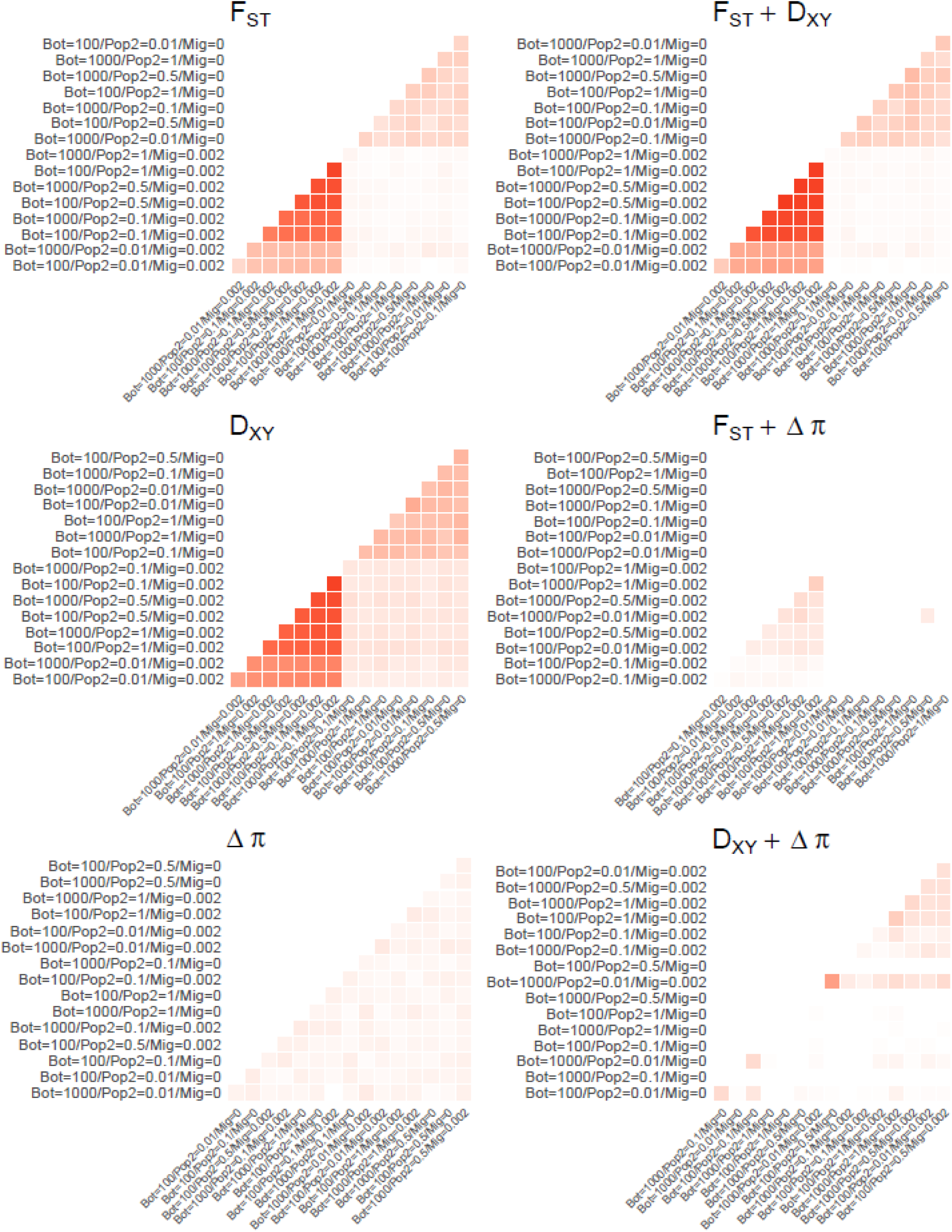
The proportional overlap of outliers above the 95% quantile, averaged across 100 downsampled datasets consisting of 95% Neutral and 5% Pheno_Div_ data for each of the 16 demographic treatments after 10,000 generations. Axis orderings were determined through hierarchical clustering. Heatmaps are shown for single measures of F_ST_, D_XY_ and Δπ in the first column, and combined measures in the second column. Heatmaps are coloured according to a common scale of 0 to 1. Treatments are labelled with founding bottleneck (Bot), DP population size (Pop2), and migration (Mig) values.

Interestingly, the clustering of F_ST_ - D_XY_ outliers in the presence of migration was greatly reduced when divergent selection was removed in both Pheno_Null_ (Figure S29-32) and neutral data (Figure S33-36), but clusters of outliers in the absence of migration were still apparent.

Because migration was the dominant factor in clustering of treatments with convergent outlier overlap, we sought to investigate what features of simulated genes drove variance in measures of divergence with treatments separated by migration factor using linear mixed models at the final sampling generation. With migration between AP and DP, selection had by far the strongest effect on F_ST_ (Table 3) in the expected positive direction. We also observed a weaker positive association with selection target (% coding) of gene (Table 3), and weaker negative associations with Pheno Gen (generation DP optimum reached) (Table 3). These fixed effects explained 39.6% of variance in F_ST_ with migration. Conversely, F_ST_ in treatments without migration was strongly positively associated with selection target % (Table 3), with a much weaker positive association with selection (Table 3). However, fixed effects here explained only 6.6% variance in F_ST_ without migration.

**Table 3:** LMM results for models of measures of divergence explained by features of simulated genes. For all models, random variables included gene ID and demographic treatment. Fixed effects with the largest effect are highlighted in bold. Selection Target and Pheno Gen were log_10_-transformed in models.

| Measure | Migration | Var. explained (%) |  | Fixed Effect | Estimate | Std. Error | df | T | P |
| --- | --- | --- | --- | --- | --- | --- | --- | --- | --- |
|  |  | Fixed | Random |  |  |  |  |  |  |
| F<sub>ST</sub> | 0.002 | 39.60 | 60.90 | **Selection** | **0.100** | **0.004** | **98** | **25.48** | **<2.00E-16** |
|  |  |  |  | Selection Target | 0.009 | 0.002 | 103 | 3.89 | 0.000177 |
|  |  |  |  | Pheno Gen | -0.007 | 0.001 | 14530 | -9.45 | <2.00E-16 |
| F<sub>ST</sub> | 0 | 6.60 | 75.40 | **Selection Target** | **0.025** | **0.002** | **97** | **11.44** | **<2.00E-16** |
|  |  |  |  | Selection | 0.011 | 0.004 | 97 | 3.04 | 0.00302 |
| D<sub>XY</sub> | 0.002 | 18.60 | 69.40 | **Selection** | **0.024** | **0.001** | **97** | **16.42** | **<2.00E-16** |
|  |  |  |  | Selection Target | -0.010 | 0.001 | 102 | -11.70 | <2.00E-16 |
|  |  |  |  | Pheno Gen | -0.003 | 0.000 | 14570 | -14.70 | <2.00E-16 |
| D<sub>XY</sub> | 0 | 13.50 | 33.30 | **Selection Target** | **-0.008** | **0.001** | **99** | **-8.00** | **2.39E-12** |
|  |  |  |  | Pheno Gen | 0.000 | 0.000 | 14480 | 2.37 | 0.018 |
| Δπ | 0.002 | 5.00 | 60.90 | **Selection** | **0.168** | **0.011** | **99** | **15.67** | **<2.00E-16** |
| Δπ | 0 | 0.70 | 76.70 | **Selection** | **0.086** | **0.018** | **99** | **4.73** | **7.63E-06** |
|  |  |  |  | Pheno Gen | -0.009 | 0.003 | 12870 | -2.70 | 0.00694 |

D_XY_ was similarly most strongly positively associated with selection in treatments with migration (Table 3), but the strength of this effect relative to the other model effects was not as large as observed for F_ST_. D_XY_ was also negatively associated with Pheno Gen (Table 3), as was F_ST_, but negatively associated with selection target % (Table 3), contrary to F_ST_. In treatments without migration, selection target % was strongly negatively associated with D_XY_ (Table 3), with a weak positive effect of Pheno Gen (Table 3).

Selection was the most important fixed effect for Δπ regardless of migration (Table 3), however both models explained minimal variance (5% with migration, 0.7% without migration). This is consistent with the previous demonstration of the erosion of Δπ over time (Figure 1).

Removing divergent selection (i.e. in Pheno_Null_ simulations) modified models for F_ST_ and D_XY_ for treatments with migration. Selection and selection target remained the most important model effects, but had similar sized effects, and ultimately variance explained dropped from 39.6% to 5.5%. Conversely, the model for Pheno_Null_ D_XY_ with migration increased variance explained from 18.6% to 32.9% compared with Pheno_Div_ D_XY_. Selection target had a greater effect in this model than strength of selection. Δπ models were unchanged. Together, these results confirm that divergent selection, and not stabilising selection within DP, drive F_ST_ and D_XY_ variation in Pheno_Div_ simulations.

## DISCUSSION

### Summary of results

Here, we show that different demographies can have dramatic effects on how measures of population divergence identify regions of the genome under selection. Effects are also strongly time-dependent. Using simulated populations, we have demonstrated the relative influences of founding bottlenecks, protracted bottlenecks, and migration on three commonly used measures of genetic divergence (F_ST_, D_XY_, Δπ), whilst demonstrating the relative usefulness of each measure for informing on selection under different demographic conditions.

We find that founding bottlenecks have little effect on population divergence measures, potentially either because populations quickly recover (before first sampling after 100 generations), or founding bottlenecks of 10% (100 individuals) were not extreme enough. Protracted bottlenecks (reductions in DP size) however, artificially inflate F_ST_ and Δπ but reduce D_XY_, and can erase the relationship of F_ST_ and D_XY_ with selection under the most extreme reductions in population size. Relative measures are, in part, driven by intra-population changes in allele frequency, which become exaggerated in smaller populations as a product of drift (B. Charlesworth 2009; Ellegren and Galtier 2016). As a consequence, we observe inflated measures of relative divergence as allele frequencies drift in smaller DP replicates.

In contrast, D_XY_ increases with larger DP size, as a product of the relationship between the number of segregating sites and the population-level mutation rate (4N_e_μ) (Hartl, Clark, and Clark 1997). D_XY_ is a measure of sequence divergence and is averaged across all sites (although similar statistics limit averaging to segregating sites only), which results in higher D_XY_ as segregating sites are introduced into either population at a rate of 4N_e_μ. This relationship with the number of segregating sites can be observed by examining the positive relationships between D_XY_ and DP_π_ (Figure S37). Overall, the relationship between D_XY_ and selection is less affected by protracted bottlenecks than F_ST_ and Δπ, likely due to the lack of allele frequency relevance. Consider for example, two SNPs with minor allele frequencies of 0/0.5 (SNP 1) and 0.5/0.5 (SNP 2) in AP/DP. Each locus contributes equally towards D_xy_ (SNP 1 = [0 x 0.5] + [1 x 0.5] = 0.5; SNP 2 = [0.5 x 0.5] + [0.5 x 0.5] = 0.5), whereas the reduction of within-population variance observed for SNP 1 inflates F_ST_ and Δπ. However, we also see evidence of D_XY_ FPR increasing with increased DP size (Table 2), suggesting this relationship with increased acquisition of segregating sites may conflict with increased efficacy of selection.

Migration, even at the relatively modest rate of 0.2% employed here, substantially reduced absolute values for all measures of divergence. However, whilst absolute values were reduced, their informativeness of selection coefficients increased dramatically (both in terms of their overall relationship with selection and in identifying outliers). Such a result is expected given the role of gene-flow in homogenising neutral loci (reducing measures of divergence), whilst retaining population divergence around adaptive loci (increasing informativeness). The well-known ‘genomic islands of divergence’ model is often invoked to explain this pattern of heterogenous genomic divergence in studies of speciation-with-gene-flow (Nosil, Funk, and Ortiz-Barrientos 2009; Turner, Hahn, and Nuzhdin 2005).

Migration also exhibited interesting temporal patterns that are useful for discussing the discrepancies observed between Δπ and both F_ST_ and D_XY_. Of the measures considered here, Δπ uniquely disregards SNP substitutions in its calculation, as fixed substitutions have no influence on intra-population heterozygosity beyond reducing the heterozygosity of proximate SNPs in the aftermath of a hard sweep. This characteristic explains why Δπ responds to selection more rapidly than F_ST_ and D_xy_ and is influenced by migration in a constant manner across time. In addition, the inability of Δπ to account for adaptive substitutions is likely why the informativeness of Δπ peaked at around 500 generations (approximate median for sweeps), decayed, and then stabilised by 10,000 generations. In contrast, the contributions of adaptive substitutions to F_ST_ and D_xy_ over time increases their predictive power. This characteristic of Δπ, however, also makes it unable to differentiate between divergent and stabilising selection. This temporal observation has important implications for studies of rapid adaptation on the scale of 10s to 100s of generations, in which our simulations suggest Δπ may be the most informative measure of divergence, whilst D_xy_ is initially uninformative for several thousand generations.

Examining the effects of demographic variation on the associations between measures of divergence themselves is a novel element to this study. The relative weaknesses of individual measures of divergences has prompted a recent movement within the literature to employ multiple measures of divergence to avoid false-positives (for e.g. Tine et al. 2014; Malinsky et al. 2015; Hämälä and Savolainen 2019). Our results provide some support for this strategy, with reductions in FPR in treatments with gene flow generally, and low FPR when combining either F_ST_ or D_XY_ with Δπ (albeit at a reasonably high FNR). However, we also find large number of overlapping outliers across combined F_ST_ - D_XY_ with a large FPR across treatments without migration. Further, these false-positives persisted in Pheno_Null_ and neutral data, whereas clusters of treatments with overlapping outliers (Figure 7) and with low FPR were restricted to simulations with divergent phenotypes. It is clear then that combining outlier sets of F_ST_ and D_XY_ only improves analyses under certain demographic histories.

Cruickshank and Hahn (2014) suggested that a disagreement between F_ST_ and D_XY_ outliers in genomic-islands-of-divergence tests highlights a particular susceptibility of F_ST_ to BGS (although see (Matthey-Doret and Whitlock 2019)). Our results are in line with the notion that D_XY_ may be more resistant to false positives due to BGS, given that reductions in DP size resulted in minimal variance in D_XY_ (assuming reductions in population size are analogous to different rates of BGS across the genome exhibit reduced N_e_). However, BGS was not specifically manipulated in these simulations, and results are difficult to disentangle with variation in efficacy of removing deleterious mutations. Further, this minimal variance may be explained by the opposing forces of selection efficacy and acquisition of segregating sites discussed previously. We also found that a greater proportion of variance could be explained by the size of selection target for D_XY_ in the absence of migration than for F_ST_. We found that F_ST_ and D_XY_ are positively correlated under several demographic scenarios, but this strong association only reflects selection when gene-flow is present and only after several thousand generations, consistent with previous simulation work (Ravinet et al. 2017). When migration is absent, and selection ineffective, F_ST_ and D_XY_ are also positively correlated; but this relationship decays over time (Figure 6C), with empirical evidence (see below) suggesting a negative relationship is likely to emerge without gene flow. No-migration treatments are consistent with Isolation-By-Distance demographic histories and, similarly, comparisons between reproductively-isolated populations or species. Indeed, negative correlations between F_ST_ and D_XY_ within and between clades of birds (Vijay et al. 2017; Irwin et al. 2016), within a radiation event of monkeyflowers (Stankowski et al. 2018) and between speciating orca populations (Foote et al. 2016) support a declining relationship between these measures over long periods of time in isolation. In agreement, we find that without migration F_ST_ and D_XY_ exhibit opposite associations with the proportion of windows made up of coding elements. Mechanistically, a larger selection target increases the rate of deleterious mutation, reducing local π directly through loss of polymorphic deleterious sites, or indirectly through the loss of linked neutral variants under a BGS model. Reductions in local π increase F_ST_ and decrease D_XY_. Over time, associations between F_ST_ and D_XY_ should stabilise, given π is generally well-conserved in stable populations even across long time periods (Romiguier et al. 2014; Van Doren et al. 2017; Dutoit et al. 2017).

By comparing our results to a second dataset that lacked divergent selection (Pheno_Null_), we found consistent support that positive associations between D_XY_ and selection are strongly dependent on the inclusion of a divergent phenotype. Positive associations with F_ST_ are attainable only in the absence of migration, and are weakly negative without, and Δπ patterns are primarily driven by variable stabilising selection and made weaker by the inclusion of phenotypic divergence. These comparisons are useful in highlighting the relative roles of adaptive allele frequency changes and substitutions in driving patterns of genetic differentiation. It is also of interest to note that overlapping outliers are readily attainable across no-migration treatments in Pheno_Null_ (Figure S32) and neutral simulations (Figure S36), but not for treatments with migration. This confirms that overlapping outliers in no-migration treatments occur due to common neutral processes within genomic windows, whereas overlapping outliers in treatments with migration are driven by the effects of divergent selection. The discrepancies between our Pheno_Div_ and other simulated datasets highlight the necessity in quantifying phenotypic differences or environmental selection pressure when interpreting patterns of variation across the genome.

### Detecting genetic convergence

In addition to understanding how outlier detection in individual pairs were affected by demography, we wanted to explore how studies looking at overlapping outliers in multiple pairs (i.e. detecting genetic convergence) were affected by demography. Our simulation design, in which an ancestral burn-in population is used to found 16 independent AP-DP pairs is analogous to replicated ecotype population pairs in model systems, such as various ecotype pairs of the three-spined stickleback, *Gasterosteus aculeatus* (Hohenlohe et al. 2010; Jones, Chan, et al. 2012); high/low predation Trinidadian guppies, *Poecilia reticulata* (Fraser et al. 2015); crab/wave ecotype periwinkles, *Littorina saxitilis* (Ravinet et al. 2016; Kess, Galindo, and Boulding 2018; Westram et al. 2014), and alpine/montane ecotypes of *Heliosperma pusillum* (Trucchi et al. 2017). We found overlapping outliers between demographic treatments and thus, that signals of convergent outliers are attainable for singularly used F_ST_ and D_XY_, and combined outlier sets for F_ST_ – Δπ, D_XY_ – Δπ and F_ST_ - D_XY_. Clustering of outliers was predominantly driven by the presence or absence of migration (with minimal overlap between clusters) (Figure 7).

The overlap of quantile-based outliers between demographic treatments without migration and ineffective selection may be explained in part probabilistically. With migration restricted, AP and DP exhibit significantly elevated measures of divergence, as seen in Figure 1. This increase results in a normal distribution of F_ST_ and D_XY_ across neutral simulations without migration. In contrast, with migration, drift is limited and random recombination influences gene flow and subsequently variation in divergence. Distributions of divergence with migration under neutrality are therefore are heavily right tail-skewed (Figure S38). Spread of data increases with longer right tails, and density of data in each overlapping distribution is limited to the median and lower quantiles.

We also expect some effect of different amounts of starting variation among genome windows following burn-ins. With gene flow restricted, this variation may promote overlap under neutral conditions given all demographic treatments are founded from a common burn-in. However, this feature of our simulations is analogous to the conserved landscapes of variation observed in natural genomes (Burri 2017; Vijay et al. 2017; Stankowski et al. 2019).

Empirically, the influence of migration on outlier overlap has been observed in replicate pairs of parasitic and non-parasitic lampreys. As we show here, when comparing outliers from disconnected and connected parasitic/non-parasitic pairs, Rougemont et al. (2017) recover greater numbers of overlapping outliers among comparisons of disconnected pairs than connected pairs. Overlapping outliers among connected pairs are however better correlated, which the authors suggest reflects selection.

### Limitations of simulations

In our analyses, we grouped genomic windows within 16 unique demographic treatments, assuming the effects of reductions in population size and migration are equivalent for all windows. This may be unrepresentative of genomes sampled from the wild, in which gene flow and effective population size can vary across genomic regions through structural variation, variable recombination rate and BGS (Gossmann, Woolfit, and Eyre-Walker 2011). A prominent example of such a mechanism is the well-characterised consequence of reduced gene flow within inversions that carry locally adapted alleles. Assuming inversion variants are fixed between populations, gene flow across the locus is limited by the prevention of recombinant haplotypes and resistance to introgression (Kirkpatrick and Barton 2006; Ravinet et al. 2017). Recent genome scan studies have highlighted convergent outliers within inversions (Jones, Grabherr, et al. 2012; Nishikawa et al. 2015; Morales et al. 2018), confirming theoretical models regarding their role in shielding adaptive haplotypes from introgression during the adaptation process. Our simulations suggest that genes with reduced migration, and reduced N_e_ as a consequence, will have inflated measures of divergence and differentiation relative to other genomic regions. Therefore, we predict this may have led to their over-representation in genome scan outliers, and increased potential to overlap across replicate populations. Thus, caution should be taken regarding the adaptive significance of these outliers relative to absolutely lower values of genetic divergence attained from regions outside of chromosomal rearrangements.

By choosing to use a factorial design here, we have increased our understanding of the interplay between individual features of demographic history and multiple measures of population divergence. However, computational limitations constrained absolute population sizes to a maximum of 1,000 individuals and generations to a maximum of 10,000 (with 10,000 generation burn-in). To mitigate these constraints, we repeated the analysis over multiple mutation rates to illustrate patterns over 100-fold variation of θ. In general, most trends were consistent, suggesting that results should be consistent across taxa of variable effective population size. However, certain patterns were exaggerated or dampened with increased or decreased mutation rate respectively. For instance, it is clear that the time lag on informativeness of D_XY_ on selection is shortest when θ is largest, which results in stronger positive associations between F_ST_ and D_XY_ by generation 10,000 (Figure S24). It is well-documented that both N_e_ (Frankham 1995) and mutation rate (μ) (Hodgkinson and Eyre-Walker 2011) are highly variable across taxa, which suggests that applying knowledge of the relationships between measures of population differentiation will vary in nature.

Furthermore, temporal variation within our simulations is confounded by the time at which there was a major shift in the DP phenotype (Figure S39), and by extension when selective sweeps occur. This variation in timing is random with respect to adaptive mutations arising *de novo*, but is also influenced by demography. For example, treatments that experience founding bottlenecks are less likely to evolve using variation in the founder, increasing dependence on *de novo* mutations for adaptation. Additionally, variation in our DP size parameter modifies the per population number of new mutations per generation. This is also true for features of simulated genes, such as size of selection target. Predictable temporal variation in the time at which adaptation is likely to occur is a probable source of variance between measures of divergence and is particularly clear for Δπ, but this was controlled for in later modelling analyses as a fixed effect.

A further consideration for these simulations concerns the architecture of the phenotype. Results reported here pertain to mutation effect sizes drawn from a distribution centred at 0 with σ = 1. This produces mutations of typically large effect, but was selected on the basis of phenotypic optima being reasonably distant, with a difference of 10. Thus, 99% of mutations in simulations produce phenotypic differences of less than a third the divergence distance of AP and DP phenotypes. There are numerous factors that influence the distribution of effect sizes in nature, including; selection, mutation, drift, gene flow, extent of pleiotropy and distance to phenotypic optima (Dittmar et al. 2016), with no single expectation for natural systems as a result. The relatively large distance between optima in our simulations, as well as the rapid change in optima implemented in simulations (Collins and De Meaux 2009), likely gives increased importance to mutations of large effect. The interactions between mutation effect size and the results presented here are beyond the scope of the current study, but we did investigate the effect of reducing σ of mutation effect distributions to 0.1 (Supporting Information; Figures S40-45). Briefly, we see reductions in the strength of correlations and associations with selection with decreasing phenotypic effects of mutation, consistent with the notion of softer sweeps on small-effect loci. We also see increased variance in the amount of time taken for simulations to reach the Pheno_Div_ optimum, which agrees with the probable importance of large effect loci in our standard dataset. We, however, retain strong positive associations between measures such as F_ST_ and D_XY_, as we see in neutral simulations, as well as overlapping outliers linked to selection at the tail ends of Pheno_Div_ distributions. Running the simulations in this way suggests that many of the patterns described here may be robust to scenarios with reduced mutation effect sizes.

It is also important when translating these results to genomic data to consider how correlations between measures of divergence depend on the selection type used in simulations. For example, F_ST_ and D_XY_ are strongly positively correlated under neutrality without migration, but under the same demographic scenarios we observe a decay in the relationship between F_ST_ and D_XY_ when divergent selection is involved. Genomes of natural populations will include regions that are neutral or nearly neutral, under stabilising selection around a common phenotypic optimum, or divergent between populations. Thus, the patterns described here may not apply to all genomic windows pooled together.

### Concluding remarks

We have used forward-in-time simulations to perform a factorial experiment in which we explored the relationships between three measures of genetic divergence, selection, and features of demographic history that are commonly variable in natural populations. In agreement with previous theoretical work, we found the reliance of measures of genetic divergence to indicate regions under selection are dependent on demography and variable through time, with a notable lag in D_XY_. Furthermore, we provided novel comparisons between measures of genetic divergence that call into question the use of multiple measures to rule out false-positives. We also demonstrated that signals of convergent evolution across independent replicates can be driven by similar demographic histories with minimal influences of selection. Therefore, we strongly advise those using overlapping outlier scans to carefully consider the demographic context of their system to avoid false-positives. In particular, the presence or absence of migration between diverging populations is a key factor determining the informativeness of genetic variation for selection, and importantly shapes our expectations of outlier overlap among replicate population pairs. It is tempting to assume that replication in study design or analysis in the form of taking multiple measures of genetic divergence can reduce the risk of attaining false-positives. We hope to emphasise in this study that this is not always the case, as false-positives (i.e. genome scan outliers that are not associated with regions under divergent selection) can be driven by non-random genomic or common demographic features that cannot be bypassed through replication. Moreover, many of the patterns we observe are variable through time, such that the relevant pitfalls of analyses will depend on the age of the populations being considered. It should thus also be important to estimate population splits, as a young replicate pair and older replicate pair with similar demographic histories should be expected to exhibit potentially different patterns of genetic variation.

Recent simulation work by Quilodrán et al. (2019), has also emphasised the influence of genomic features and demography, and includes simulation software for estimating the distribution of genetic variation over user-defined chromosomes. Such an approach is particularly useful for systems with chromosome-level genome assemblies in order to gain a sense for how features such as recombination, gene density, and selection targets may produce false-positives under certain demographies. The software employed here, SLiM (Haller and Messer 2019), may also be used to this end, and the scripts accompanying this study will facilitate similar analyses over system-specific genome regions. Further, recent work on genomic landscapes of linked selection (Stankowski et al. 2019) has highlighted that much of the total variance of genetic divergence such as F_ST_, D_XY_, and π can be explained by the major principal component (PC1) over numerous pairwise comparisons. These population comparisons need not reflect divergent phenotypes, as PC1 reflects genomic features associated with diversity landscapes. Adopting this approach may be useful for systems lacking a chromosome-level genome assembly by estimating SNPs or regions with non-random elevated measures of divergence associated with genome features. Such SNPs or regions may be particularly prone to false-positives under certain demographic histories.

## ACKNOWLEDGEMENTS

The authors wish to thank all members of the Fraser lab for insightful discussions on this project, as well as members of the University of Exeter’s EB theme and the University of Sussex’s EBE group. The authors would also like to thank Kai Zeng and Adam Eyre-Walker for feedback on earlier copies of this manuscript, and two anonymous reviewers for inciteful and helpful comments.

## AUTHOR CONTRIBUTIONS

JRW and BAF conceived the project, developed the experimental design and contributed towards the writing of the manuscript. JRW wrote and implemented the pipeline for simulations and analysis.

## DATA ACCESSIBILITY

A full pipeline including all bash, Eidos and R scripts necessary to repeat this analysis can be downloaded from Github (https://github.com/JimWhiting91/Contingent_Convergence_Pipeline). Supplementary figures (S1-48) have been uploaded through the GSA figshare portal. Supplementary figures include results for different mutation rates, Pheno_Null_ and neutral data across sampling generations along with additional figures.

