## Supporting Figures for "Contingent Convergence: The ability to detect convergent genomic evolution is dependent on population size and migration"

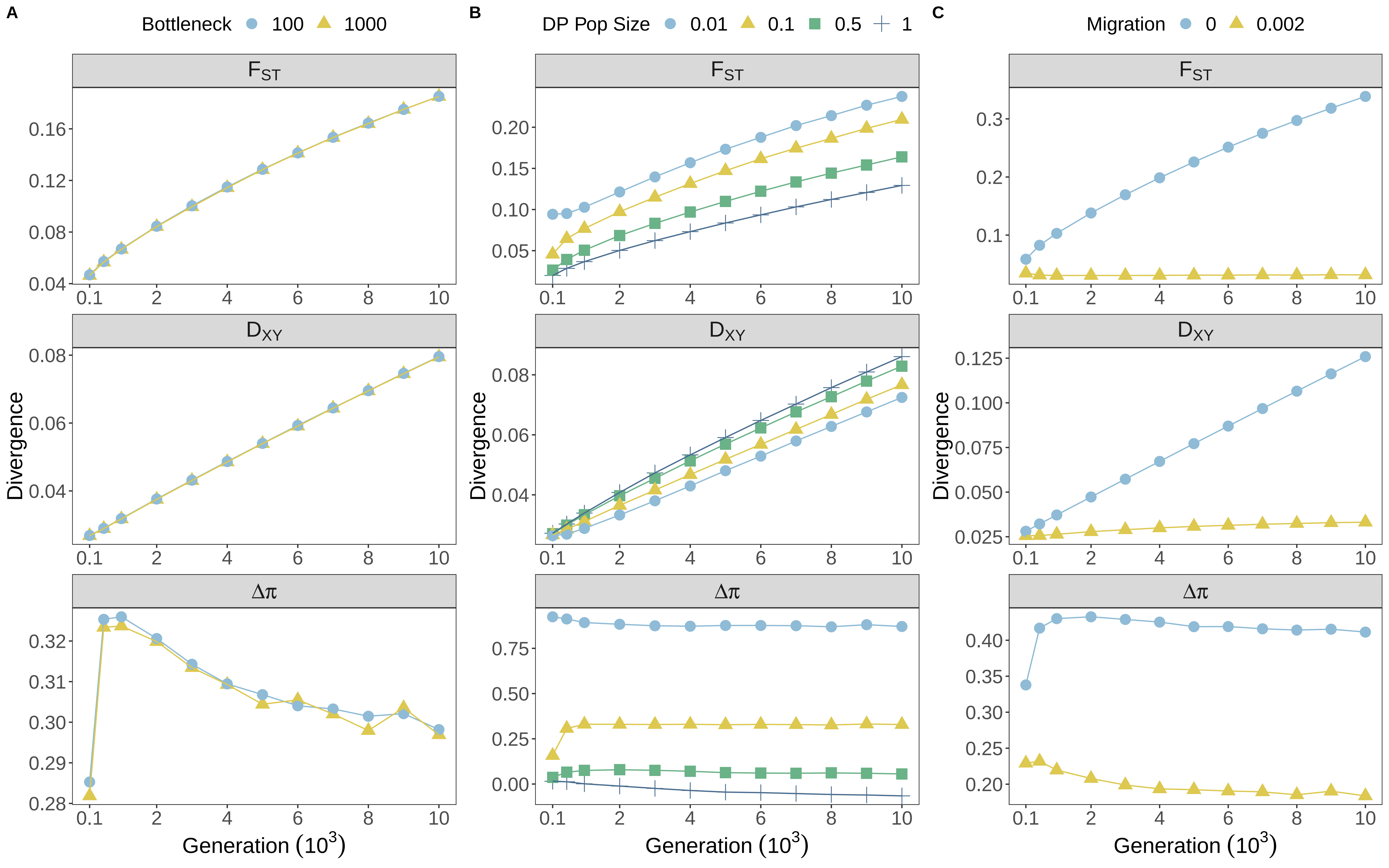

A
Bottleneck 100 1000
B
DP Pop Size 0.01 0.1 0.5 1
C
Migration 0 0.002
$F_{ST}$
$D_{XY}$
$\Delta\pi$
Divergence
Generation ($10^3$)

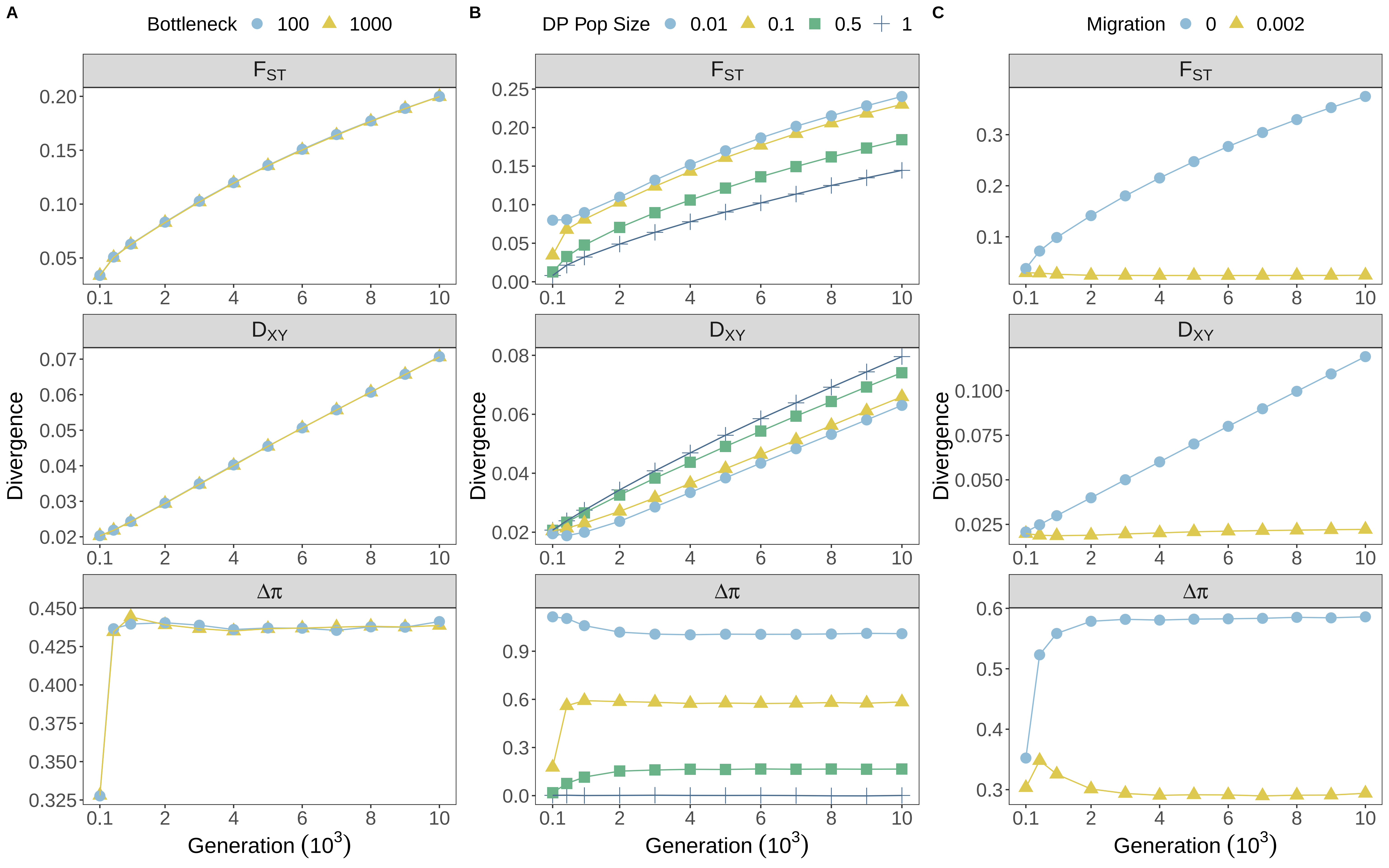

A
Bottleneck 100 1000
B
DP Pop Size 0.01 0.1 0.5 1
C
Migration 0 0.002
F_ST
D_XY
Δπ
Divergence
Generation (10^3)

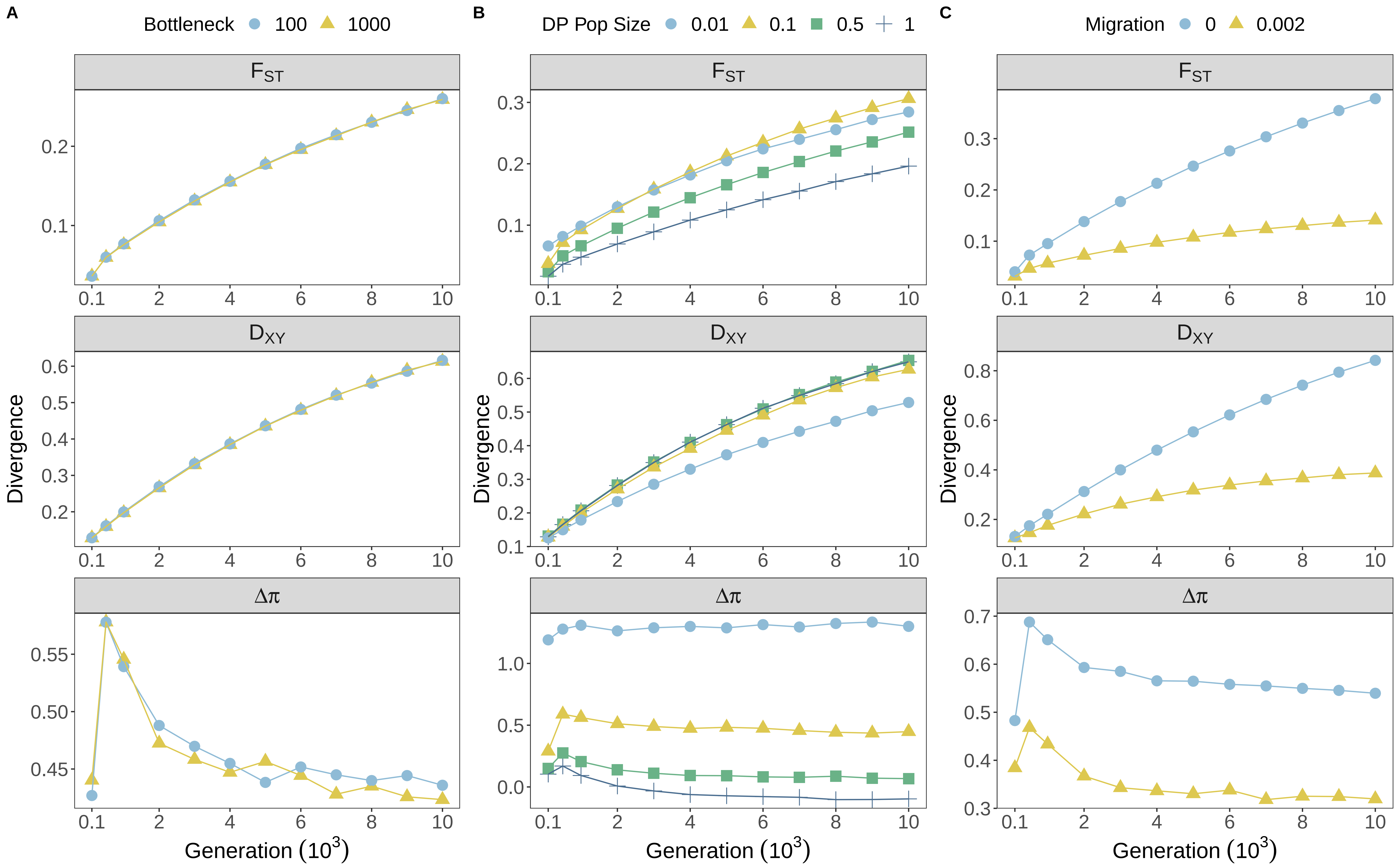
A
Bottleneck 100 1000
B
DP Pop Size 0.01 0.1 0.5 1
C
Migration 0 0.002
$F_{ST}$
$D_{XY}$
$\Delta\pi$
Divergence
Generation ($10^3$)

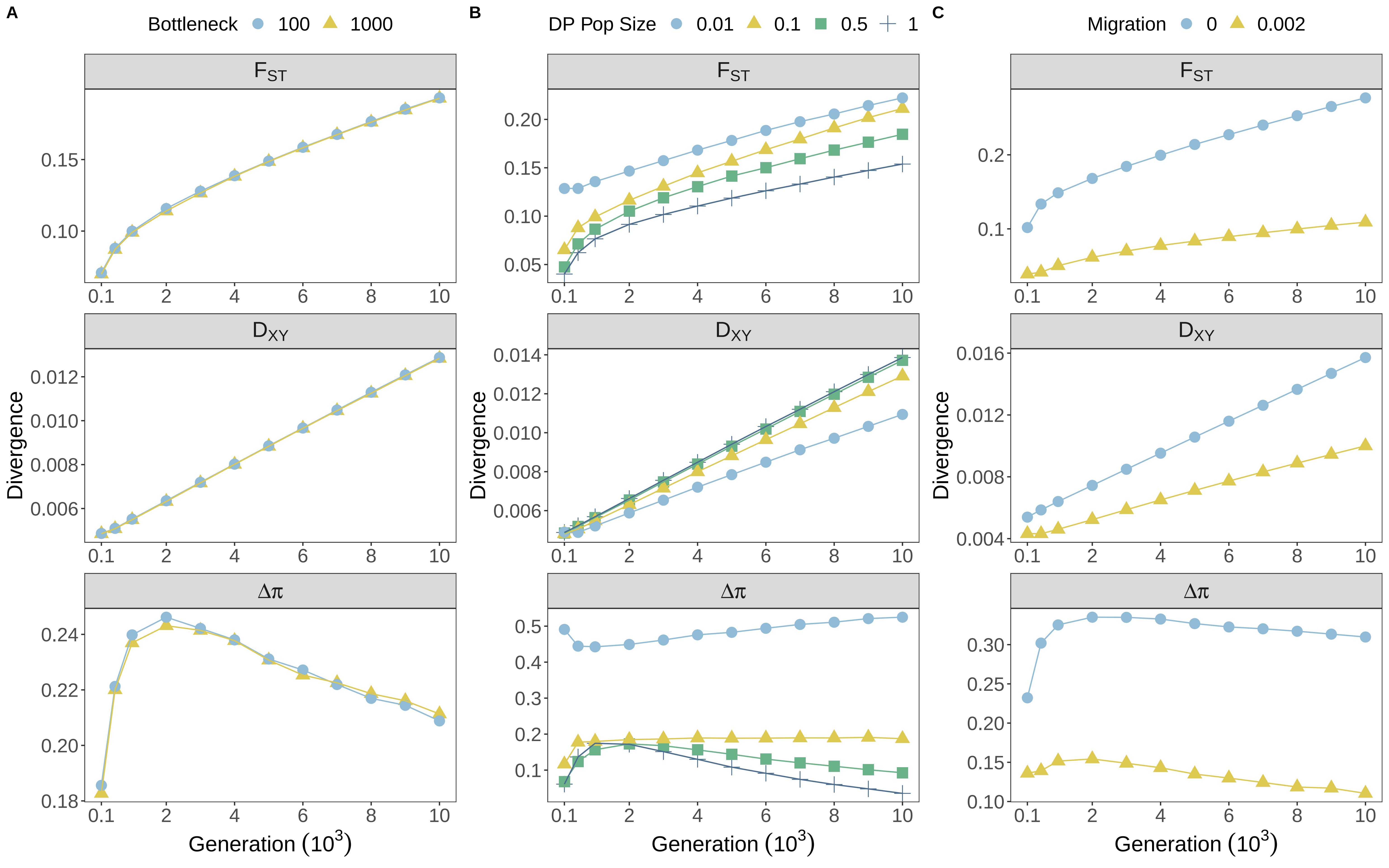

A
Bottleneck 100 1000
B
DP Pop Size 0.01 0.1 0.5 1
C
Migration 0 0.002
$F_{ST}$
$D_{XY}$
$\Delta\pi$
Divergence
Generation ($10^3$)

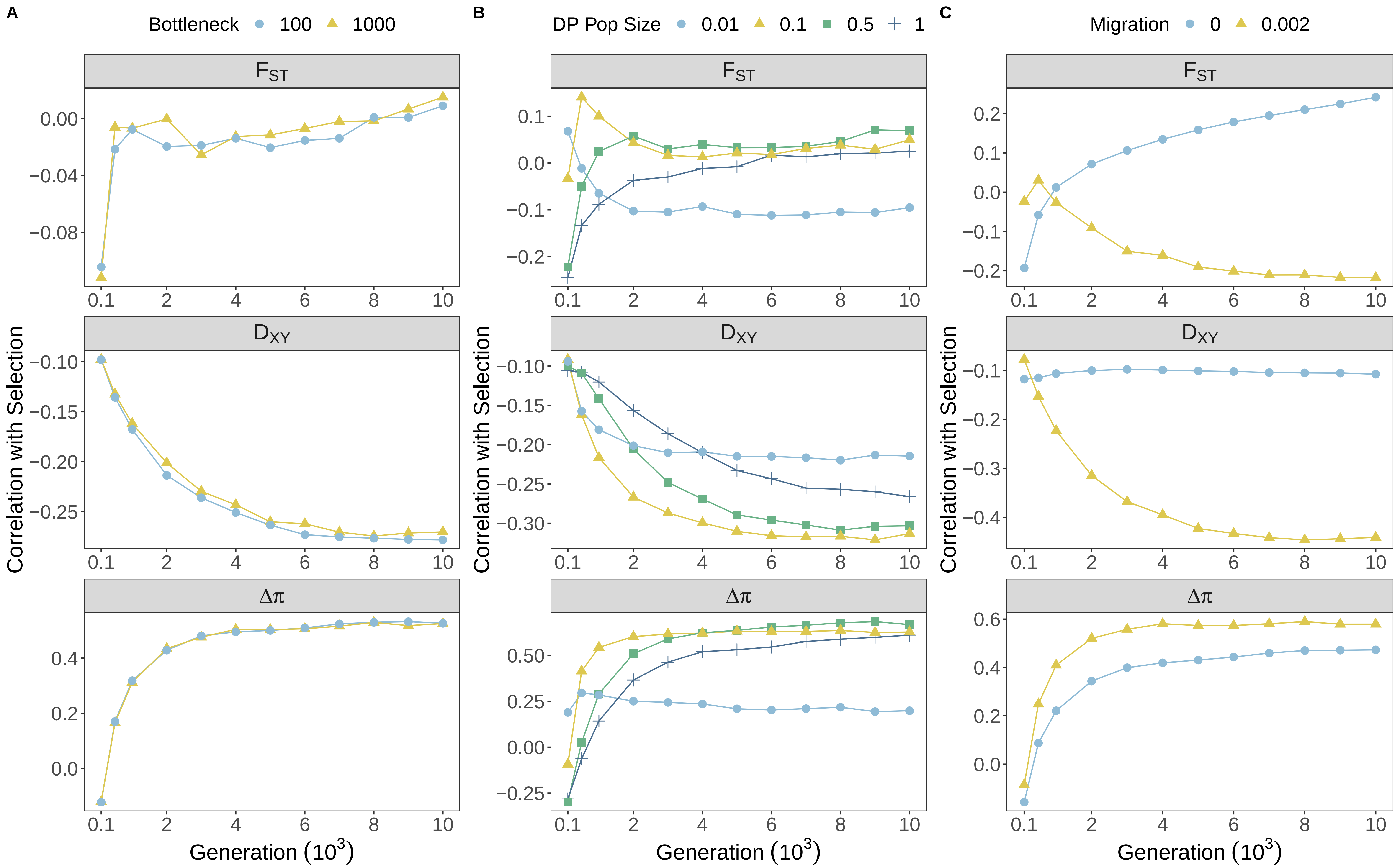
A
Bottleneck 100 1000
B
DP Pop Size 0.01 0.1 0.5 1
C
Migration 0 0.002
$F_{ST}$
$D_{XY}$
$\Delta\pi$
Correlation with Selection
Generation ($10^3$)

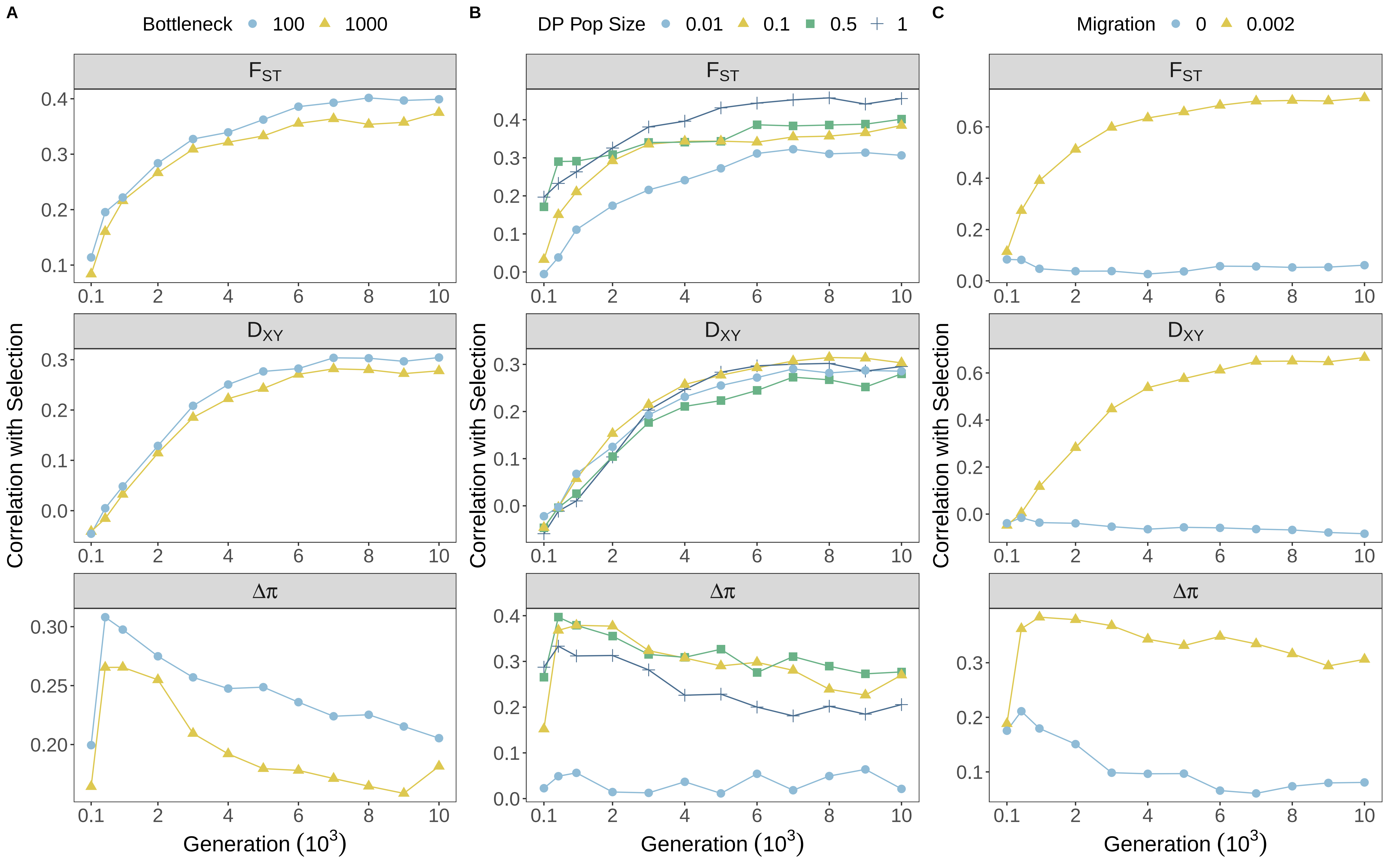
A
Bottleneck 100 1000
B
DP Pop Size 0.01 0.1 0.5 1
C
Migration 0 0.002
$F_{ST}$
$D_{XY}$
$\Delta\pi$
Correlation with Selection
Generation ($10^3$)

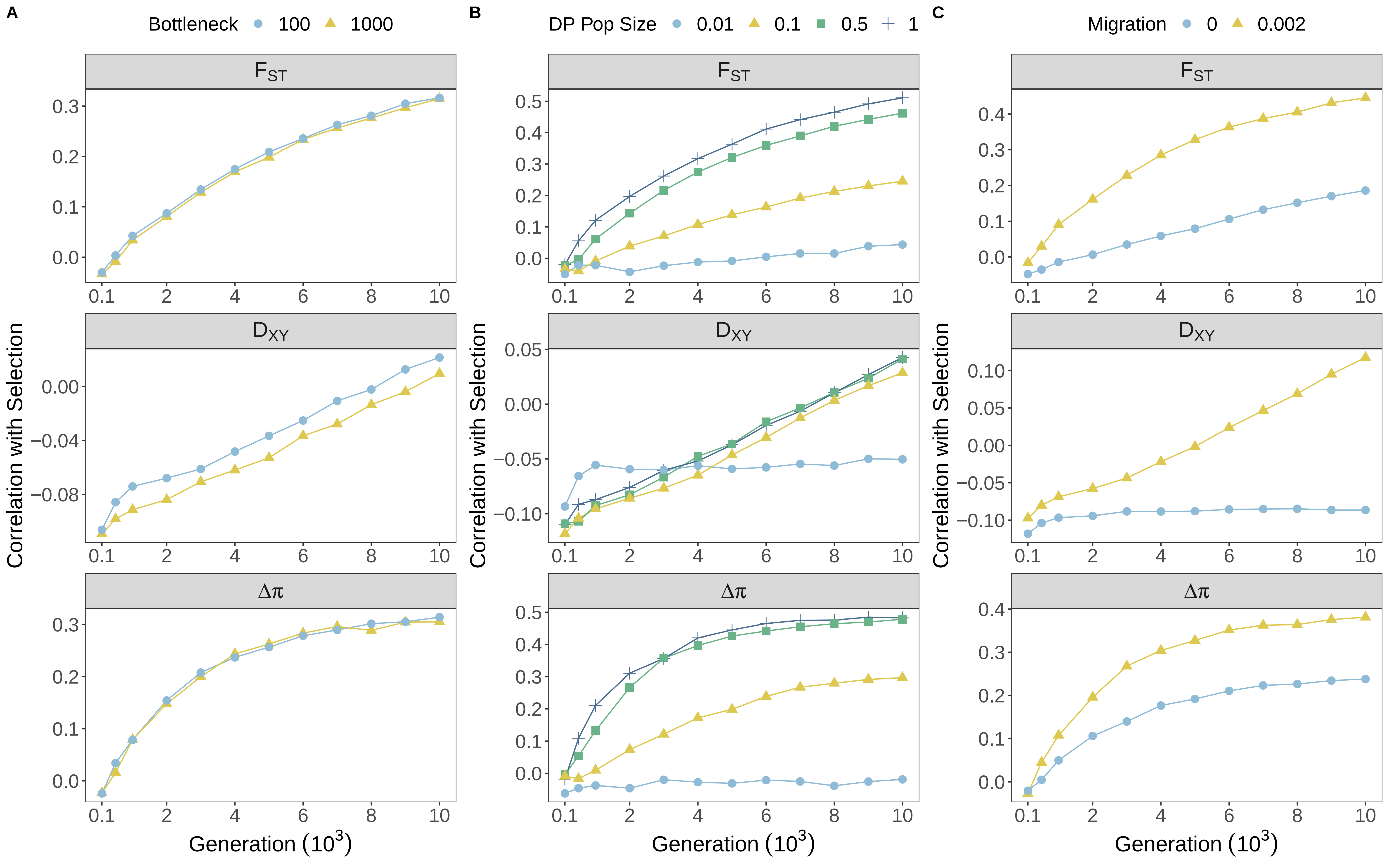
A
Bottleneck
100
1000
B
DP Pop Size
0.01
0.1
0.5
1
C
Migration
0
0.002
$F_{ST}$
$D_{XY}$
$\Delta\pi$
Correlation with Selection
Generation ($10^3$)

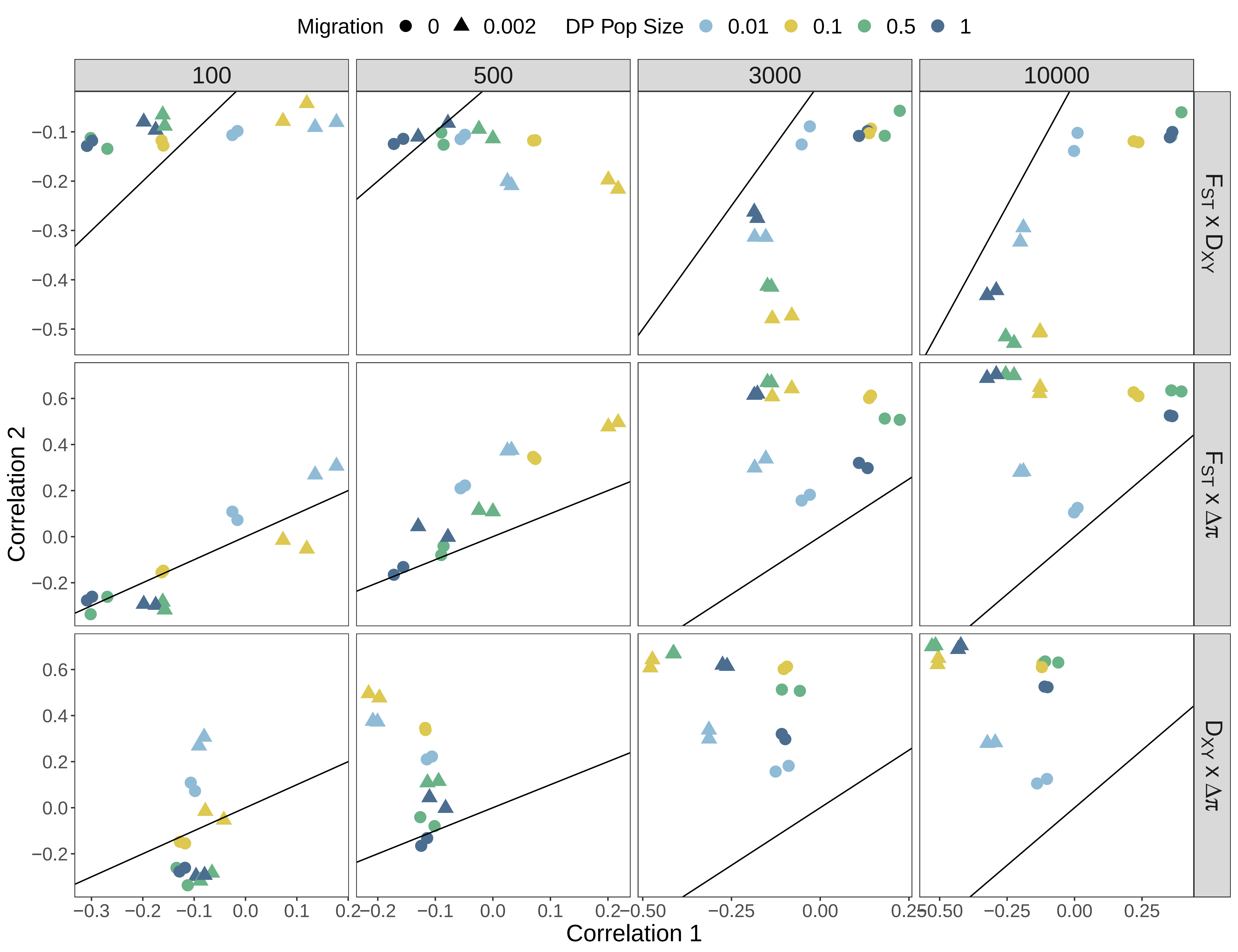

Migration 0 0.002 DP Pop Size 0.01 0.1 0.5 1
100
500
3000
10000
F_ST x D_XY
F_ST x Δπ
D_XY x Δπ
Correlation 2
Correlation 1

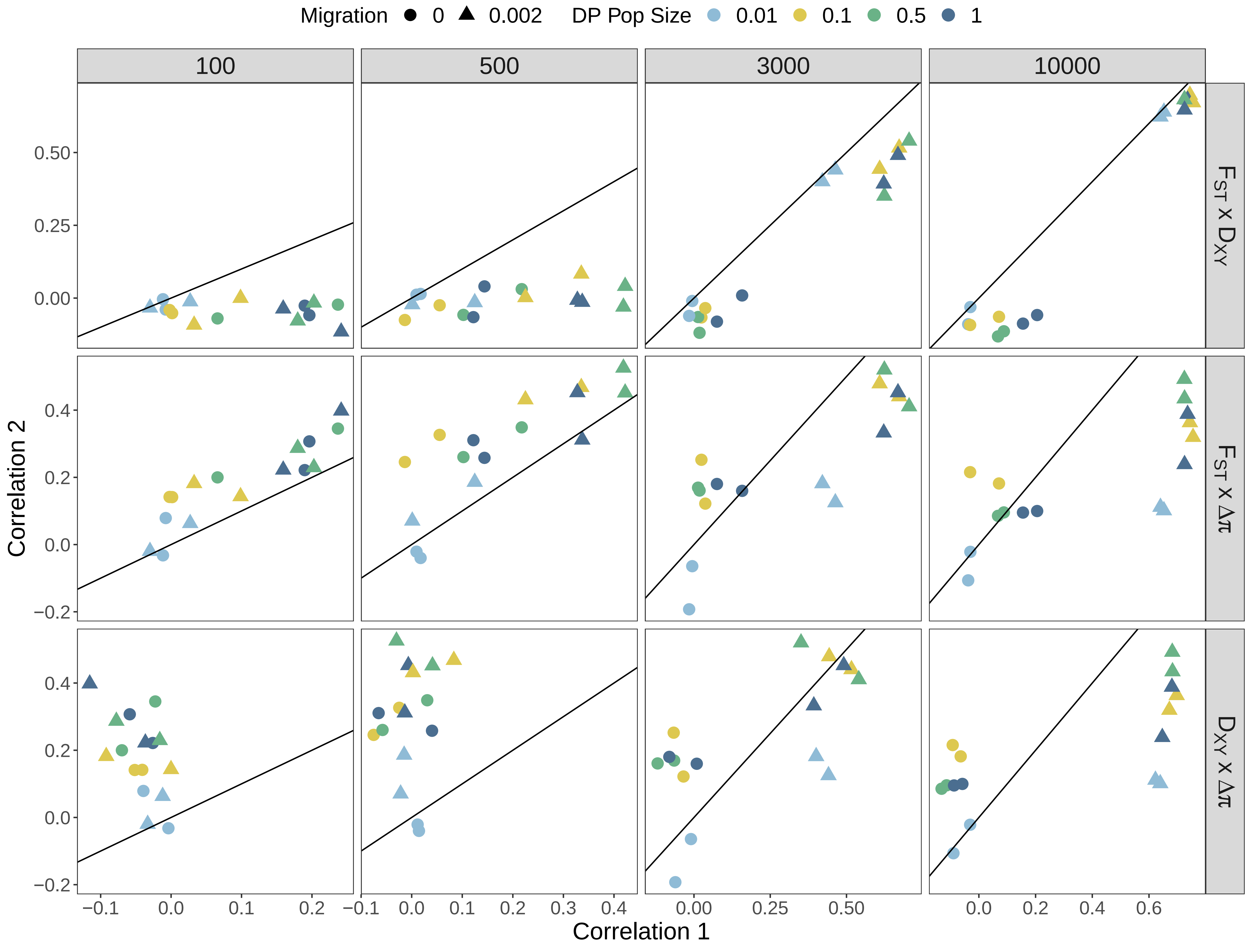

Migration 0 0.002 DP Pop Size 0.01 0.1 0.5 1
100
500
3000
10000
FST x DXY
FST x Δπ
DXY x Δπ
Correlation 2
Correlation 1

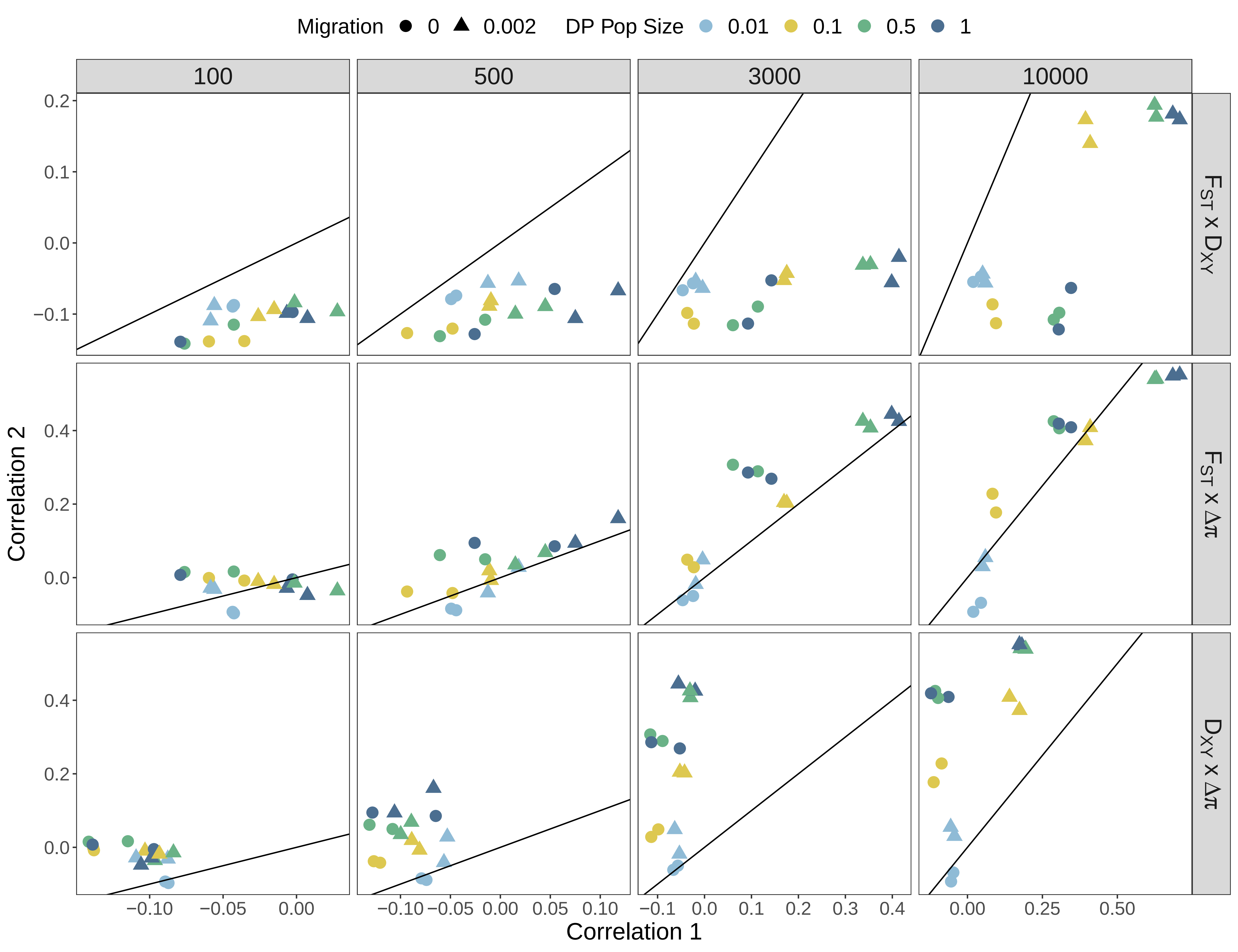

Migration 0 0.002 DP Pop Size 0.01 0.1 0.5 1
100
500
3000
10000
$F_{ST}$ x $D_{XY}$
$F_{ST}$ x $\Delta\pi$
$D_{XY}$ x $\Delta\pi$
Correlation 2
Correlation 1

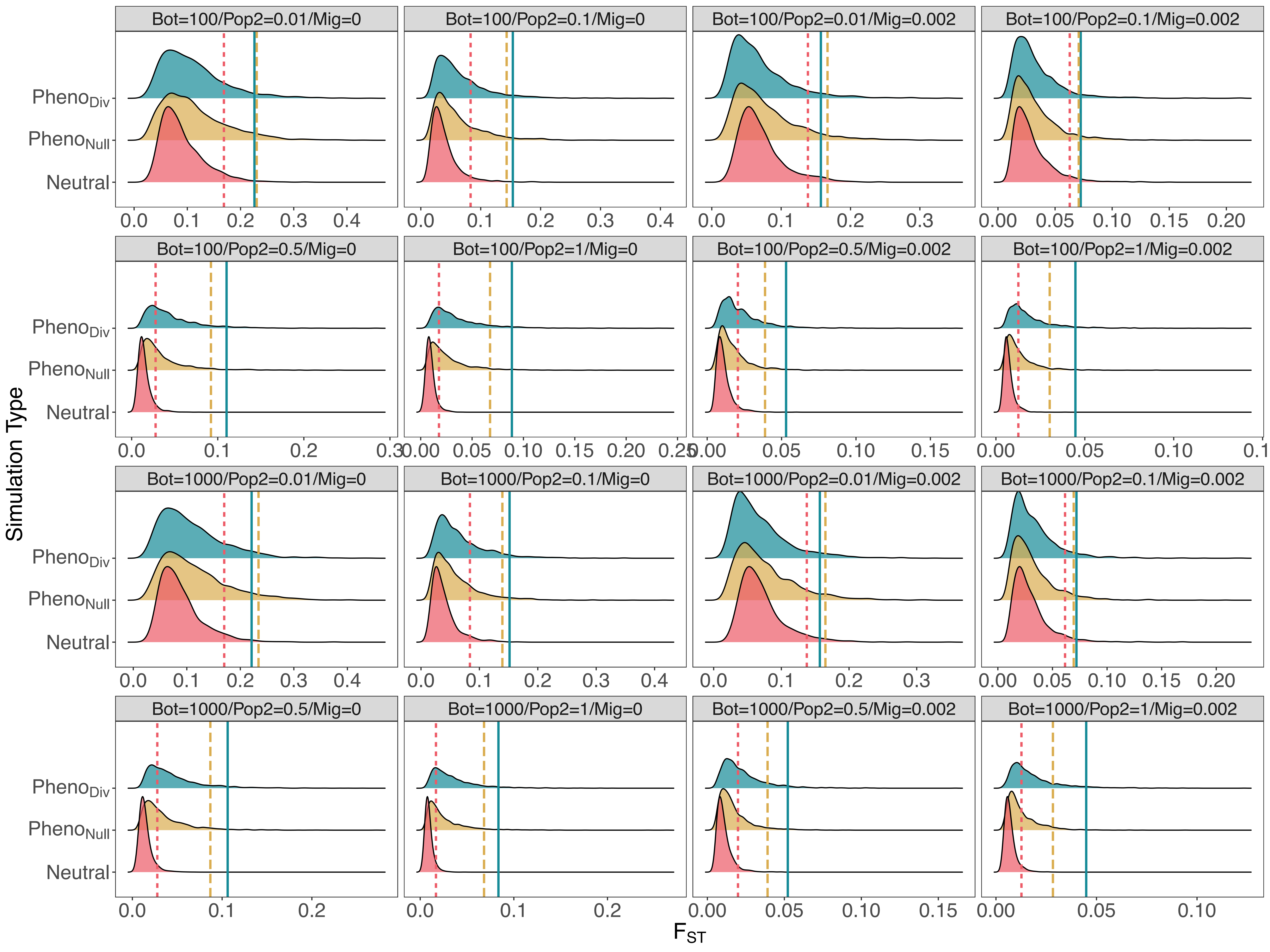

Bot=100/Pop2=0.01/Mig=0
Bot=100/Pop2=0.1/Mig=0
Bot=100/Pop2=0.01/Mig=0.002
Bot=100/Pop2=0.1/Mig=0.002
Bot=100/Pop2=0.5/Mig=0
Bot=100/Pop2=1/Mig=0
Bot=100/Pop2=0.5/Mig=0.002
Bot=100/Pop2=1/Mig=0.002
Bot=1000/Pop2=0.01/Mig=0
Bot=1000/Pop2=0.1/Mig=0
Bot=1000/Pop2=0.01/Mig=0.002
Bot=1000/Pop2=0.1/Mig=0.002
Bot=1000/Pop2=0.5/Mig=0
Bot=1000/Pop2=1/Mig=0
Bot=1000/Pop2=0.5/Mig=0.002
Bot=1000/Pop2=1/Mig=0.002
$Pheno_{Div}$
$Pheno_{Null}$
Neutral
Simulation Type
$F_{ST}$

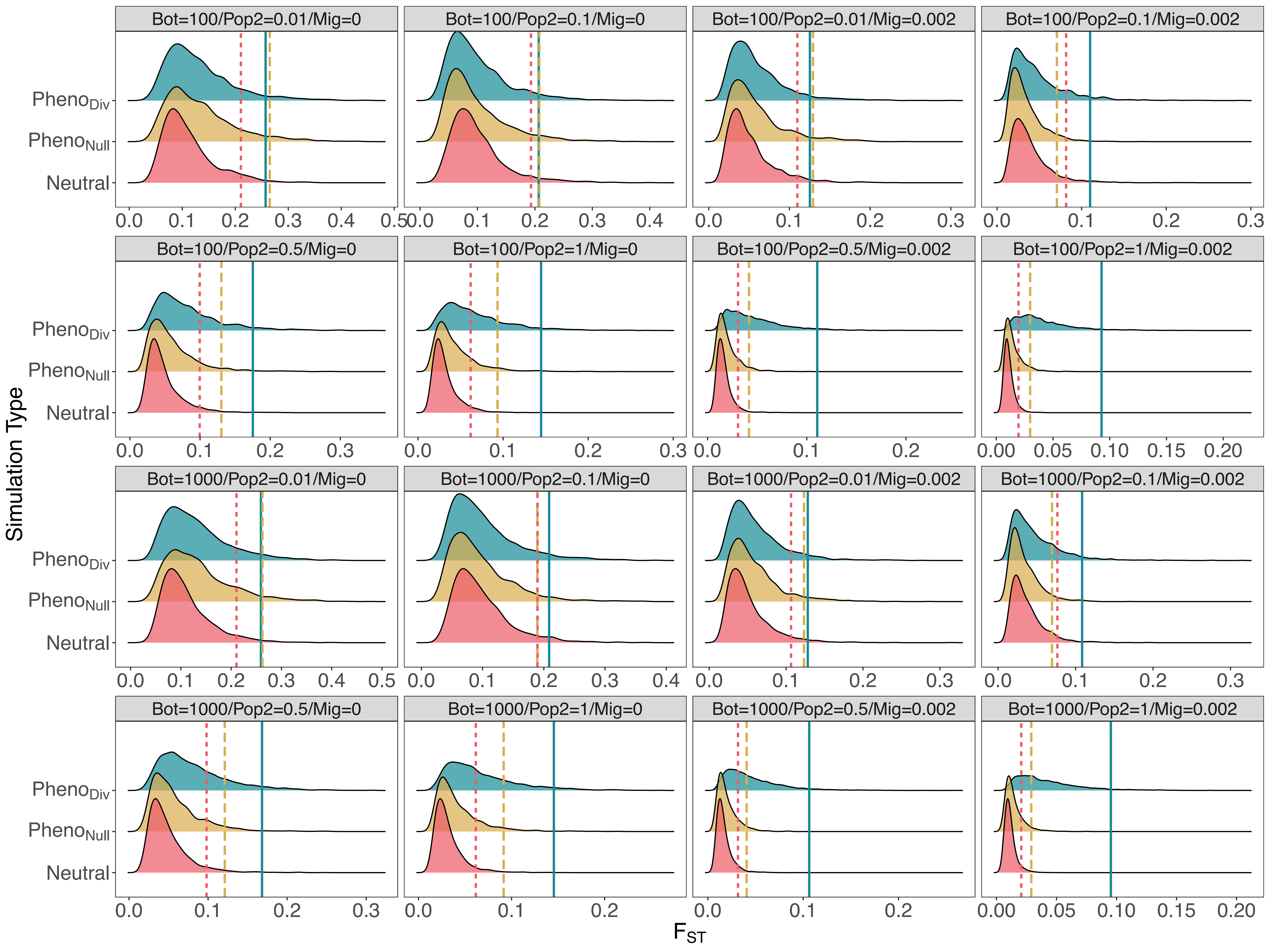
Bot=100/Pop2=0.01/Mig=0
Bot=100/Pop2=0.1/Mig=0
Bot=100/Pop2=0.01/Mig=0.002
Bot=100/Pop2=0.1/Mig=0.002
Bot=100/Pop2=0.5/Mig=0
Bot=100/Pop2=1/Mig=0
Bot=100/Pop2=0.5/Mig=0.002
Bot=100/Pop2=1/Mig=0.002
Bot=1000/Pop2=0.01/Mig=0
Bot=1000/Pop2=0.1/Mig=0
Bot=1000/Pop2=0.01/Mig=0.002
Bot=1000/Pop2=0.1/Mig=0.002
Bot=1000/Pop2=0.5/Mig=0
Bot=1000/Pop2=1/Mig=0
Bot=1000/Pop2=0.5/Mig=0.002
Bot=1000/Pop2=1/Mig=0.002
Pheno$_{Div}$
Pheno$_{Null}$
Neutral
Simulation Type
$F_{ST}$

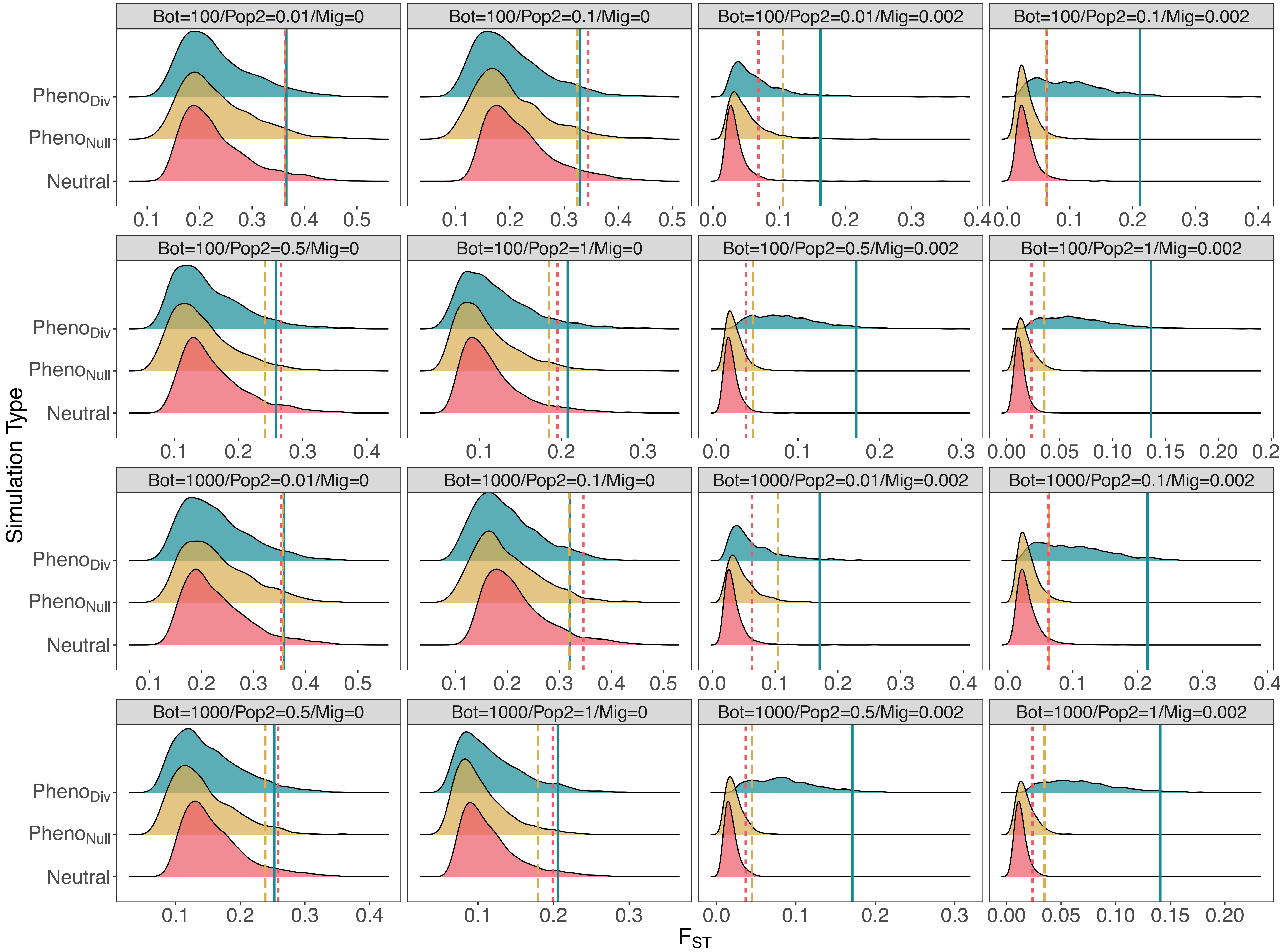
Bot=100/Pop2=0.01/Mig=0
Bot=100/Pop2=0.1/Mig=0
Bot=100/Pop2=0.01/Mig=0.002
Bot=100/Pop2=0.1/Mig=0.002
Bot=100/Pop2=0.5/Mig=0
Bot=100/Pop2=1/Mig=0
Bot=100/Pop2=0.5/Mig=0.002
Bot=100/Pop2=1/Mig=0.002
Bot=1000/Pop2=0.01/Mig=0
Bot=1000/Pop2=0.1/Mig=0
Bot=1000/Pop2=0.01/Mig=0.002
Bot=1000/Pop2=0.1/Mig=0.002
Bot=1000/Pop2=0.5/Mig=0
Bot=1000/Pop2=1/Mig=0
Bot=1000/Pop2=0.5/Mig=0.002
Bot=1000/Pop2=1/Mig=0.002
$Pheno_{Div}$
$Pheno_{Null}$
Neutral
Simulation Type
$F_{ST}$

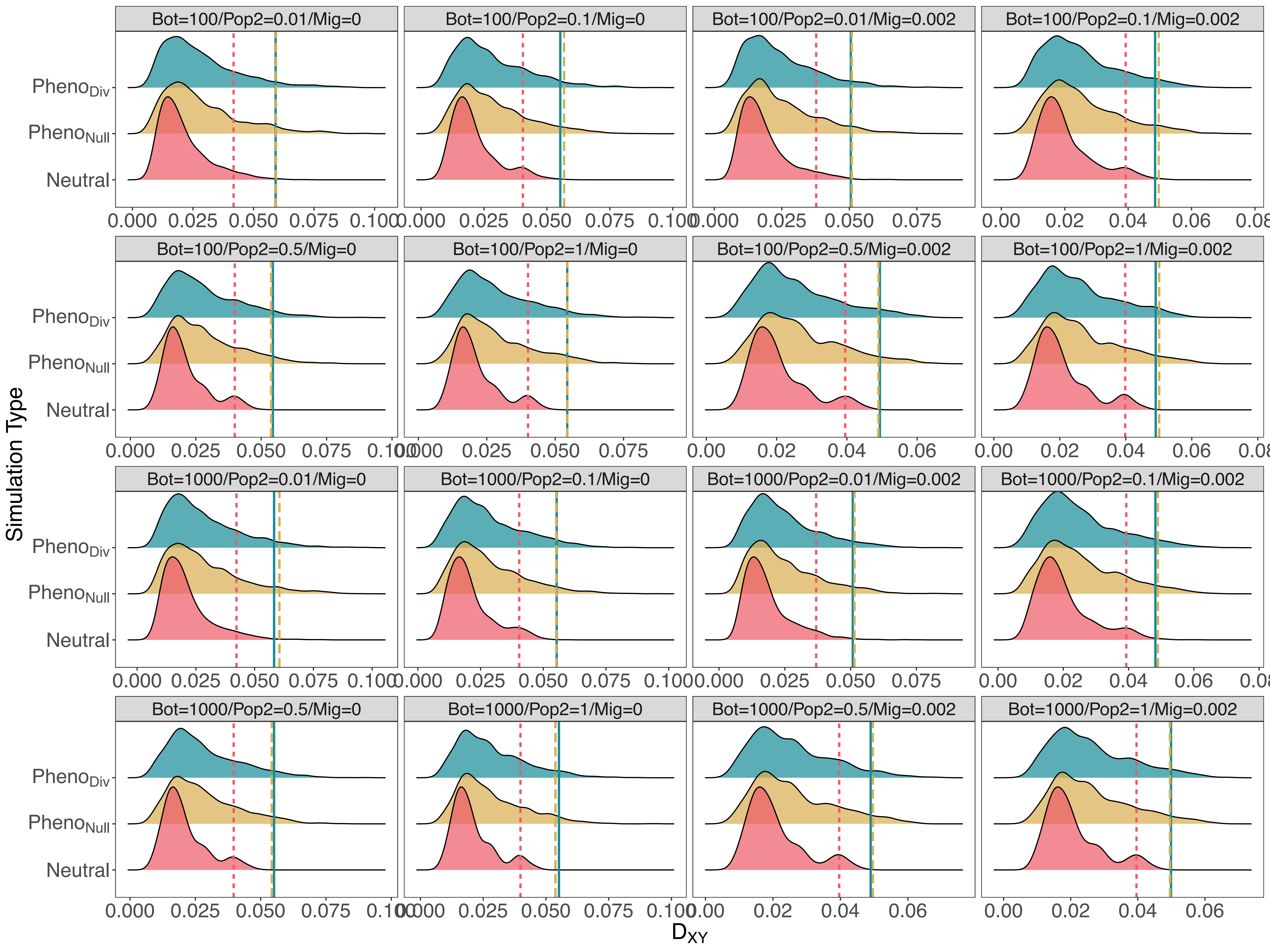
Bot=100/Pop2=0.01/Mig=0
Bot=100/Pop2=0.1/Mig=0
Bot=100/Pop2=0.01/Mig=0.002
Bot=100/Pop2=0.1/Mig=0.002
Bot=100/Pop2=0.5/Mig=0
Bot=100/Pop2=1/Mig=0
Bot=100/Pop2=0.5/Mig=0.002
Bot=100/Pop2=1/Mig=0.002
Bot=1000/Pop2=0.01/Mig=0
Bot=1000/Pop2=0.1/Mig=0
Bot=1000/Pop2=0.01/Mig=0.002
Bot=1000/Pop2=0.1/Mig=0.002
Bot=1000/Pop2=0.5/Mig=0
Bot=1000/Pop2=1/Mig=0
Bot=1000/Pop2=0.5/Mig=0.002
Bot=1000/Pop2=1/Mig=0.002
Simulation Type
Pheno$_{Div}$
Pheno$_{Null}$
Neutral
$D_{XY}$

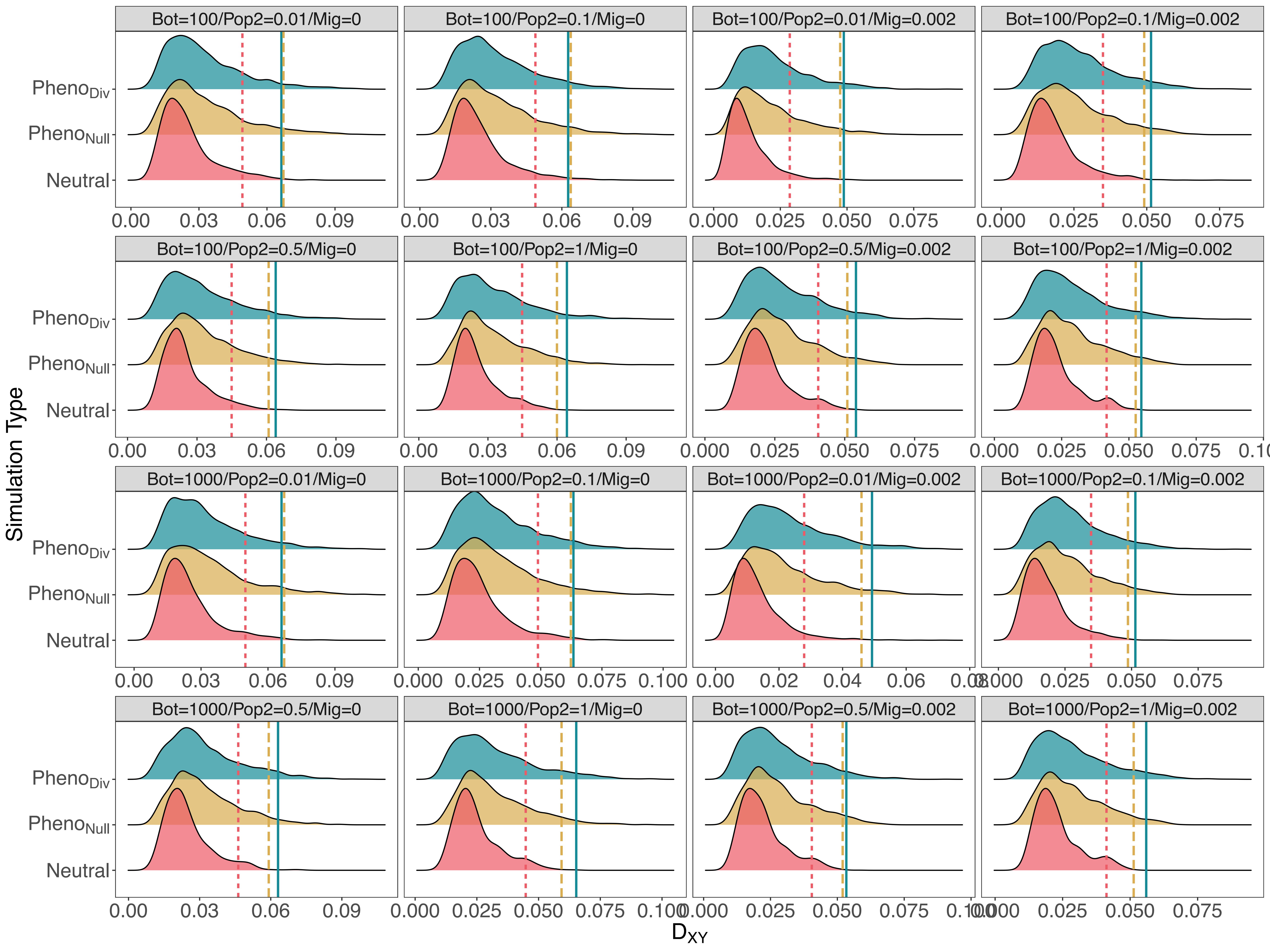

Bot=100/Pop2=0.01/Mig=0
Bot=100/Pop2=0.1/Mig=0
Bot=100/Pop2=0.01/Mig=0.002
Bot=100/Pop2=0.1/Mig=0.002
Bot=100/Pop2=0.5/Mig=0
Bot=100/Pop2=1/Mig=0
Bot=100/Pop2=0.5/Mig=0.002
Bot=100/Pop2=1/Mig=0.002
Bot=1000/Pop2=0.01/Mig=0
Bot=1000/Pop2=0.1/Mig=0
Bot=1000/Pop2=0.01/Mig=0.002
Bot=1000/Pop2=0.1/Mig=0.002
Bot=1000/Pop2=0.5/Mig=0
Bot=1000/Pop2=1/Mig=0
Bot=1000/Pop2=0.5/Mig=0.002
Bot=1000/Pop2=1/Mig=0.002
Pheno$_{Div}$
Pheno$_{Null}$
Neutral
Simulation Type
D$_{XY}$

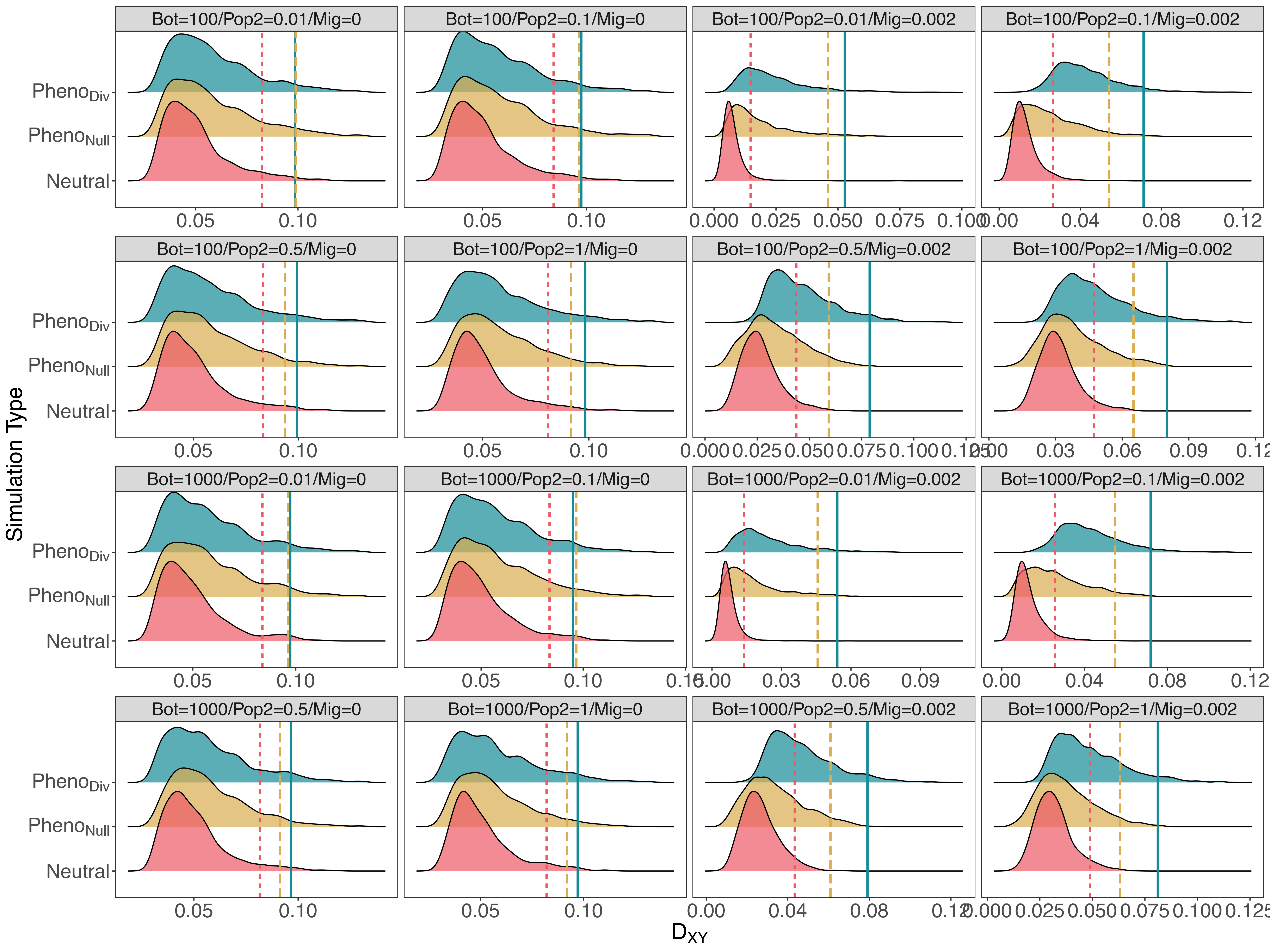

Bot=100/Pop2=0.01/Mig=0
Bot=100/Pop2=0.1/Mig=0
Bot=100/Pop2=0.01/Mig=0.002
Bot=100/Pop2=0.1/Mig=0.002
Bot=100/Pop2=0.5/Mig=0
Bot=100/Pop2=1/Mig=0
Bot=100/Pop2=0.5/Mig=0.002
Bot=100/Pop2=1/Mig=0.002
Bot=1000/Pop2=0.01/Mig=0
Bot=1000/Pop2=0.1/Mig=0
Bot=1000/Pop2=0.01/Mig=0.002
Bot=1000/Pop2=0.1/Mig=0.002
Bot=1000/Pop2=0.5/Mig=0
Bot=1000/Pop2=1/Mig=0
Bot=1000/Pop2=0.5/Mig=0.002
Bot=1000/Pop2=1/Mig=0.002
$Pheno_{Div}$
$Pheno_{Null}$
Neutral
Simulation Type
$D_{XY}$

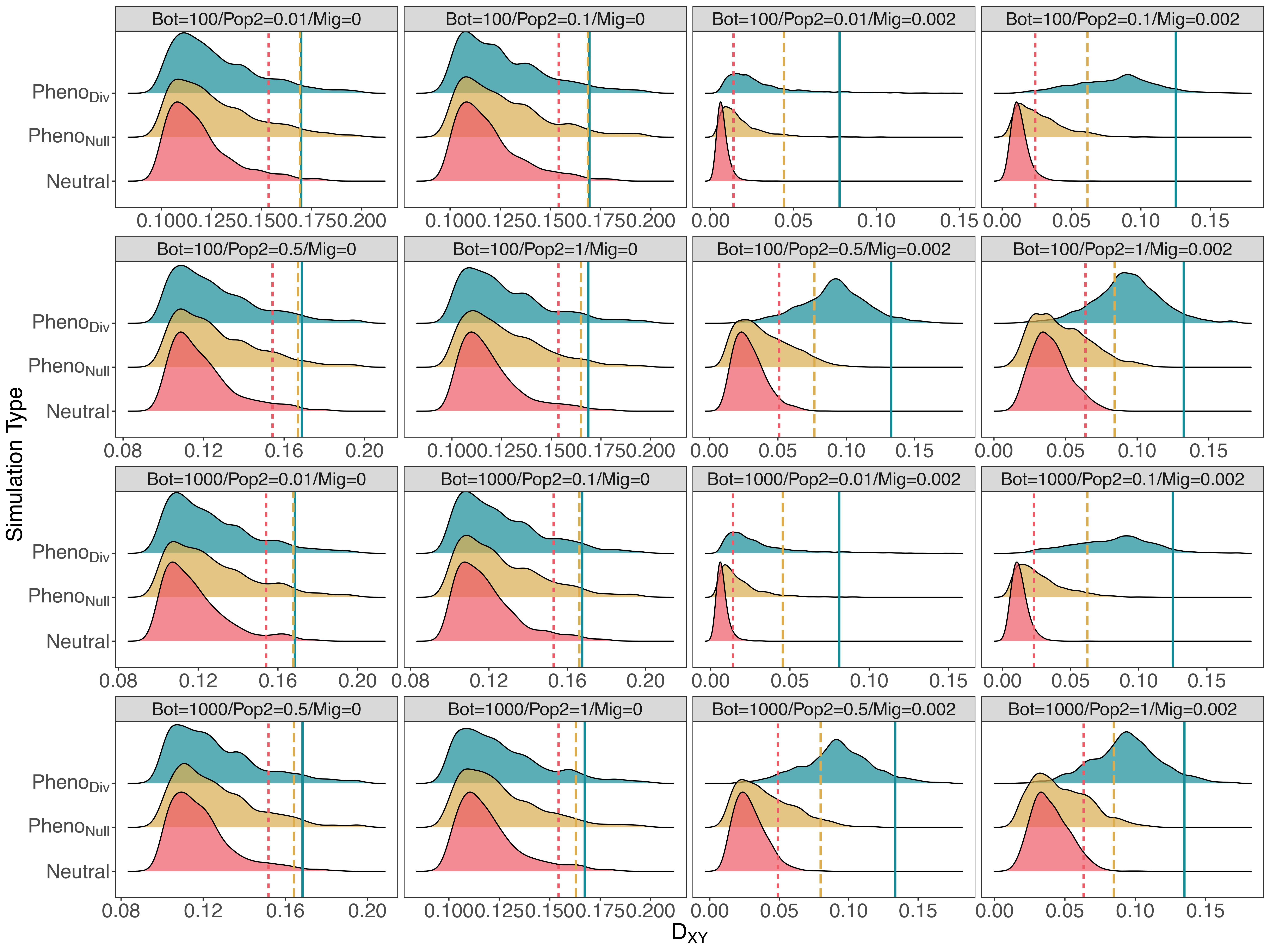

Bot=100/Pop2=0.01/Mig=0
Bot=100/Pop2=0.1/Mig=0
Bot=100/Pop2=0.01/Mig=0.002
Bot=100/Pop2=0.1/Mig=0.002
Bot=100/Pop2=0.5/Mig=0
Bot=100/Pop2=1/Mig=0
Bot=100/Pop2=0.5/Mig=0.002
Bot=100/Pop2=1/Mig=0.002
Bot=1000/Pop2=0.01/Mig=0
Bot=1000/Pop2=0.1/Mig=0
Bot=1000/Pop2=0.01/Mig=0.002
Bot=1000/Pop2=0.1/Mig=0.002
Bot=1000/Pop2=0.5/Mig=0
Bot=1000/Pop2=1/Mig=0
Bot=1000/Pop2=0.5/Mig=0.002
Bot=1000/Pop2=1/Mig=0.002
$Pheno_{Div}$
$Pheno_{Null}$
Neutral
Simulation Type
$D_{XY}$

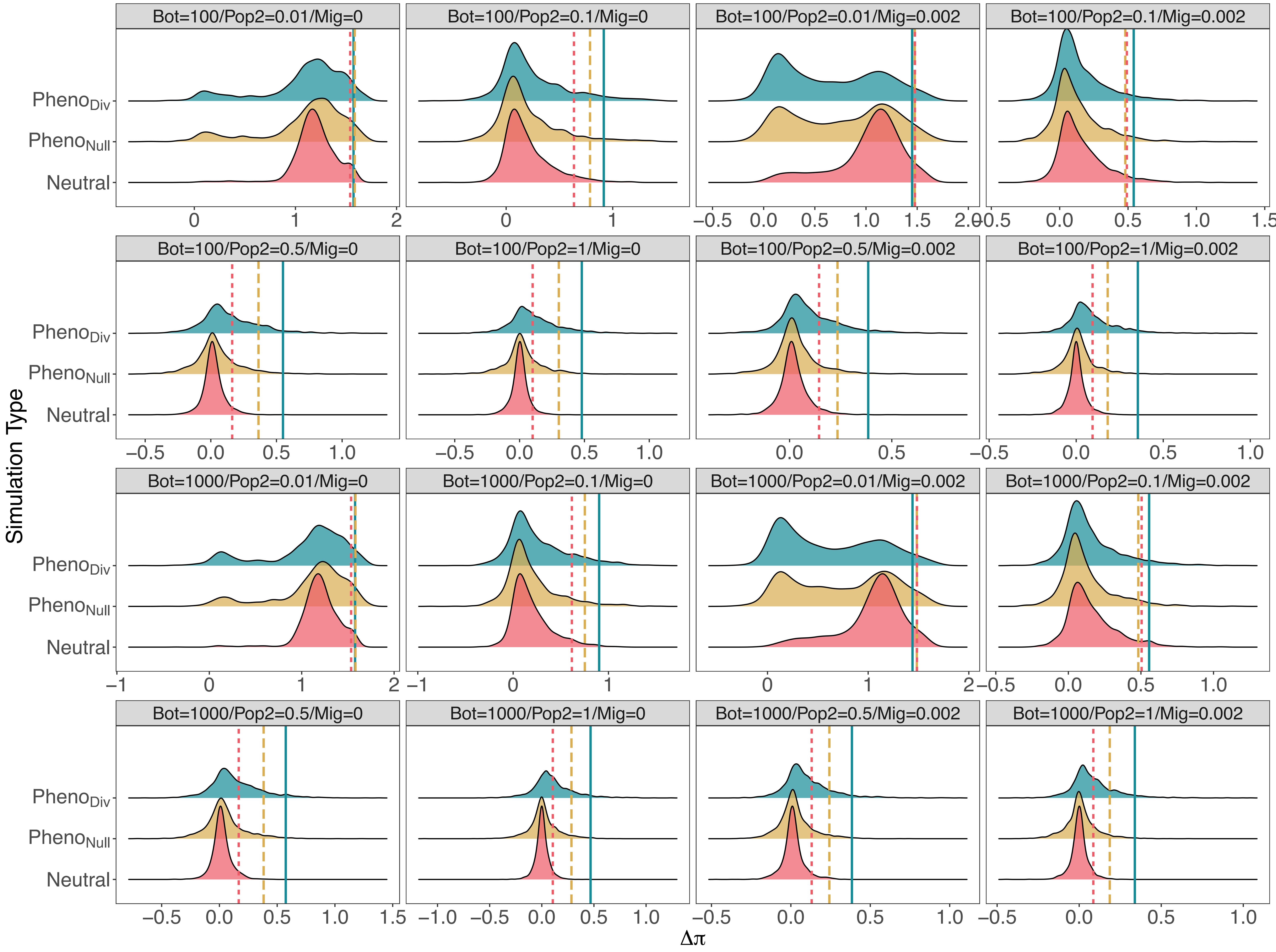

Bot=100/Pop2=0.01/Mig=0
Bot=100/Pop2=0.1/Mig=0
Bot=100/Pop2=0.01/Mig=0.002
Bot=100/Pop2=0.1/Mig=0.002
Bot=100/Pop2=0.5/Mig=0
Bot=100/Pop2=1/Mig=0
Bot=100/Pop2=0.5/Mig=0.002
Bot=100/Pop2=1/Mig=0.002
Bot=1000/Pop2=0.01/Mig=0
Bot=1000/Pop2=0.1/Mig=0
Bot=1000/Pop2=0.01/Mig=0.002
Bot=1000/Pop2=0.1/Mig=0.002
Bot=1000/Pop2=0.5/Mig=0
Bot=1000/Pop2=1/Mig=0
Bot=1000/Pop2=0.5/Mig=0.002
Bot=1000/Pop2=1/Mig=0.002
$Pheno_{Div}$
$Pheno_{Null}$
Neutral
Simulation Type
$\Delta\pi$

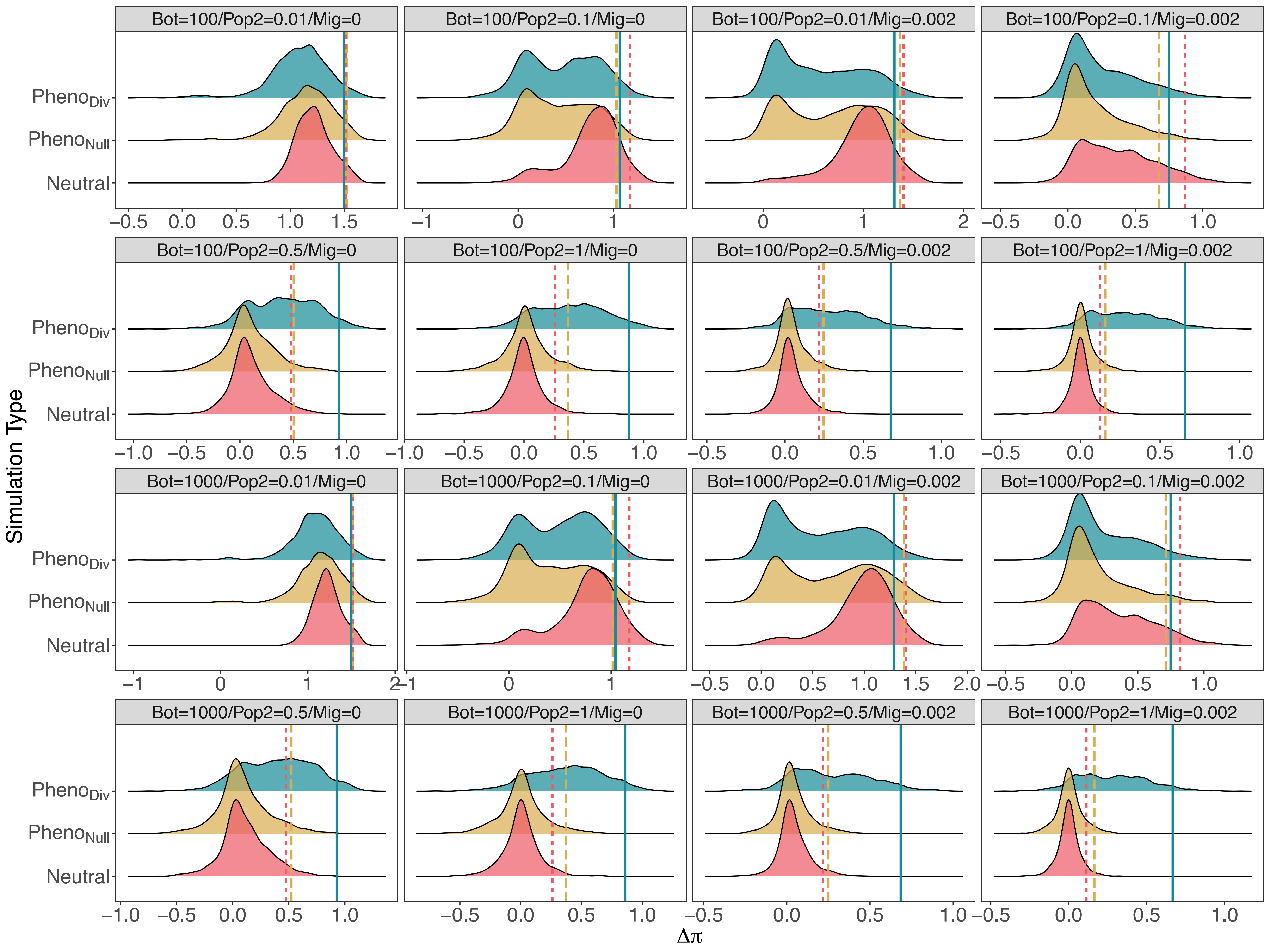

Bot=100/Pop2=0.01/Mig=0
Bot=100/Pop2=0.1/Mig=0
Bot=100/Pop2=0.01/Mig=0.002
Bot=100/Pop2=0.1/Mig=0.002
Bot=100/Pop2=0.5/Mig=0
Bot=100/Pop2=1/Mig=0
Bot=100/Pop2=0.5/Mig=0.002
Bot=100/Pop2=1/Mig=0.002
Bot=1000/Pop2=0.01/Mig=0
Bot=1000/Pop2=0.1/Mig=0
Bot=1000/Pop2=0.01/Mig=0.002
Bot=1000/Pop2=0.1/Mig=0.002
Bot=1000/Pop2=0.5/Mig=0
Bot=1000/Pop2=1/Mig=0
Bot=1000/Pop2=0.5/Mig=0.002
Bot=1000/Pop2=1/Mig=0.002
$Pheno_{Div}$
$Pheno_{Null}$
Neutral
Simulation Type
$\Delta\pi$

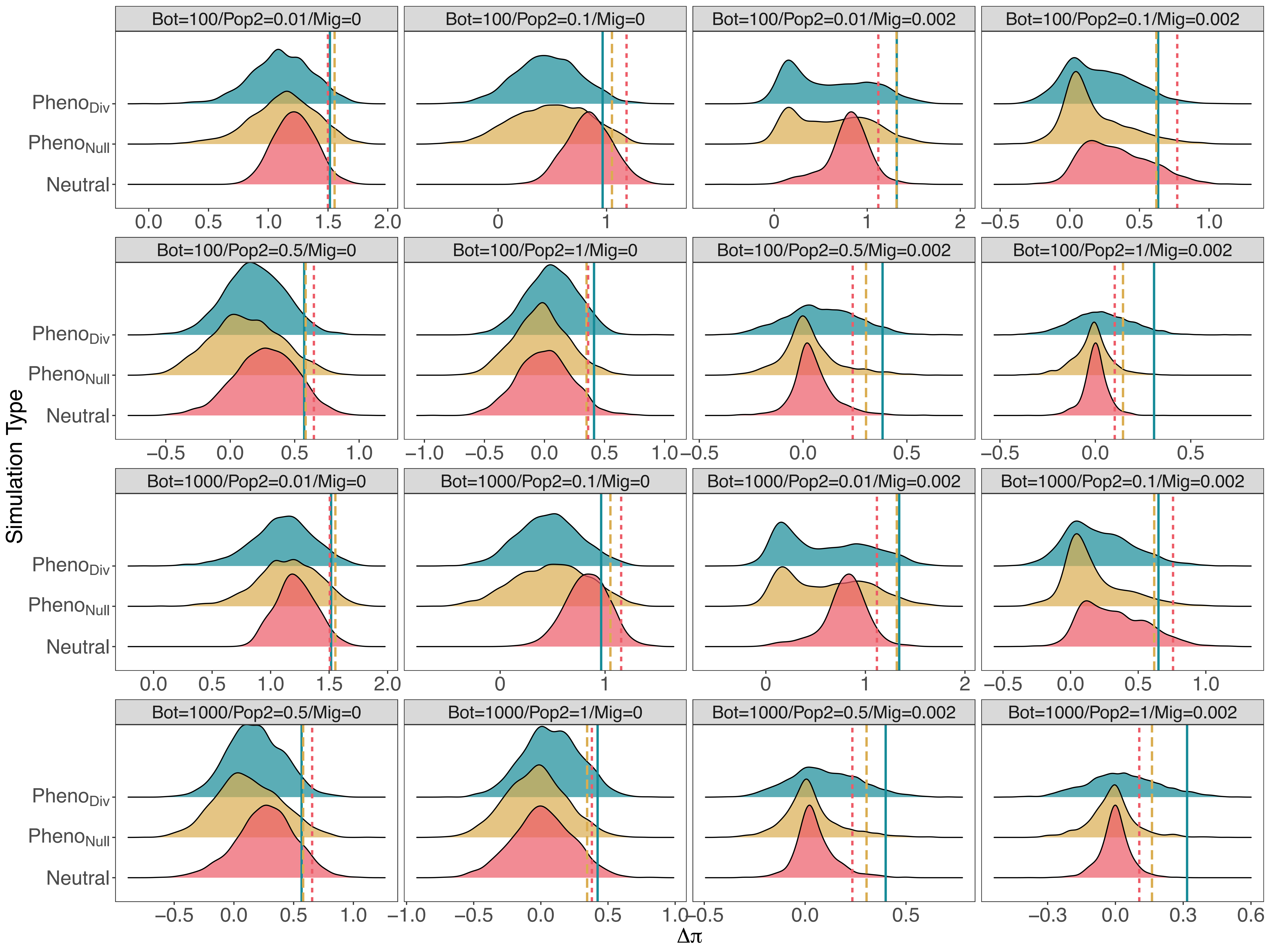

Bot=100/Pop2=0.01/Mig=0
Bot=100/Pop2=0.1/Mig=0
Bot=100/Pop2=0.01/Mig=0.002
Bot=100/Pop2=0.1/Mig=0.002
Bot=100/Pop2=0.5/Mig=0
Bot=100/Pop2=1/Mig=0
Bot=100/Pop2=0.5/Mig=0.002
Bot=100/Pop2=1/Mig=0.002
Bot=1000/Pop2=0.01/Mig=0
Bot=1000/Pop2=0.1/Mig=0
Bot=1000/Pop2=0.01/Mig=0.002
Bot=1000/Pop2=0.1/Mig=0.002
Bot=1000/Pop2=0.5/Mig=0
Bot=1000/Pop2=1/Mig=0
Bot=1000/Pop2=0.5/Mig=0.002
Bot=1000/Pop2=1/Mig=0.002
$Pheno_{Div}$
$Pheno_{Null}$
Neutral
Simulation Type
$\Delta\pi$

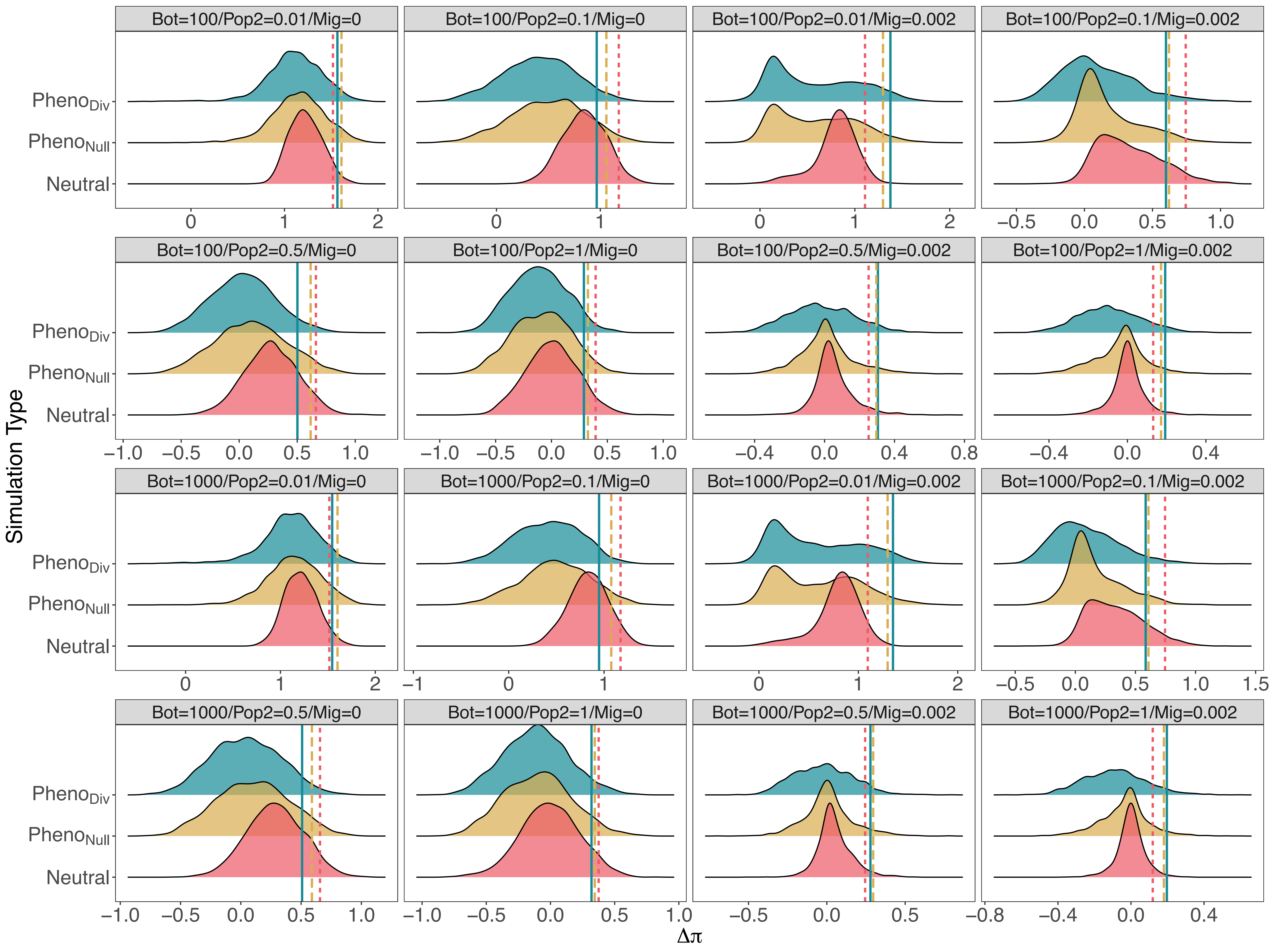

Bot=100/Pop2=0.01/Mig=0
Bot=100/Pop2=0.1/Mig=0
Bot=100/Pop2=0.01/Mig=0.002
Bot=100/Pop2=0.1/Mig=0.002
Bot=100/Pop2=0.5/Mig=0
Bot=100/Pop2=1/Mig=0
Bot=100/Pop2=0.5/Mig=0.002
Bot=100/Pop2=1/Mig=0.002
Bot=1000/Pop2=0.01/Mig=0
Bot=1000/Pop2=0.1/Mig=0
Bot=1000/Pop2=0.01/Mig=0.002
Bot=1000/Pop2=0.1/Mig=0.002
Bot=1000/Pop2=0.5/Mig=0
Bot=1000/Pop2=1/Mig=0
Bot=1000/Pop2=0.5/Mig=0.002
Bot=1000/Pop2=1/Mig=0.002
Simulation Type
$Pheno_{Div}$
$Pheno_{Null}$
Neutral
$\Delta\pi$

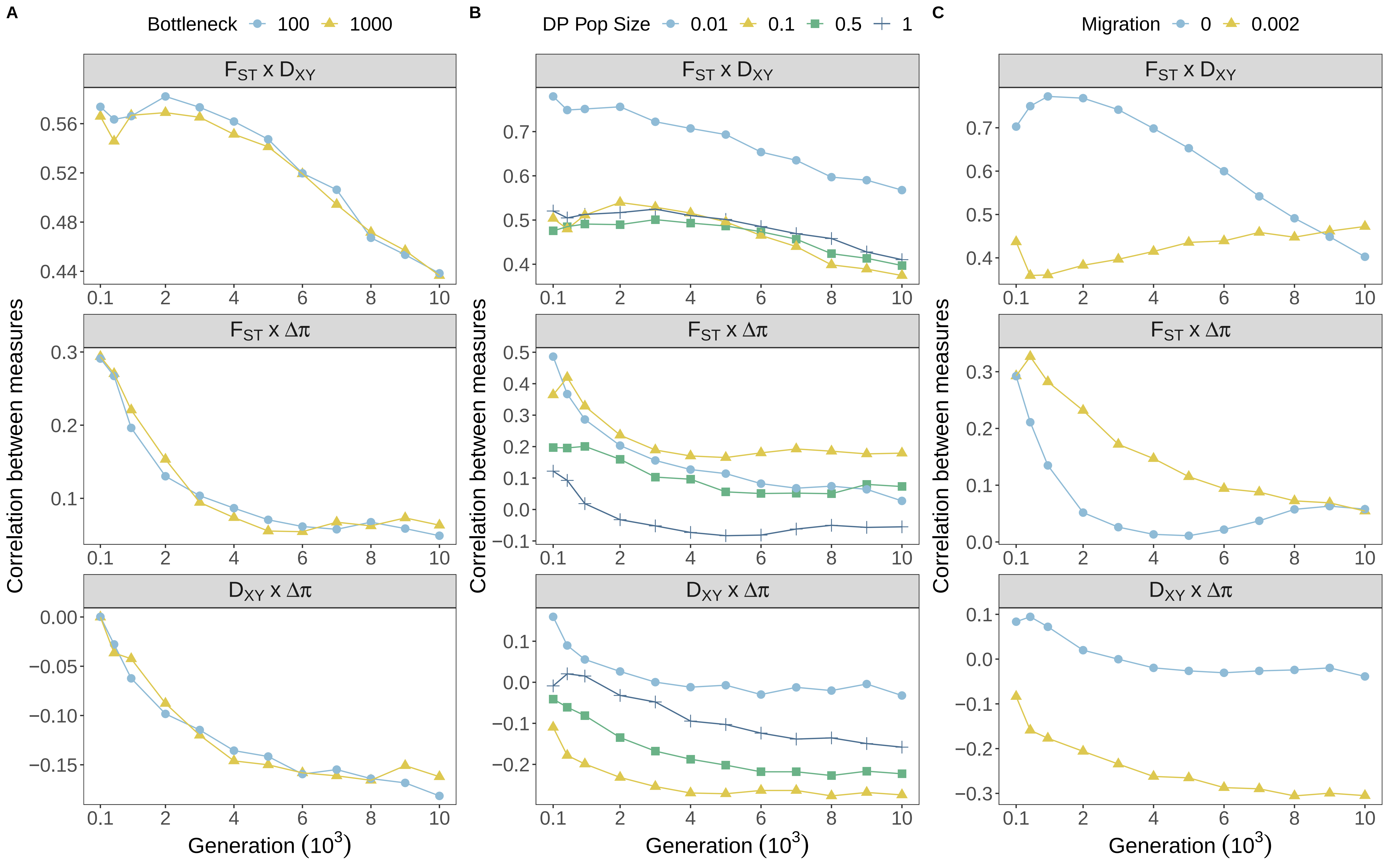
A
Bottleneck 100 1000
B
DP Pop Size 0.01 0.1 0.5 1
C
Migration 0 0.002
$F_{ST}$ x $D_{XY}$
$F_{ST}$ x $\Delta\pi$
$D_{XY}$ x $\Delta\pi$
Correlation between measures
Generation ($10^3$)

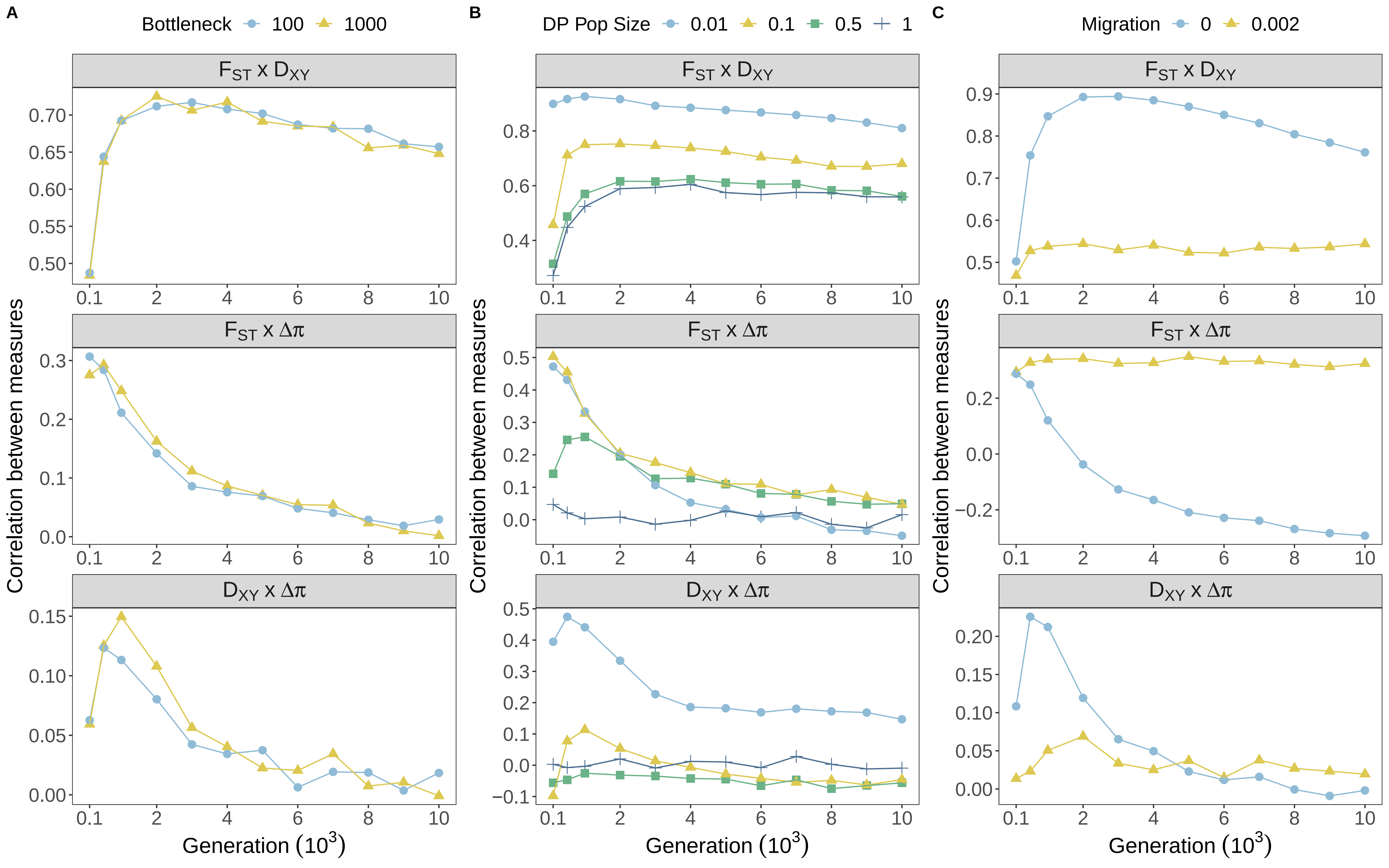

A
Bottleneck 100 1000
B
DP Pop Size 0.01 0.1 0.5 1
C
Migration 0 0.002
$F_{ST}$ x $D_{XY}$
$F_{ST}$ x $\Delta\pi$
$D_{XY}$ x $\Delta\pi$
Correlation between measures
Generation ($10^3$)

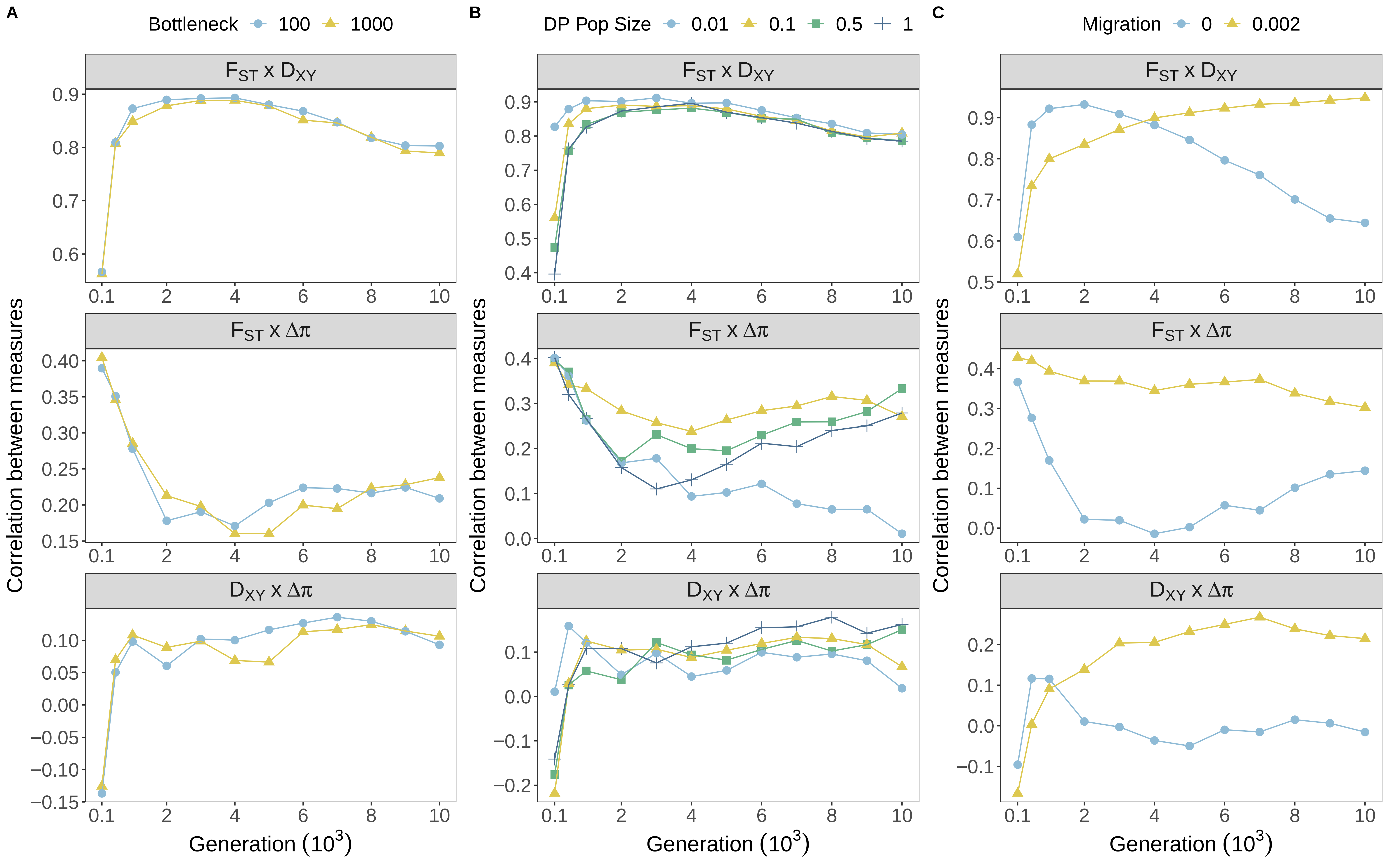
A
Bottleneck 100 1000
B
DP Pop Size 0.01 0.1 0.5 1
C
Migration 0 0.002
FST x DXY
FST x Δπ
DXY x Δπ
Correlation between measures
Generation (10^3)

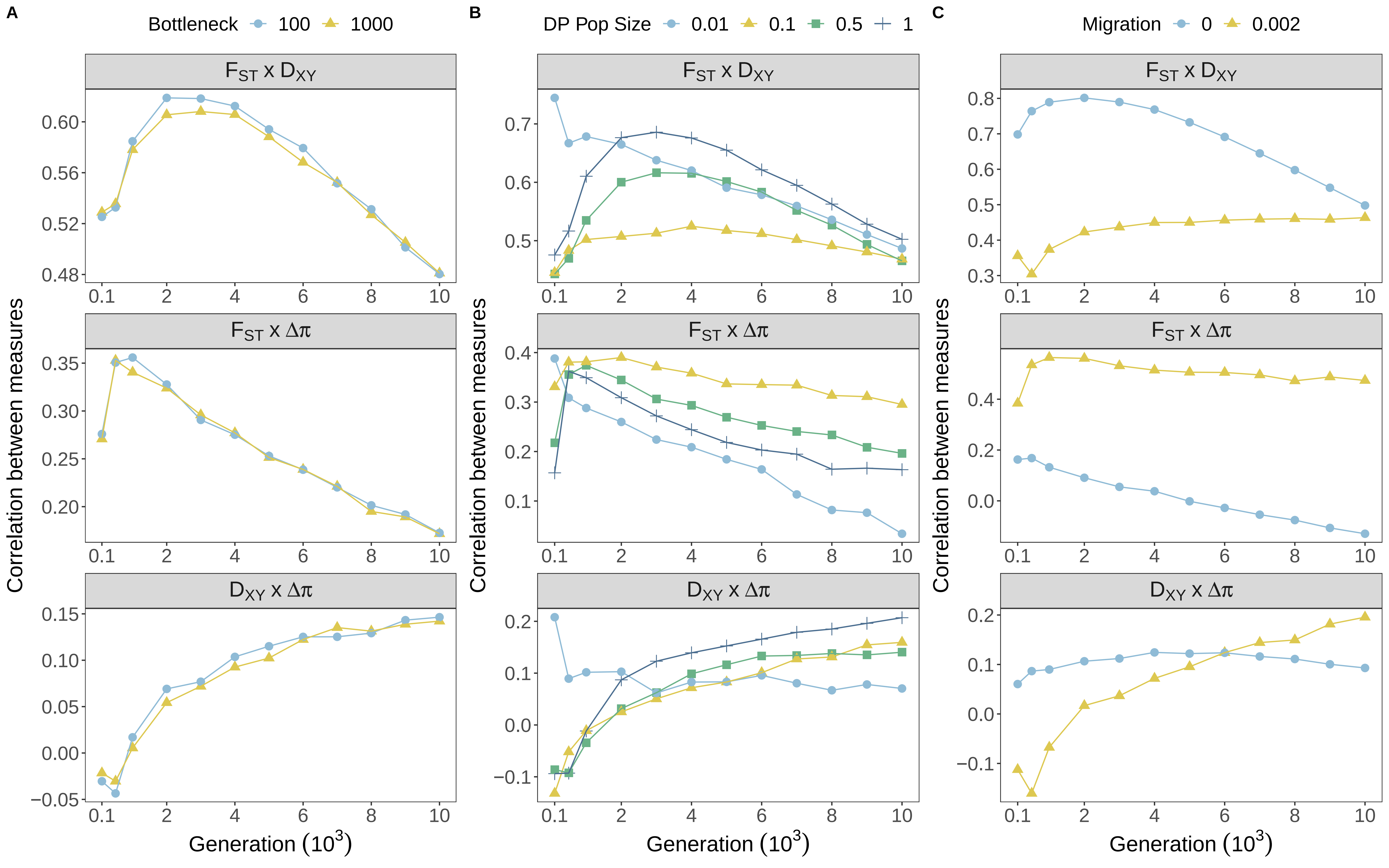

A
Bottleneck 100 1000
B
DP Pop Size 0.01 0.1 0.5 1
C
Migration 0 0.002
$F_{ST}$ x $D_{XY}$
$F_{ST}$ x $\Delta\pi$
$D_{XY}$ x $\Delta\pi$
Correlation between measures
Generation ($10^3$)

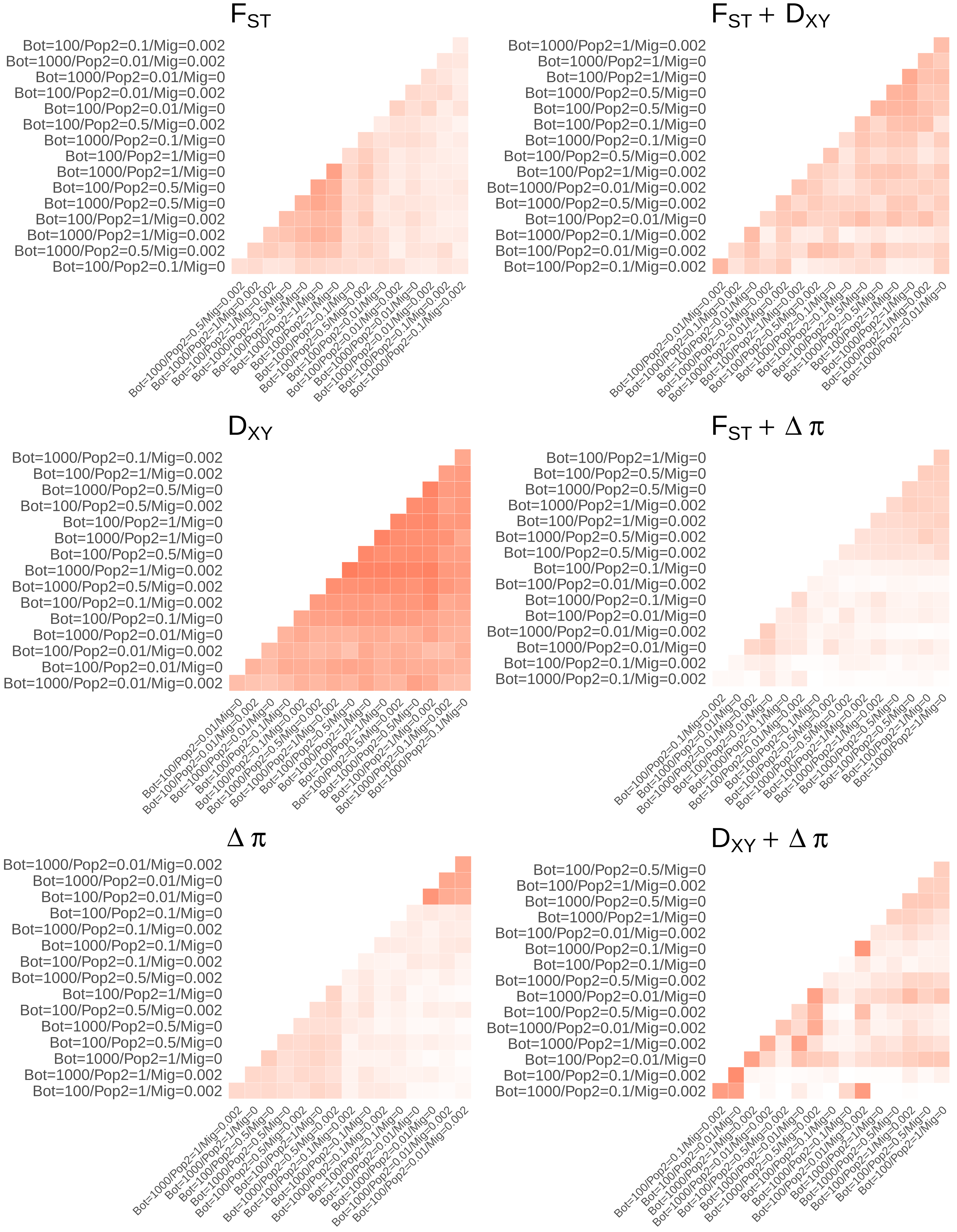

$F_{ST}$
Bot=100/Pop2=0.1/Mig=0.002
Bot=1000/Pop2=0.01/Mig=0.002
Bot=1000/Pop2=0.01/Mig=0
Bot=100/Pop2=0.01/Mig=0.002
Bot=100/Pop2=0.01/Mig=0
Bot=100/Pop2=0.5/Mig=0.002
Bot=1000/Pop2=0.1/Mig=0
Bot=100/Pop2=1/Mig=0
Bot=1000/Pop2=1/Mig=0
Bot=100/Pop2=0.5/Mig=0
Bot=1000/Pop2=0.5/Mig=0
Bot=100/Pop2=1/Mig=0.002
Bot=1000/Pop2=1/Mig=0.002
Bot=1000/Pop2=0.5/Mig=0.002
Bot=100/Pop2=0.1/Mig=0
$F_{ST} + D_{XY}$
Bot=1000/Pop2=1/Mig=0.002
Bot=1000/Pop2=1/Mig=0
Bot=100/Pop2=1/Mig=0
Bot=1000/Pop2=0.5/Mig=0
Bot=100/Pop2=0.5/Mig=0
Bot=100/Pop2=0.1/Mig=0
Bot=1000/Pop2=0.1/Mig=0
Bot=100/Pop2=0.5/Mig=0.002
Bot=100/Pop2=1/Mig=0.002
Bot=1000/Pop2=0.01/Mig=0.002
Bot=1000/Pop2=0.5/Mig=0.002
Bot=100/Pop2=0.01/Mig=0
Bot=1000/Pop2=0.1/Mig=0.002
Bot=100/Pop2=0.01/Mig=0.002
Bot=100/Pop2=0.1/Mig=0.002
$D_{XY}$
Bot=1000/Pop2=0.1/Mig=0.002
Bot=100/Pop2=1/Mig=0.002
Bot=1000/Pop2=0.5/Mig=0
Bot=100/Pop2=0.5/Mig=0.002
Bot=100/Pop2=1/Mig=0
Bot=1000/Pop2=1/Mig=0
Bot=100/Pop2=0.5/Mig=0
Bot=1000/Pop2=1/Mig=0.002
Bot=1000/Pop2=0.5/Mig=0.002
Bot=100/Pop2=0.1/Mig=0.002
Bot=100/Pop2=0.1/Mig=0
Bot=1000/Pop2=0.01/Mig=0
Bot=100/Pop2=0.01/Mig=0.002
Bot=100/Pop2=0.01/Mig=0
Bot=1000/Pop2=0.01/Mig=0.002
$F_{ST} + \Delta\pi$
Bot=100/Pop2=1/Mig=0
Bot=100/Pop2=0.5/Mig=0
Bot=1000/Pop2=0.5/Mig=0
Bot=1000/Pop2=1/Mig=0.002
Bot=100/Pop2=1/Mig=0.002
Bot=1000/Pop2=0.5/Mig=0.002
Bot=100/Pop2=0.5/Mig=0.002
Bot=100/Pop2=0.1/Mig=0
Bot=100/Pop2=0.01/Mig=0.002
Bot=1000/Pop2=0.1/Mig=0
Bot=100/Pop2=0.01/Mig=0
Bot=1000/Pop2=0.01/Mig=0.002
Bot=1000/Pop2=0.01/Mig=0
Bot=100/Pop2=0.1/Mig=0.002
Bot=1000/Pop2=0.1/Mig=0.002
$\Delta\pi$
Bot=1000/Pop2=0.01/Mig=0.002
Bot=1000/Pop2=0.01/Mig=0
Bot=100/Pop2=0.01/Mig=0
Bot=100/Pop2=0.1/Mig=0
Bot=1000/Pop2=0.1/Mig=0.002
Bot=1000/Pop2=0.1/Mig=0
Bot=100/Pop2=0.1/Mig=0.002
Bot=1000/Pop2=0.5/Mig=0.002
Bot=100/Pop2=1/Mig=0
Bot=100/Pop2=0.5/Mig=0.002
Bot=1000/Pop2=0.5/Mig=0
Bot=100/Pop2=0.5/Mig=0
Bot=1000/Pop2=1/Mig=0
Bot=1000/Pop2=1/Mig=0.002
Bot=100/Pop2=1/Mig=0.002
$D_{XY} + \Delta\pi$
Bot=100/Pop2=0.5/Mig=0
Bot=100/Pop2=1/Mig=0.002
Bot=1000/Pop2=0.5/Mig=0
Bot=1000/Pop2=1/Mig=0
Bot=100/Pop2=0.01/Mig=0.002
Bot=1000/Pop2=0.1/Mig=0
Bot=100/Pop2=0.1/Mig=0
Bot=1000/Pop2=0.5/Mig=0.002
Bot=1000/Pop2=0.01/Mig=0
Bot=100/Pop2=0.5/Mig=0.002
Bot=1000/Pop2=0.01/Mig=0.002
Bot=1000/Pop2=1/Mig=0.002
Bot=100/Pop2=0.01/Mig=0
Bot=100/Pop2=0.1/Mig=0.002
Bot=1000/Pop2=0.1/Mig=0.002

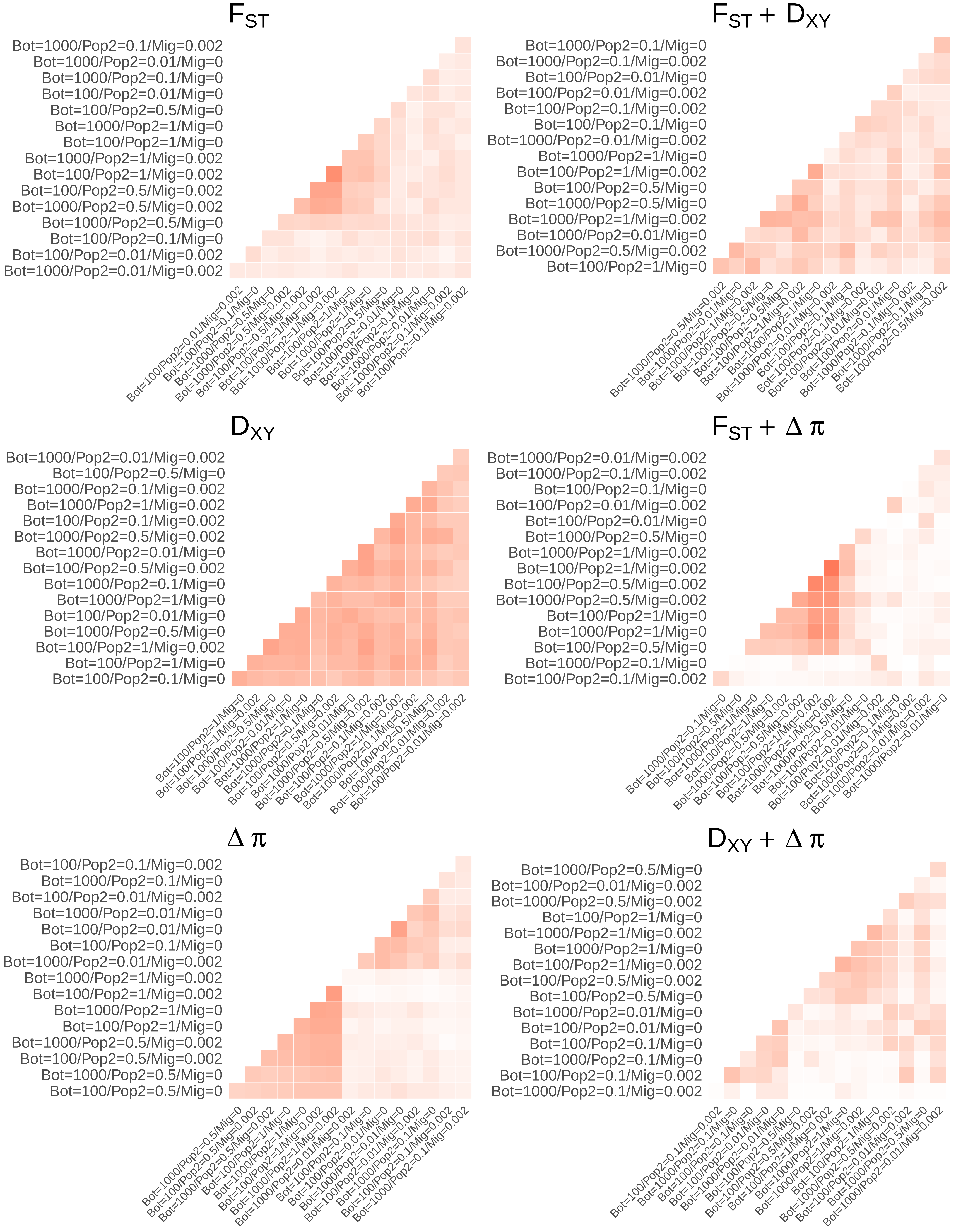

$F_{ST}$
Bot=1000/Pop2=0.1/Mig=0.002
Bot=1000/Pop2=0.01/Mig=0
Bot=1000/Pop2=0.1/Mig=0
Bot=100/Pop2=0.01/Mig=0
Bot=100/Pop2=0.5/Mig=0
Bot=1000/Pop2=1/Mig=0
Bot=100/Pop2=1/Mig=0
Bot=1000/Pop2=1/Mig=0.002
Bot=100/Pop2=1/Mig=0.002
Bot=100/Pop2=0.5/Mig=0.002
Bot=1000/Pop2=0.5/Mig=0.002
Bot=1000/Pop2=0.5/Mig=0
Bot=100/Pop2=0.1/Mig=0
Bot=100/Pop2=0.01/Mig=0.002
Bot=1000/Pop2=0.01/Mig=0.002
$F_{ST} + D_{XY}$
Bot=1000/Pop2=0.1/Mig=0
Bot=1000/Pop2=0.1/Mig=0.002
Bot=100/Pop2=0.01/Mig=0
Bot=100/Pop2=0.01/Mig=0.002
Bot=100/Pop2=0.1/Mig=0.002
Bot=100/Pop2=0.1/Mig=0
Bot=1000/Pop2=0.01/Mig=0.002
Bot=1000/Pop2=1/Mig=0
Bot=100/Pop2=1/Mig=0.002
Bot=100/Pop2=0.5/Mig=0
Bot=1000/Pop2=0.5/Mig=0
Bot=1000/Pop2=1/Mig=0.002
Bot=1000/Pop2=0.01/Mig=0
Bot=1000/Pop2=0.5/Mig=0.002
Bot=100/Pop2=1/Mig=0
$D_{XY}$
Bot=1000/Pop2=0.01/Mig=0.002
Bot=100/Pop2=0.5/Mig=0
Bot=1000/Pop2=0.1/Mig=0.002
Bot=1000/Pop2=1/Mig=0.002
Bot=100/Pop2=0.1/Mig=0.002
Bot=1000/Pop2=0.5/Mig=0.002
Bot=1000/Pop2=0.01/Mig=0
Bot=100/Pop2=0.5/Mig=0.002
Bot=1000/Pop2=0.1/Mig=0
Bot=1000/Pop2=1/Mig=0
Bot=100/Pop2=0.01/Mig=0
Bot=1000/Pop2=0.5/Mig=0
Bot=100/Pop2=1/Mig=0.002
Bot=100/Pop2=1/Mig=0
Bot=100/Pop2=0.1/Mig=0
$F_{ST} + \Delta\pi$
Bot=1000/Pop2=0.01/Mig=0.002
Bot=1000/Pop2=0.1/Mig=0.002
Bot=100/Pop2=0.1/Mig=0
Bot=100/Pop2=0.01/Mig=0.002
Bot=100/Pop2=0.01/Mig=0
Bot=1000/Pop2=0.5/Mig=0
Bot=1000/Pop2=1/Mig=0.002
Bot=100/Pop2=1/Mig=0.002
Bot=100/Pop2=0.5/Mig=0.002
Bot=1000/Pop2=0.5/Mig=0.002
Bot=100/Pop2=1/Mig=0
Bot=1000/Pop2=1/Mig=0
Bot=100/Pop2=0.5/Mig=0
Bot=1000/Pop2=0.1/Mig=0
Bot=100/Pop2=0.1/Mig=0.002
$\Delta\pi$
Bot=100/Pop2=0.1/Mig=0.002
Bot=1000/Pop2=0.1/Mig=0
Bot=100/Pop2=0.01/Mig=0.002
Bot=1000/Pop2=0.01/Mig=0
Bot=100/Pop2=0.01/Mig=0
Bot=100/Pop2=0.1/Mig=0
Bot=1000/Pop2=0.01/Mig=0.002
Bot=1000/Pop2=1/Mig=0.002
Bot=100/Pop2=1/Mig=0.002
Bot=1000/Pop2=1/Mig=0
Bot=100/Pop2=1/Mig=0
Bot=1000/Pop2=0.5/Mig=0.002
Bot=100/Pop2=0.5/Mig=0.002
Bot=1000/Pop2=0.5/Mig=0
Bot=100/Pop2=0.5/Mig=0
$D_{XY} + \Delta\pi$
Bot=1000/Pop2=0.5/Mig=0
Bot=100/Pop2=0.01/Mig=0.002
Bot=1000/Pop2=0.5/Mig=0.002
Bot=100/Pop2=1/Mig=0
Bot=1000/Pop2=1/Mig=0.002
Bot=1000/Pop2=1/Mig=0
Bot=100/Pop2=1/Mig=0.002
Bot=100/Pop2=0.5/Mig=0.002
Bot=100/Pop2=0.5/Mig=0
Bot=1000/Pop2=0.01/Mig=0
Bot=100/Pop2=0.01/Mig=0
Bot=100/Pop2=0.1/Mig=0
Bot=1000/Pop2=0.1/Mig=0
Bot=100/Pop2=0.1/Mig=0.002
Bot=1000/Pop2=0.1/Mig=0.002

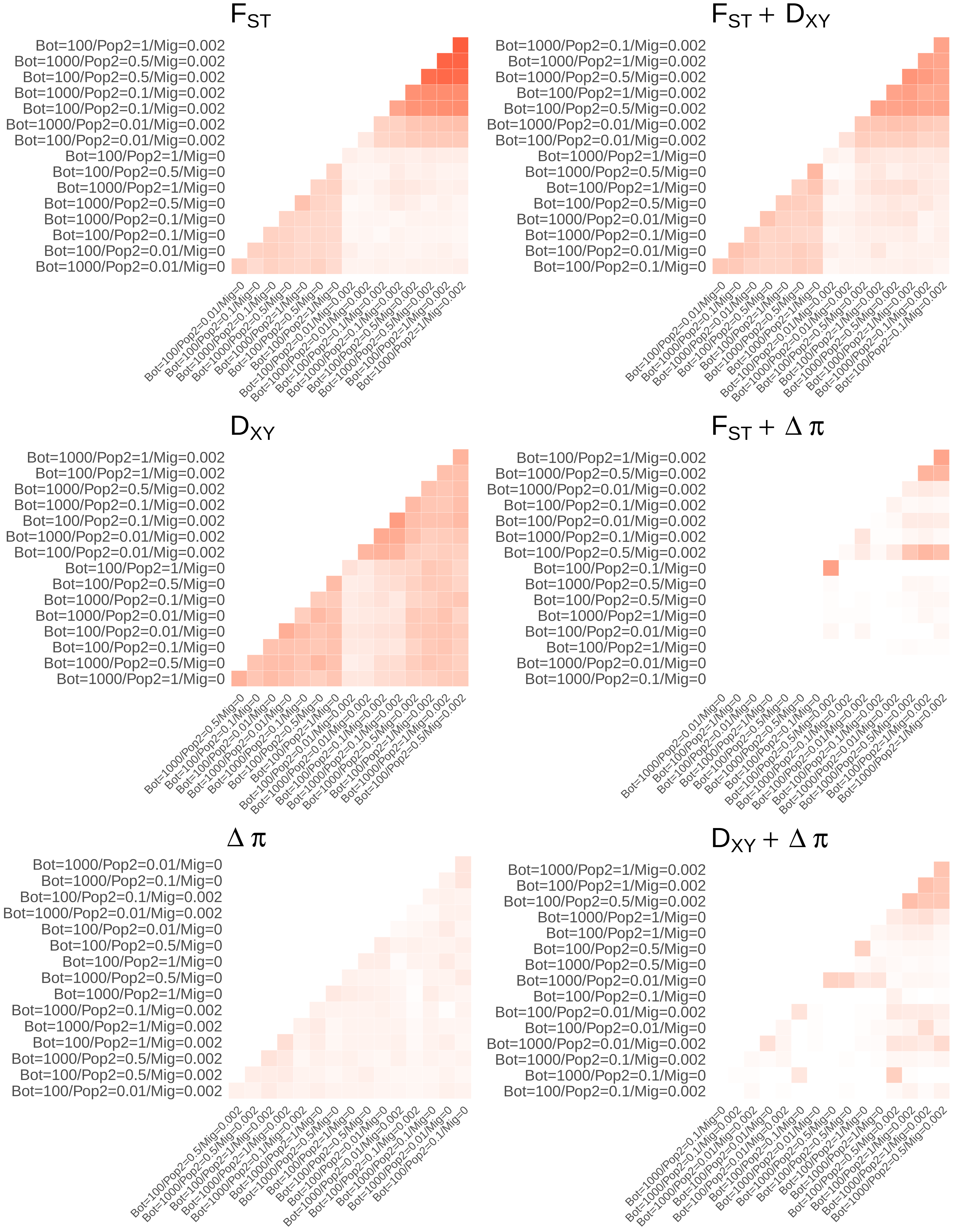
$F_{ST}$
Bot=100/Pop2=1/Mig=0.002
Bot=1000/Pop2=0.5/Mig=0.002
Bot=100/Pop2=0.5/Mig=0.002
Bot=1000/Pop2=0.1/Mig=0.002
Bot=100/Pop2=0.1/Mig=0.002
Bot=1000/Pop2=0.01/Mig=0.002
Bot=100/Pop2=0.01/Mig=0.002
Bot=100/Pop2=1/Mig=0
Bot=100/Pop2=0.5/Mig=0
Bot=1000/Pop2=1/Mig=0
Bot=1000/Pop2=0.5/Mig=0
Bot=1000/Pop2=0.1/Mig=0
Bot=100/Pop2=0.1/Mig=0
Bot=100/Pop2=0.01/Mig=0
Bot=1000/Pop2=0.01/Mig=0
$F_{ST}$ + $D_{XY}$
Bot=1000/Pop2=0.1/Mig=0.002
Bot=1000/Pop2=1/Mig=0.002
Bot=1000/Pop2=0.5/Mig=0.002
Bot=100/Pop2=1/Mig=0.002
Bot=100/Pop2=0.5/Mig=0.002
Bot=1000/Pop2=0.01/Mig=0.002
Bot=100/Pop2=0.01/Mig=0.002
Bot=1000/Pop2=1/Mig=0
Bot=1000/Pop2=0.5/Mig=0
Bot=100/Pop2=1/Mig=0
Bot=100/Pop2=0.5/Mig=0
Bot=1000/Pop2=0.01/Mig=0
Bot=1000/Pop2=0.1/Mig=0
Bot=100/Pop2=0.01/Mig=0
Bot=100/Pop2=0.1/Mig=0
$D_{XY}$
Bot=1000/Pop2=1/Mig=0.002
Bot=100/Pop2=1/Mig=0.002
Bot=1000/Pop2=0.5/Mig=0.002
Bot=1000/Pop2=0.1/Mig=0.002
Bot=100/Pop2=0.1/Mig=0.002
Bot=1000/Pop2=0.01/Mig=0.002
Bot=100/Pop2=0.01/Mig=0.002
Bot=100/Pop2=1/Mig=0
Bot=100/Pop2=0.5/Mig=0
Bot=1000/Pop2=0.1/Mig=0
Bot=1000/Pop2=0.01/Mig=0
Bot=100/Pop2=0.01/Mig=0
Bot=100/Pop2=0.1/Mig=0
Bot=1000/Pop2=0.5/Mig=0
Bot=1000/Pop2=1/Mig=0
$F_{ST}$ + $\Delta\pi$
Bot=100/Pop2=1/Mig=0.002
Bot=1000/Pop2=0.5/Mig=0.002
Bot=1000/Pop2=0.01/Mig=0.002
Bot=100/Pop2=0.1/Mig=0.002
Bot=100/Pop2=0.01/Mig=0.002
Bot=1000/Pop2=0.1/Mig=0.002
Bot=100/Pop2=0.5/Mig=0.002
Bot=100/Pop2=0.1/Mig=0
Bot=1000/Pop2=0.5/Mig=0
Bot=100/Pop2=0.5/Mig=0
Bot=1000/Pop2=1/Mig=0
Bot=100/Pop2=0.01/Mig=0
Bot=100/Pop2=1/Mig=0
Bot=1000/Pop2=0.01/Mig=0
Bot=1000/Pop2=0.1/Mig=0
$\Delta\pi$
Bot=1000/Pop2=0.01/Mig=0
Bot=1000/Pop2=0.1/Mig=0
Bot=100/Pop2=0.1/Mig=0.002
Bot=1000/Pop2=0.01/Mig=0.002
Bot=100/Pop2=0.01/Mig=0
Bot=100/Pop2=0.5/Mig=0
Bot=100/Pop2=1/Mig=0
Bot=1000/Pop2=0.5/Mig=0
Bot=1000/Pop2=1/Mig=0
Bot=1000/Pop2=0.1/Mig=0.002
Bot=1000/Pop2=1/Mig=0.002
Bot=100/Pop2=1/Mig=0.002
Bot=1000/Pop2=0.5/Mig=0.002
Bot=100/Pop2=0.5/Mig=0.002
Bot=100/Pop2=0.01/Mig=0.002
$D_{XY}$ + $\Delta\pi$
Bot=1000/Pop2=1/Mig=0.002
Bot=100/Pop2=1/Mig=0.002
Bot=100/Pop2=0.5/Mig=0.002
Bot=1000/Pop2=1/Mig=0
Bot=100/Pop2=1/Mig=0
Bot=100/Pop2=0.5/Mig=0
Bot=1000/Pop2=0.5/Mig=0
Bot=1000/Pop2=0.01/Mig=0
Bot=100/Pop2=0.1/Mig=0
Bot=100/Pop2=0.01/Mig=0.002
Bot=100/Pop2=0.01/Mig=0
Bot=1000/Pop2=0.01/Mig=0.002
Bot=1000/Pop2=0.1/Mig=0.002
Bot=1000/Pop2=0.1/Mig=0
Bot=100/Pop2=0.1/Mig=0.002

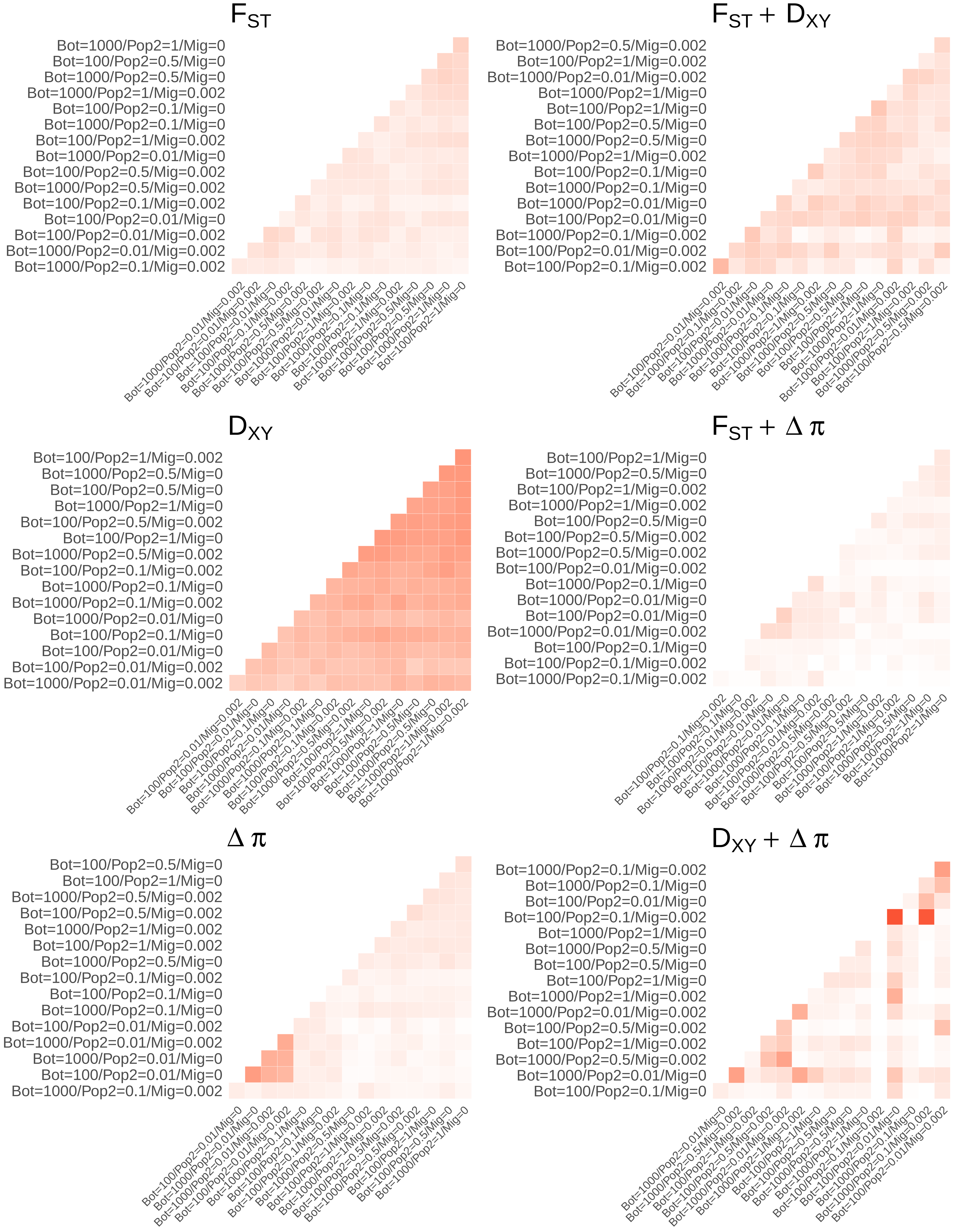
$F_{ST}$
Bot=1000/Pop2=1/Mig=0
Bot=100/Pop2=0.5/Mig=0
Bot=1000/Pop2=0.5/Mig=0
Bot=1000/Pop2=1/Mig=0.002
Bot=100/Pop2=0.1/Mig=0
Bot=1000/Pop2=0.1/Mig=0
Bot=100/Pop2=1/Mig=0.002
Bot=1000/Pop2=0.01/Mig=0
Bot=100/Pop2=0.5/Mig=0.002
Bot=1000/Pop2=0.5/Mig=0.002
Bot=100/Pop2=0.1/Mig=0.002
Bot=100/Pop2=0.01/Mig=0
Bot=100/Pop2=0.01/Mig=0.002
Bot=1000/Pop2=0.01/Mig=0.002
Bot=1000/Pop2=0.1/Mig=0.002
$F_{ST} + D_{XY}$
Bot=1000/Pop2=0.5/Mig=0.002
Bot=100/Pop2=1/Mig=0.002
Bot=1000/Pop2=0.01/Mig=0.002
Bot=1000/Pop2=1/Mig=0
Bot=100/Pop2=1/Mig=0
Bot=100/Pop2=0.5/Mig=0
Bot=1000/Pop2=0.5/Mig=0
Bot=1000/Pop2=1/Mig=0.002
Bot=100/Pop2=0.1/Mig=0
Bot=1000/Pop2=0.1/Mig=0
Bot=1000/Pop2=0.01/Mig=0
Bot=100/Pop2=0.01/Mig=0
Bot=1000/Pop2=0.1/Mig=0.002
Bot=100/Pop2=0.01/Mig=0.002
Bot=100/Pop2=0.1/Mig=0.002
$D_{XY}$
Bot=100/Pop2=1/Mig=0.002
Bot=1000/Pop2=0.5/Mig=0
Bot=100/Pop2=0.5/Mig=0
Bot=1000/Pop2=1/Mig=0
Bot=100/Pop2=0.5/Mig=0.002
Bot=100/Pop2=1/Mig=0
Bot=1000/Pop2=0.5/Mig=0.002
Bot=100/Pop2=0.1/Mig=0.002
Bot=1000/Pop2=0.1/Mig=0
Bot=1000/Pop2=0.1/Mig=0.002
Bot=1000/Pop2=0.01/Mig=0
Bot=100/Pop2=0.1/Mig=0
Bot=100/Pop2=0.01/Mig=0
Bot=100/Pop2=0.01/Mig=0.002
Bot=1000/Pop2=0.01/Mig=0.002
$F_{ST} + \Delta\pi$
Bot=100/Pop2=1/Mig=0
Bot=1000/Pop2=0.5/Mig=0
Bot=100/Pop2=1/Mig=0.002
Bot=1000/Pop2=1/Mig=0.002
Bot=100/Pop2=0.5/Mig=0
Bot=100/Pop2=0.5/Mig=0.002
Bot=1000/Pop2=0.5/Mig=0.002
Bot=100/Pop2=0.01/Mig=0.002
Bot=1000/Pop2=0.1/Mig=0
Bot=1000/Pop2=0.01/Mig=0
Bot=100/Pop2=0.01/Mig=0
Bot=1000/Pop2=0.01/Mig=0.002
Bot=100/Pop2=0.1/Mig=0
Bot=100/Pop2=0.1/Mig=0.002
Bot=1000/Pop2=0.1/Mig=0.002
$\Delta\pi$
Bot=100/Pop2=0.5/Mig=0
Bot=100/Pop2=1/Mig=0
Bot=1000/Pop2=0.5/Mig=0.002
Bot=100/Pop2=0.5/Mig=0.002
Bot=1000/Pop2=1/Mig=0.002
Bot=100/Pop2=1/Mig=0.002
Bot=1000/Pop2=0.5/Mig=0
Bot=100/Pop2=0.1/Mig=0.002
Bot=100/Pop2=0.1/Mig=0
Bot=1000/Pop2=0.1/Mig=0
Bot=100/Pop2=0.01/Mig=0.002
Bot=1000/Pop2=0.01/Mig=0.002
Bot=1000/Pop2=0.01/Mig=0
Bot=100/Pop2=0.01/Mig=0
Bot=1000/Pop2=0.1/Mig=0.002
$D_{XY} + \Delta\pi$
Bot=1000/Pop2=0.1/Mig=0.002
Bot=1000/Pop2=0.1/Mig=0
Bot=100/Pop2=0.01/Mig=0
Bot=100/Pop2=0.1/Mig=0.002
Bot=1000/Pop2=1/Mig=0
Bot=1000/Pop2=0.5/Mig=0
Bot=100/Pop2=0.5/Mig=0
Bot=100/Pop2=1/Mig=0
Bot=1000/Pop2=1/Mig=0.002
Bot=1000/Pop2=0.01/Mig=0.002
Bot=100/Pop2=0.5/Mig=0.002
Bot=100/Pop2=1/Mig=0.002
Bot=1000/Pop2=0.5/Mig=0.002
Bot=1000/Pop2=0.01/Mig=0
Bot=100/Pop2=0.1/Mig=0

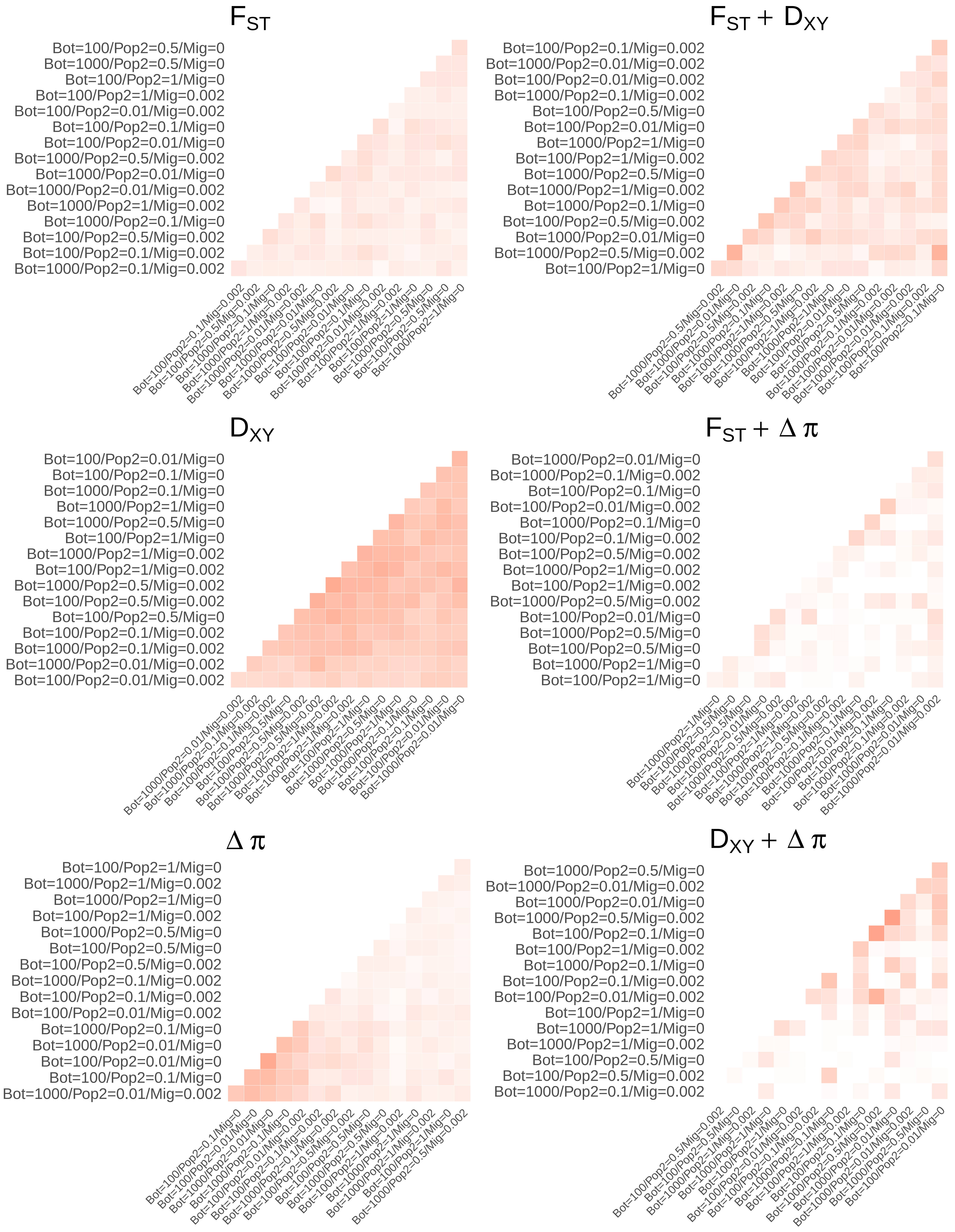

$F_{ST}$
Bot=100/Pop2=0.5/Mig=0
Bot=1000/Pop2=0.5/Mig=0
Bot=100/Pop2=1/Mig=0
Bot=100/Pop2=1/Mig=0.002
Bot=100/Pop2=0.01/Mig=0.002
Bot=100/Pop2=0.1/Mig=0
Bot=100/Pop2=0.01/Mig=0
Bot=1000/Pop2=0.5/Mig=0.002
Bot=1000/Pop2=0.01/Mig=0
Bot=1000/Pop2=0.01/Mig=0.002
Bot=1000/Pop2=1/Mig=0.002
Bot=1000/Pop2=0.1/Mig=0
Bot=100/Pop2=0.5/Mig=0.002
Bot=100/Pop2=0.1/Mig=0.002
Bot=1000/Pop2=0.1/Mig=0.002
$F_{ST} + D_{XY}$
Bot=100/Pop2=0.1/Mig=0.002
Bot=1000/Pop2=0.01/Mig=0.002
Bot=100/Pop2=0.01/Mig=0.002
Bot=1000/Pop2=0.1/Mig=0.002
Bot=100/Pop2=0.5/Mig=0
Bot=100/Pop2=0.01/Mig=0
Bot=1000/Pop2=1/Mig=0
Bot=100/Pop2=1/Mig=0.002
Bot=1000/Pop2=0.5/Mig=0
Bot=1000/Pop2=1/Mig=0.002
Bot=1000/Pop2=0.1/Mig=0
Bot=100/Pop2=0.5/Mig=0.002
Bot=1000/Pop2=0.01/Mig=0
Bot=1000/Pop2=0.5/Mig=0.002
Bot=100/Pop2=1/Mig=0
$D_{XY}$
Bot=100/Pop2=0.01/Mig=0
Bot=100/Pop2=0.1/Mig=0
Bot=1000/Pop2=0.1/Mig=0
Bot=1000/Pop2=1/Mig=0
Bot=1000/Pop2=0.5/Mig=0
Bot=100/Pop2=1/Mig=0
Bot=1000/Pop2=1/Mig=0.002
Bot=100/Pop2=1/Mig=0.002
Bot=1000/Pop2=0.5/Mig=0.002
Bot=100/Pop2=0.5/Mig=0.002
Bot=100/Pop2=0.5/Mig=0
Bot=100/Pop2=0.1/Mig=0.002
Bot=1000/Pop2=0.1/Mig=0.002
Bot=1000/Pop2=0.01/Mig=0.002
Bot=100/Pop2=0.01/Mig=0.002
$F_{ST} + \Delta\pi$
Bot=1000/Pop2=0.01/Mig=0
Bot=1000/Pop2=0.1/Mig=0.002
Bot=100/Pop2=0.1/Mig=0
Bot=100/Pop2=0.01/Mig=0.002
Bot=1000/Pop2=0.1/Mig=0
Bot=100/Pop2=0.1/Mig=0.002
Bot=100/Pop2=0.5/Mig=0.002
Bot=1000/Pop2=1/Mig=0.002
Bot=100/Pop2=1/Mig=0.002
Bot=1000/Pop2=0.5/Mig=0.002
Bot=100/Pop2=0.01/Mig=0
Bot=1000/Pop2=0.5/Mig=0
Bot=100/Pop2=0.5/Mig=0
Bot=1000/Pop2=1/Mig=0
Bot=100/Pop2=1/Mig=0
$\Delta\pi$
Bot=100/Pop2=1/Mig=0
Bot=1000/Pop2=1/Mig=0.002
Bot=1000/Pop2=1/Mig=0
Bot=100/Pop2=1/Mig=0.002
Bot=1000/Pop2=0.5/Mig=0
Bot=100/Pop2=0.5/Mig=0
Bot=100/Pop2=0.5/Mig=0.002
Bot=1000/Pop2=0.1/Mig=0.002
Bot=100/Pop2=0.1/Mig=0.002
Bot=100/Pop2=0.01/Mig=0.002
Bot=1000/Pop2=0.1/Mig=0
Bot=1000/Pop2=0.01/Mig=0
Bot=100/Pop2=0.01/Mig=0
Bot=100/Pop2=0.1/Mig=0
Bot=1000/Pop2=0.01/Mig=0.002
$D_{XY} + \Delta\pi$
Bot=1000/Pop2=0.5/Mig=0
Bot=1000/Pop2=0.01/Mig=0.002
Bot=1000/Pop2=0.01/Mig=0
Bot=1000/Pop2=0.5/Mig=0.002
Bot=100/Pop2=0.1/Mig=0
Bot=100/Pop2=1/Mig=0.002
Bot=1000/Pop2=0.1/Mig=0
Bot=100/Pop2=0.1/Mig=0.002
Bot=100/Pop2=0.01/Mig=0.002
Bot=100/Pop2=1/Mig=0
Bot=1000/Pop2=1/Mig=0
Bot=1000/Pop2=1/Mig=0.002
Bot=100/Pop2=0.5/Mig=0
Bot=100/Pop2=0.5/Mig=0.002
Bot=1000/Pop2=0.1/Mig=0.002

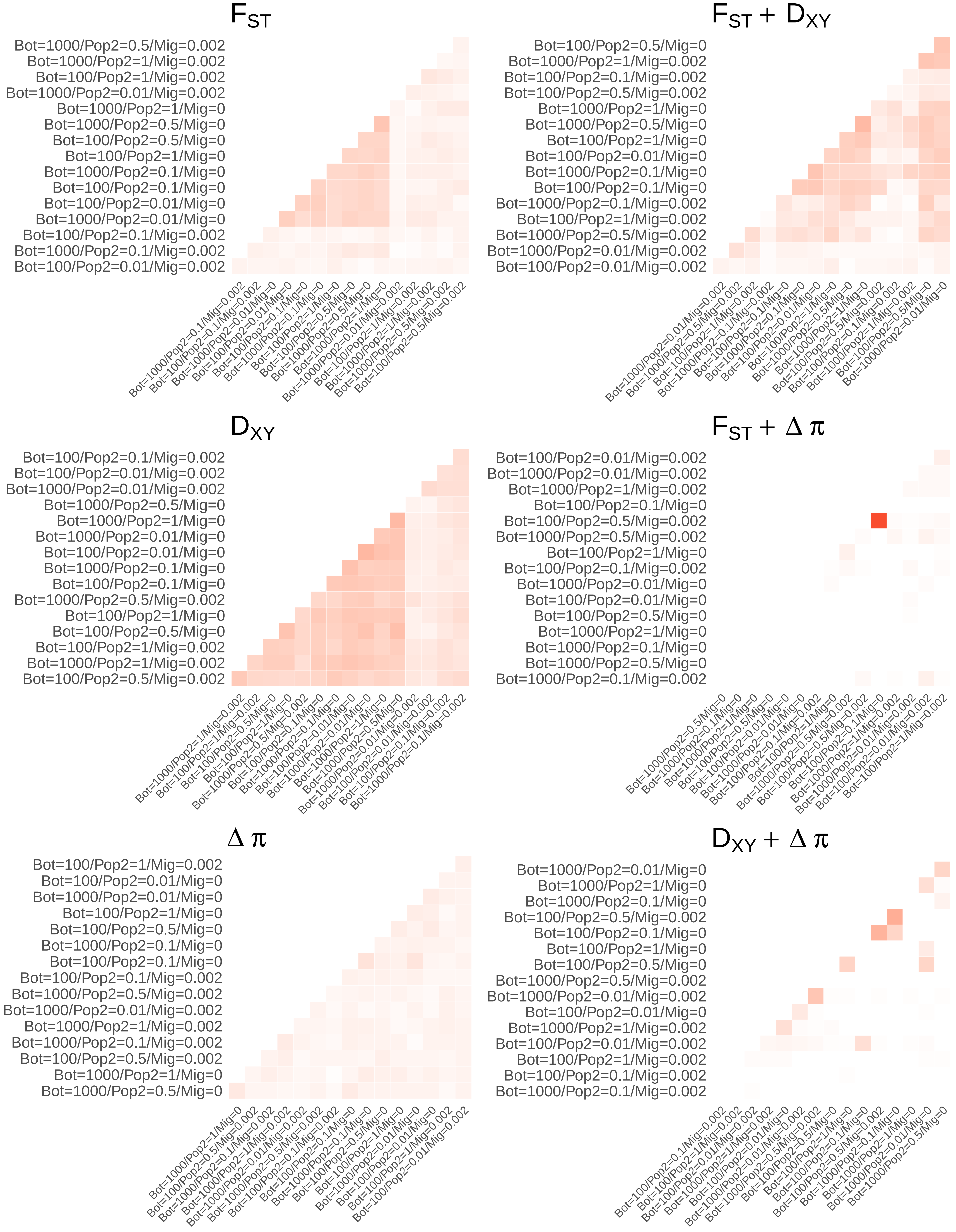

F_ST
Bot=1000/Pop2=0.5/Mig=0.002
Bot=1000/Pop2=1/Mig=0.002
Bot=100/Pop2=1/Mig=0.002
Bot=1000/Pop2=0.01/Mig=0.002
Bot=1000/Pop2=1/Mig=0
Bot=1000/Pop2=0.5/Mig=0
Bot=100/Pop2=0.5/Mig=0
Bot=100/Pop2=1/Mig=0
Bot=1000/Pop2=0.1/Mig=0
Bot=100/Pop2=0.1/Mig=0
Bot=100/Pop2=0.01/Mig=0
Bot=1000/Pop2=0.01/Mig=0
Bot=100/Pop2=0.1/Mig=0.002
Bot=1000/Pop2=0.1/Mig=0.002
Bot=100/Pop2=0.01/Mig=0.002
F_ST + D_XY
Bot=100/Pop2=0.5/Mig=0
Bot=1000/Pop2=1/Mig=0.002
Bot=100/Pop2=0.1/Mig=0.002
Bot=100/Pop2=0.5/Mig=0.002
Bot=1000/Pop2=1/Mig=0
Bot=1000/Pop2=0.5/Mig=0
Bot=100/Pop2=1/Mig=0
Bot=100/Pop2=0.01/Mig=0
Bot=1000/Pop2=0.1/Mig=0
Bot=100/Pop2=0.1/Mig=0
Bot=1000/Pop2=0.1/Mig=0.002
Bot=100/Pop2=1/Mig=0.002
Bot=1000/Pop2=0.5/Mig=0.002
Bot=1000/Pop2=0.01/Mig=0.002
Bot=100/Pop2=0.01/Mig=0.002
D_XY
Bot=100/Pop2=0.1/Mig=0.002
Bot=100/Pop2=0.01/Mig=0.002
Bot=1000/Pop2=0.01/Mig=0.002
Bot=1000/Pop2=0.5/Mig=0
Bot=1000/Pop2=1/Mig=0
Bot=1000/Pop2=0.01/Mig=0
Bot=100/Pop2=0.01/Mig=0
Bot=1000/Pop2=0.1/Mig=0
Bot=100/Pop2=0.1/Mig=0
Bot=1000/Pop2=0.5/Mig=0.002
Bot=100/Pop2=1/Mig=0
Bot=100/Pop2=0.5/Mig=0
Bot=100/Pop2=1/Mig=0.002
Bot=1000/Pop2=1/Mig=0.002
Bot=100/Pop2=0.5/Mig=0.002
F_ST + Δ π
Bot=100/Pop2=0.01/Mig=0.002
Bot=1000/Pop2=0.01/Mig=0.002
Bot=1000/Pop2=1/Mig=0.002
Bot=100/Pop2=0.1/Mig=0
Bot=100/Pop2=0.5/Mig=0.002
Bot=1000/Pop2=0.5/Mig=0.002
Bot=100/Pop2=1/Mig=0
Bot=100/Pop2=0.1/Mig=0.002
Bot=1000/Pop2=0.01/Mig=0
Bot=100/Pop2=0.01/Mig=0
Bot=100/Pop2=0.5/Mig=0
Bot=1000/Pop2=1/Mig=0
Bot=1000/Pop2=0.1/Mig=0
Bot=1000/Pop2=0.5/Mig=0
Bot=1000/Pop2=0.1/Mig=0.002
Δ π
Bot=100/Pop2=1/Mig=0.002
Bot=100/Pop2=0.01/Mig=0
Bot=1000/Pop2=0.01/Mig=0
Bot=100/Pop2=1/Mig=0
Bot=100/Pop2=0.5/Mig=0
Bot=1000/Pop2=0.1/Mig=0
Bot=100/Pop2=0.1/Mig=0
Bot=100/Pop2=0.1/Mig=0.002
Bot=1000/Pop2=0.5/Mig=0.002
Bot=1000/Pop2=0.01/Mig=0.002
Bot=1000/Pop2=1/Mig=0.002
Bot=1000/Pop2=0.1/Mig=0.002
Bot=100/Pop2=0.5/Mig=0.002
Bot=1000/Pop2=1/Mig=0
Bot=1000/Pop2=0.5/Mig=0
D_XY + Δ π
Bot=1000/Pop2=0.01/Mig=0
Bot=1000/Pop2=1/Mig=0
Bot=1000/Pop2=0.1/Mig=0
Bot=100/Pop2=0.5/Mig=0.002
Bot=100/Pop2=0.1/Mig=0
Bot=100/Pop2=1/Mig=0
Bot=100/Pop2=0.5/Mig=0
Bot=1000/Pop2=0.5/Mig=0.002
Bot=1000/Pop2=0.01/Mig=0.002
Bot=100/Pop2=0.01/Mig=0
Bot=1000/Pop2=1/Mig=0.002
Bot=100/Pop2=0.01/Mig=0.002
Bot=100/Pop2=1/Mig=0.002
Bot=100/Pop2=0.1/Mig=0.002
Bot=1000/Pop2=0.1/Mig=0.002

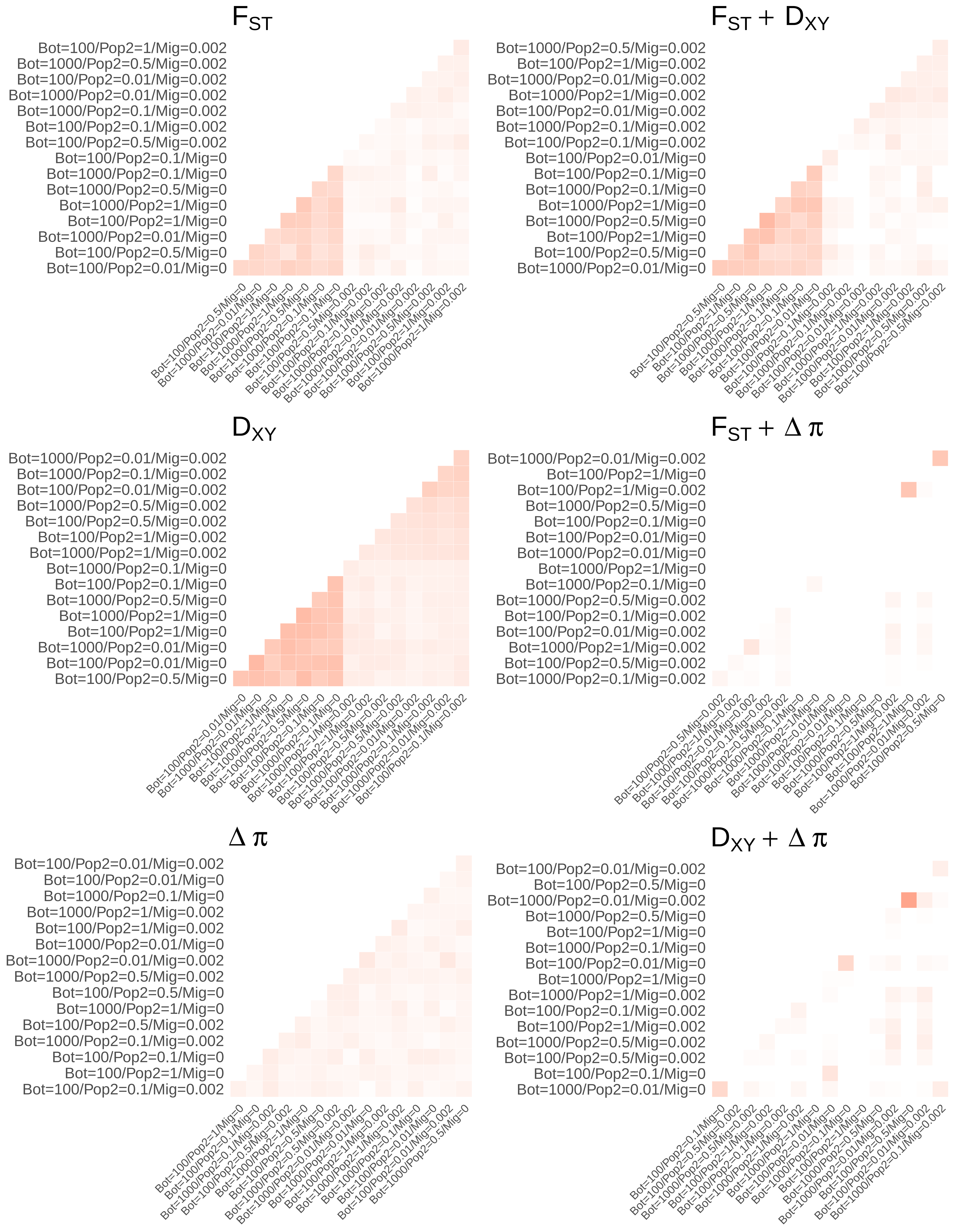
$F_{ST}$
Bot=100/Pop2=1/Mig=0.002
Bot=1000/Pop2=0.5/Mig=0.002
Bot=100/Pop2=0.01/Mig=0.002
Bot=1000/Pop2=0.01/Mig=0.002
Bot=1000/Pop2=0.1/Mig=0.002
Bot=100/Pop2=0.1/Mig=0.002
Bot=100/Pop2=0.5/Mig=0.002
Bot=100/Pop2=0.1/Mig=0
Bot=1000/Pop2=0.1/Mig=0
Bot=1000/Pop2=0.5/Mig=0
Bot=1000/Pop2=1/Mig=0
Bot=100/Pop2=1/Mig=0
Bot=1000/Pop2=0.01/Mig=0
Bot=100/Pop2=0.5/Mig=0
Bot=100/Pop2=0.01/Mig=0
$F_{ST} + D_{XY}$
Bot=1000/Pop2=0.5/Mig=0.002
Bot=100/Pop2=1/Mig=0.002
Bot=1000/Pop2=0.01/Mig=0.002
Bot=1000/Pop2=1/Mig=0.002
Bot=100/Pop2=0.01/Mig=0.002
Bot=1000/Pop2=0.1/Mig=0.002
Bot=100/Pop2=0.1/Mig=0.002
Bot=100/Pop2=0.01/Mig=0
Bot=100/Pop2=0.1/Mig=0
Bot=1000/Pop2=0.1/Mig=0
Bot=1000/Pop2=1/Mig=0
Bot=1000/Pop2=0.5/Mig=0
Bot=100/Pop2=1/Mig=0
Bot=100/Pop2=0.5/Mig=0
Bot=1000/Pop2=0.01/Mig=0
$D_{XY}$
Bot=1000/Pop2=0.01/Mig=0.002
Bot=1000/Pop2=0.1/Mig=0.002
Bot=100/Pop2=0.01/Mig=0.002
Bot=1000/Pop2=0.5/Mig=0.002
Bot=100/Pop2=0.5/Mig=0.002
Bot=100/Pop2=1/Mig=0.002
Bot=1000/Pop2=1/Mig=0.002
Bot=1000/Pop2=0.1/Mig=0
Bot=100/Pop2=0.1/Mig=0
Bot=1000/Pop2=0.5/Mig=0
Bot=1000/Pop2=1/Mig=0
Bot=100/Pop2=1/Mig=0
Bot=1000/Pop2=0.01/Mig=0
Bot=100/Pop2=0.01/Mig=0
Bot=100/Pop2=0.5/Mig=0
$F_{ST} + \Delta\pi$
Bot=1000/Pop2=0.01/Mig=0.002
Bot=100/Pop2=1/Mig=0
Bot=100/Pop2=1/Mig=0.002
Bot=1000/Pop2=0.5/Mig=0
Bot=100/Pop2=0.1/Mig=0
Bot=100/Pop2=0.01/Mig=0
Bot=1000/Pop2=0.01/Mig=0
Bot=1000/Pop2=1/Mig=0
Bot=1000/Pop2=0.1/Mig=0
Bot=1000/Pop2=0.5/Mig=0.002
Bot=100/Pop2=0.1/Mig=0.002
Bot=100/Pop2=0.01/Mig=0.002
Bot=1000/Pop2=1/Mig=0.002
Bot=100/Pop2=0.5/Mig=0.002
Bot=1000/Pop2=0.1/Mig=0.002
$\Delta\pi$
Bot=100/Pop2=0.01/Mig=0.002
Bot=100/Pop2=0.01/Mig=0
Bot=1000/Pop2=0.1/Mig=0
Bot=1000/Pop2=1/Mig=0.002
Bot=100/Pop2=1/Mig=0.002
Bot=1000/Pop2=0.01/Mig=0
Bot=1000/Pop2=0.01/Mig=0.002
Bot=1000/Pop2=0.5/Mig=0.002
Bot=100/Pop2=0.5/Mig=0
Bot=1000/Pop2=1/Mig=0
Bot=100/Pop2=0.5/Mig=0.002
Bot=1000/Pop2=0.1/Mig=0.002
Bot=100/Pop2=0.1/Mig=0
Bot=100/Pop2=1/Mig=0
Bot=100/Pop2=0.1/Mig=0.002
$D_{XY} + \Delta\pi$
Bot=100/Pop2=0.01/Mig=0.002
Bot=100/Pop2=0.5/Mig=0
Bot=1000/Pop2=0.01/Mig=0.002
Bot=1000/Pop2=0.5/Mig=0
Bot=100/Pop2=1/Mig=0
Bot=1000/Pop2=0.1/Mig=0
Bot=100/Pop2=0.01/Mig=0
Bot=1000/Pop2=1/Mig=0
Bot=1000/Pop2=1/Mig=0.002
Bot=100/Pop2=0.1/Mig=0.002
Bot=100/Pop2=1/Mig=0.002
Bot=1000/Pop2=0.5/Mig=0.002
Bot=100/Pop2=0.5/Mig=0.002
Bot=100/Pop2=0.1/Mig=0
Bot=1000/Pop2=0.01/Mig=0

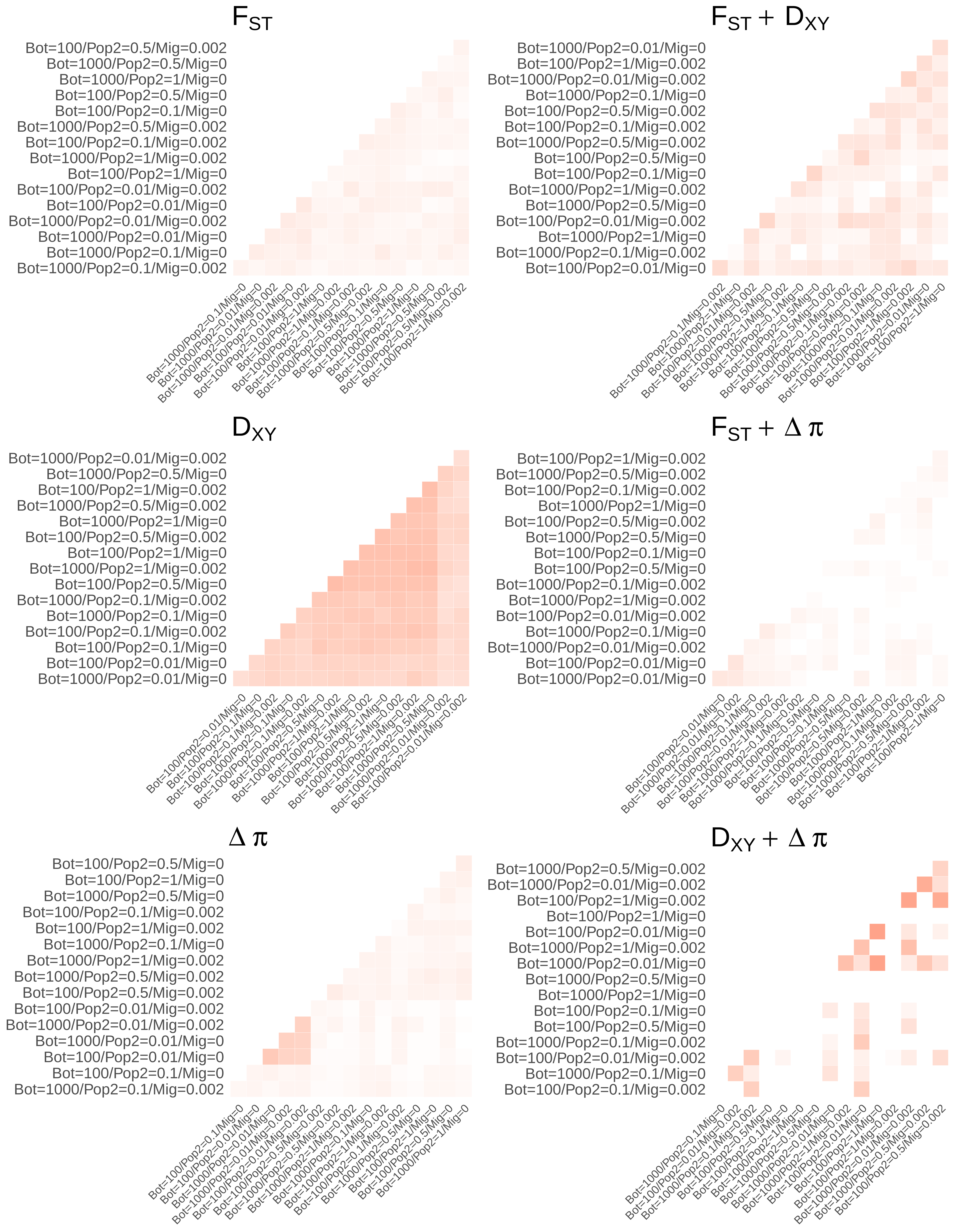

$F_{ST}$
Bot=100/Pop2=0.5/Mig=0.002
Bot=1000/Pop2=0.5/Mig=0
Bot=1000/Pop2=1/Mig=0
Bot=100/Pop2=0.5/Mig=0
Bot=100/Pop2=0.1/Mig=0
Bot=1000/Pop2=0.5/Mig=0.002
Bot=100/Pop2=0.1/Mig=0.002
Bot=1000/Pop2=1/Mig=0.002
Bot=100/Pop2=1/Mig=0
Bot=100/Pop2=0.01/Mig=0.002
Bot=100/Pop2=0.01/Mig=0
Bot=1000/Pop2=0.01/Mig=0.002
Bot=1000/Pop2=0.01/Mig=0
Bot=1000/Pop2=0.1/Mig=0
Bot=1000/Pop2=0.1/Mig=0.002
$F_{ST} + D_{XY}$
Bot=1000/Pop2=0.01/Mig=0
Bot=100/Pop2=1/Mig=0.002
Bot=1000/Pop2=0.01/Mig=0.002
Bot=1000/Pop2=0.1/Mig=0
Bot=100/Pop2=0.5/Mig=0.002
Bot=100/Pop2=0.1/Mig=0.002
Bot=1000/Pop2=0.5/Mig=0.002
Bot=100/Pop2=0.5/Mig=0
Bot=100/Pop2=0.1/Mig=0
Bot=1000/Pop2=1/Mig=0.002
Bot=1000/Pop2=0.5/Mig=0
Bot=100/Pop2=0.01/Mig=0.002
Bot=1000/Pop2=1/Mig=0
Bot=1000/Pop2=0.1/Mig=0.002
Bot=100/Pop2=0.01/Mig=0
$D_{XY}$
Bot=1000/Pop2=0.01/Mig=0.002
Bot=1000/Pop2=0.5/Mig=0
Bot=100/Pop2=1/Mig=0.002
Bot=1000/Pop2=0.5/Mig=0.002
Bot=1000/Pop2=1/Mig=0
Bot=100/Pop2=0.5/Mig=0.002
Bot=100/Pop2=1/Mig=0
Bot=1000/Pop2=1/Mig=0.002
Bot=100/Pop2=0.5/Mig=0
Bot=1000/Pop2=0.1/Mig=0.002
Bot=1000/Pop2=0.1/Mig=0
Bot=100/Pop2=0.1/Mig=0.002
Bot=100/Pop2=0.1/Mig=0
Bot=100/Pop2=0.01/Mig=0
Bot=1000/Pop2=0.01/Mig=0
$F_{ST} + \Delta\pi$
Bot=100/Pop2=1/Mig=0.002
Bot=1000/Pop2=0.5/Mig=0.002
Bot=100/Pop2=0.1/Mig=0.002
Bot=1000/Pop2=1/Mig=0
Bot=100/Pop2=0.5/Mig=0.002
Bot=1000/Pop2=0.5/Mig=0
Bot=100/Pop2=0.1/Mig=0
Bot=100/Pop2=0.5/Mig=0
Bot=1000/Pop2=0.1/Mig=0.002
Bot=1000/Pop2=1/Mig=0.002
Bot=100/Pop2=0.01/Mig=0.002
Bot=1000/Pop2=0.1/Mig=0
Bot=1000/Pop2=0.01/Mig=0.002
Bot=100/Pop2=0.01/Mig=0
Bot=1000/Pop2=0.01/Mig=0
$\Delta\pi$
Bot=100/Pop2=0.5/Mig=0
Bot=100/Pop2=1/Mig=0
Bot=1000/Pop2=0.5/Mig=0
Bot=100/Pop2=0.1/Mig=0.002
Bot=100/Pop2=1/Mig=0.002
Bot=1000/Pop2=0.1/Mig=0
Bot=1000/Pop2=1/Mig=0.002
Bot=1000/Pop2=0.5/Mig=0.002
Bot=100/Pop2=0.5/Mig=0.002
Bot=100/Pop2=0.01/Mig=0.002
Bot=1000/Pop2=0.01/Mig=0.002
Bot=1000/Pop2=0.01/Mig=0
Bot=100/Pop2=0.01/Mig=0
Bot=100/Pop2=0.1/Mig=0
Bot=1000/Pop2=0.1/Mig=0.002
$D_{XY} + \Delta\pi$
Bot=1000/Pop2=0.5/Mig=0.002
Bot=1000/Pop2=0.01/Mig=0.002
Bot=100/Pop2=1/Mig=0.002
Bot=100/Pop2=1/Mig=0
Bot=100/Pop2=0.01/Mig=0
Bot=1000/Pop2=1/Mig=0.002
Bot=1000/Pop2=0.01/Mig=0
Bot=1000/Pop2=0.5/Mig=0
Bot=1000/Pop2=1/Mig=0
Bot=100/Pop2=0.1/Mig=0
Bot=100/Pop2=0.5/Mig=0
Bot=1000/Pop2=0.1/Mig=0.002
Bot=100/Pop2=0.01/Mig=0.002
Bot=1000/Pop2=0.1/Mig=0
Bot=100/Pop2=0.1/Mig=0.002

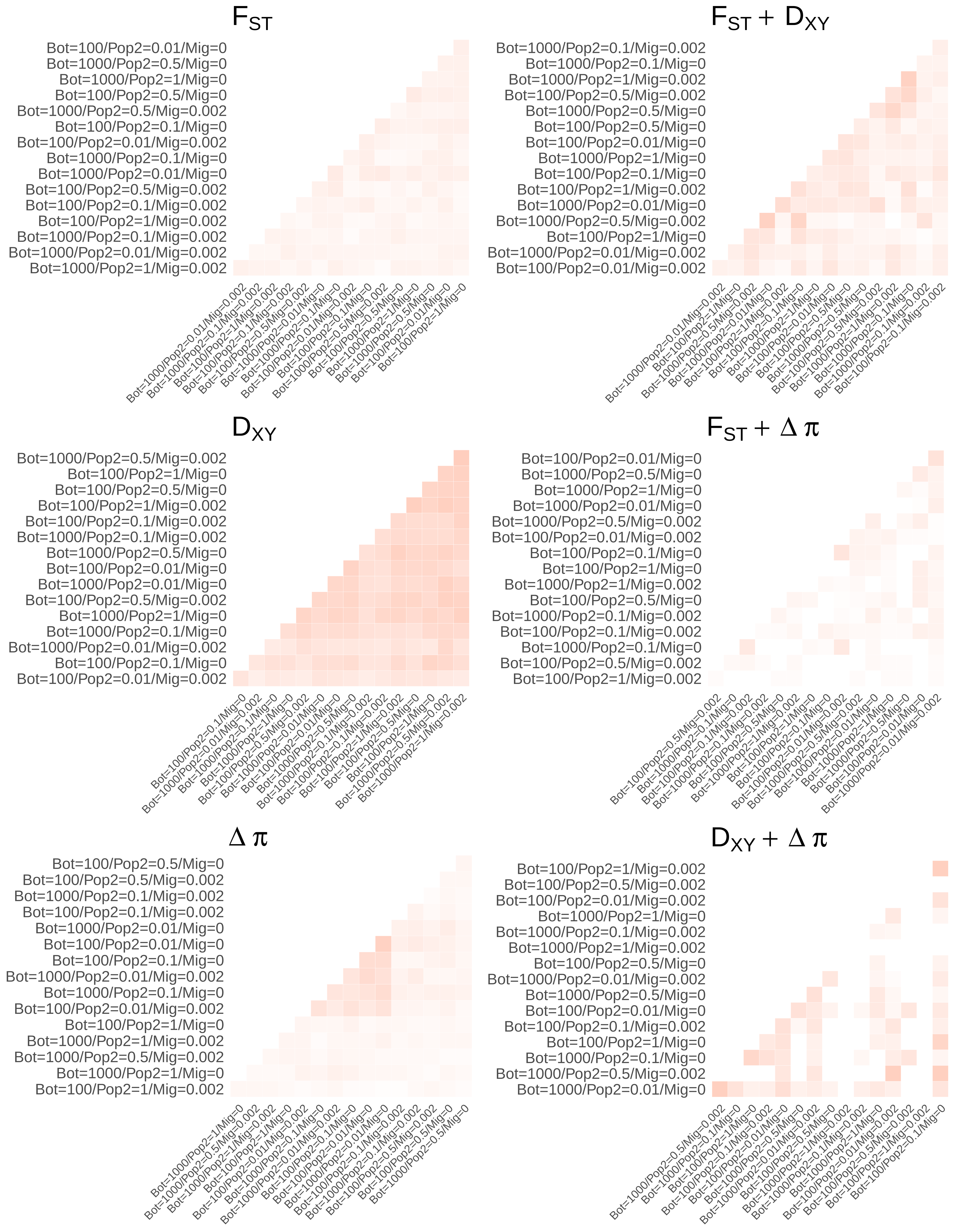

$F_{ST}$
Bot=100/Pop2=0.01/Mig=0
Bot=1000/Pop2=0.5/Mig=0
Bot=1000/Pop2=1/Mig=0
Bot=100/Pop2=0.5/Mig=0
Bot=1000/Pop2=0.5/Mig=0.002
Bot=100/Pop2=0.1/Mig=0
Bot=100/Pop2=0.01/Mig=0.002
Bot=1000/Pop2=0.1/Mig=0
Bot=1000/Pop2=0.01/Mig=0
Bot=100/Pop2=0.5/Mig=0.002
Bot=100/Pop2=0.1/Mig=0.002
Bot=100/Pop2=1/Mig=0.002
Bot=1000/Pop2=0.1/Mig=0.002
Bot=1000/Pop2=0.01/Mig=0.002
Bot=1000/Pop2=1/Mig=0.002
$F_{ST} + D_{XY}$
Bot=1000/Pop2=0.1/Mig=0.002
Bot=1000/Pop2=0.1/Mig=0
Bot=1000/Pop2=1/Mig=0.002
Bot=100/Pop2=0.5/Mig=0.002
Bot=1000/Pop2=0.5/Mig=0
Bot=100/Pop2=0.5/Mig=0
Bot=100/Pop2=0.01/Mig=0
Bot=1000/Pop2=1/Mig=0
Bot=100/Pop2=0.1/Mig=0
Bot=100/Pop2=1/Mig=0.002
Bot=1000/Pop2=0.01/Mig=0
Bot=1000/Pop2=0.5/Mig=0.002
Bot=100/Pop2=1/Mig=0
Bot=1000/Pop2=0.01/Mig=0.002
Bot=100/Pop2=0.01/Mig=0.002
$D_{XY}$
Bot=1000/Pop2=0.5/Mig=0.002
Bot=100/Pop2=1/Mig=0
Bot=100/Pop2=0.5/Mig=0
Bot=100/Pop2=1/Mig=0.002
Bot=100/Pop2=0.1/Mig=0.002
Bot=1000/Pop2=0.1/Mig=0.002
Bot=1000/Pop2=0.5/Mig=0
Bot=100/Pop2=0.01/Mig=0
Bot=1000/Pop2=0.01/Mig=0
Bot=100/Pop2=0.5/Mig=0.002
Bot=1000/Pop2=1/Mig=0
Bot=1000/Pop2=0.1/Mig=0
Bot=1000/Pop2=0.01/Mig=0.002
Bot=100/Pop2=0.1/Mig=0
Bot=100/Pop2=0.01/Mig=0.002
$F_{ST} + \Delta\pi$
Bot=100/Pop2=0.01/Mig=0
Bot=1000/Pop2=0.5/Mig=0
Bot=1000/Pop2=1/Mig=0
Bot=1000/Pop2=0.01/Mig=0
Bot=1000/Pop2=0.5/Mig=0.002
Bot=100/Pop2=0.01/Mig=0.002
Bot=100/Pop2=0.1/Mig=0
Bot=100/Pop2=1/Mig=0
Bot=1000/Pop2=1/Mig=0.002
Bot=100/Pop2=0.5/Mig=0
Bot=1000/Pop2=0.1/Mig=0.002
Bot=100/Pop2=0.1/Mig=0.002
Bot=1000/Pop2=0.1/Mig=0
Bot=100/Pop2=0.5/Mig=0.002
Bot=100/Pop2=1/Mig=0.002
$\Delta\pi$
Bot=100/Pop2=0.5/Mig=0
Bot=100/Pop2=0.5/Mig=0.002
Bot=1000/Pop2=0.1/Mig=0.002
Bot=100/Pop2=0.1/Mig=0.002
Bot=1000/Pop2=0.01/Mig=0
Bot=100/Pop2=0.01/Mig=0
Bot=100/Pop2=0.1/Mig=0
Bot=1000/Pop2=0.01/Mig=0.002
Bot=1000/Pop2=0.1/Mig=0
Bot=100/Pop2=0.01/Mig=0.002
Bot=100/Pop2=1/Mig=0
Bot=1000/Pop2=1/Mig=0.002
Bot=1000/Pop2=0.5/Mig=0.002
Bot=1000/Pop2=1/Mig=0
Bot=100/Pop2=1/Mig=0.002
$D_{XY} + \Delta\pi$
Bot=100/Pop2=1/Mig=0.002
Bot=100/Pop2=0.5/Mig=0.002
Bot=100/Pop2=0.01/Mig=0.002
Bot=1000/Pop2=1/Mig=0
Bot=1000/Pop2=0.1/Mig=0.002
Bot=1000/Pop2=1/Mig=0.002
Bot=100/Pop2=0.5/Mig=0
Bot=1000/Pop2=0.01/Mig=0.002
Bot=1000/Pop2=0.5/Mig=0
Bot=100/Pop2=0.01/Mig=0
Bot=100/Pop2=0.1/Mig=0.002
Bot=100/Pop2=1/Mig=0
Bot=1000/Pop2=0.1/Mig=0
Bot=1000/Pop2=0.5/Mig=0.002
Bot=1000/Pop2=0.01/Mig=0

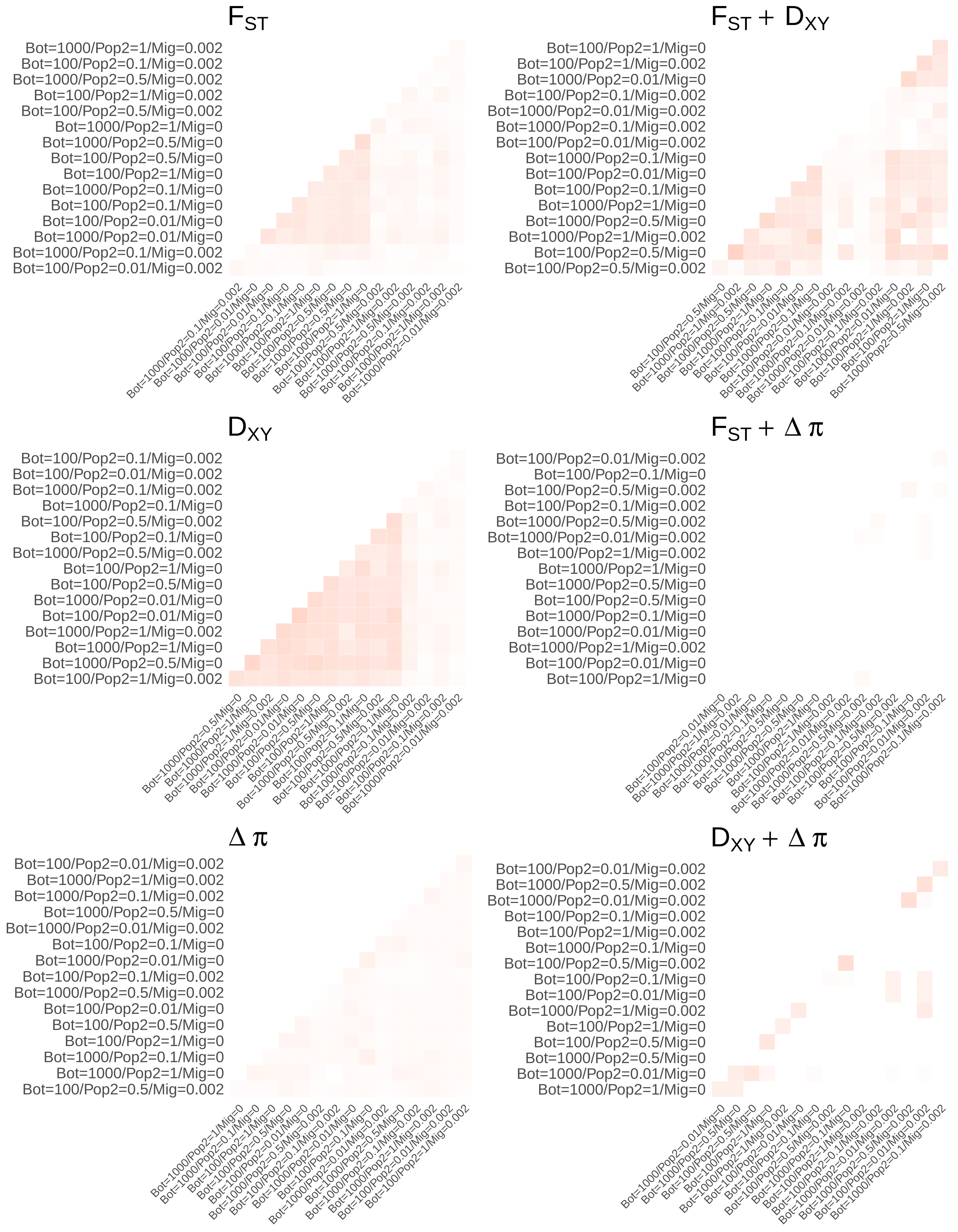

$F_{ST}$
Bot=1000/Pop2=1/Mig=0.002
Bot=100/Pop2=0.1/Mig=0.002
Bot=1000/Pop2=0.5/Mig=0.002
Bot=100/Pop2=1/Mig=0.002
Bot=100/Pop2=0.5/Mig=0.002
Bot=1000/Pop2=1/Mig=0
Bot=1000/Pop2=0.5/Mig=0
Bot=100/Pop2=0.5/Mig=0
Bot=100/Pop2=1/Mig=0
Bot=1000/Pop2=0.1/Mig=0
Bot=100/Pop2=0.1/Mig=0
Bot=100/Pop2=0.01/Mig=0
Bot=1000/Pop2=0.01/Mig=0
Bot=1000/Pop2=0.1/Mig=0.002
Bot=100/Pop2=0.01/Mig=0.002
$F_{ST} + D_{XY}$
Bot=100/Pop2=1/Mig=0
Bot=100/Pop2=1/Mig=0.002
Bot=1000/Pop2=0.01/Mig=0
Bot=100/Pop2=0.1/Mig=0.002
Bot=1000/Pop2=0.01/Mig=0.002
Bot=1000/Pop2=0.1/Mig=0.002
Bot=100/Pop2=0.01/Mig=0.002
Bot=1000/Pop2=0.1/Mig=0
Bot=100/Pop2=0.01/Mig=0
Bot=100/Pop2=0.1/Mig=0
Bot=1000/Pop2=1/Mig=0
Bot=1000/Pop2=0.5/Mig=0
Bot=1000/Pop2=1/Mig=0.002
Bot=100/Pop2=0.5/Mig=0
Bot=100/Pop2=0.5/Mig=0.002
$D_{XY}$
Bot=100/Pop2=0.1/Mig=0.002
Bot=100/Pop2=0.01/Mig=0.002
Bot=1000/Pop2=0.1/Mig=0.002
Bot=1000/Pop2=0.1/Mig=0
Bot=100/Pop2=0.5/Mig=0.002
Bot=100/Pop2=0.1/Mig=0
Bot=1000/Pop2=0.5/Mig=0.002
Bot=100/Pop2=1/Mig=0
Bot=100/Pop2=0.5/Mig=0
Bot=1000/Pop2=0.01/Mig=0
Bot=100/Pop2=0.01/Mig=0
Bot=1000/Pop2=1/Mig=0.002
Bot=1000/Pop2=1/Mig=0
Bot=1000/Pop2=0.5/Mig=0
Bot=100/Pop2=1/Mig=0.002
$F_{ST} + \Delta\pi$
Bot=100/Pop2=0.01/Mig=0.002
Bot=100/Pop2=0.1/Mig=0
Bot=100/Pop2=0.5/Mig=0.002
Bot=100/Pop2=0.1/Mig=0.002
Bot=1000/Pop2=0.5/Mig=0.002
Bot=1000/Pop2=0.01/Mig=0.002
Bot=100/Pop2=1/Mig=0.002
Bot=1000/Pop2=1/Mig=0
Bot=1000/Pop2=0.5/Mig=0
Bot=100/Pop2=0.5/Mig=0
Bot=1000/Pop2=0.1/Mig=0
Bot=1000/Pop2=0.01/Mig=0
Bot=1000/Pop2=1/Mig=0.002
Bot=100/Pop2=0.01/Mig=0
Bot=100/Pop2=1/Mig=0
$\Delta\pi$
Bot=100/Pop2=0.01/Mig=0.002
Bot=1000/Pop2=1/Mig=0.002
Bot=1000/Pop2=0.1/Mig=0.002
Bot=1000/Pop2=0.5/Mig=0
Bot=1000/Pop2=0.01/Mig=0.002
Bot=100/Pop2=0.1/Mig=0
Bot=1000/Pop2=0.01/Mig=0
Bot=100/Pop2=0.1/Mig=0.002
Bot=1000/Pop2=0.5/Mig=0.002
Bot=100/Pop2=0.01/Mig=0
Bot=100/Pop2=0.5/Mig=0
Bot=100/Pop2=1/Mig=0
Bot=1000/Pop2=0.1/Mig=0
Bot=1000/Pop2=1/Mig=0
Bot=100/Pop2=0.5/Mig=0.002
$D_{XY} + \Delta\pi$
Bot=100/Pop2=0.01/Mig=0.002
Bot=1000/Pop2=0.5/Mig=0.002
Bot=1000/Pop2=0.01/Mig=0.002
Bot=100/Pop2=0.1/Mig=0.002
Bot=100/Pop2=1/Mig=0.002
Bot=1000/Pop2=0.1/Mig=0
Bot=100/Pop2=0.5/Mig=0.002
Bot=100/Pop2=0.1/Mig=0
Bot=100/Pop2=0.01/Mig=0
Bot=1000/Pop2=1/Mig=0.002
Bot=100/Pop2=1/Mig=0
Bot=100/Pop2=0.5/Mig=0
Bot=1000/Pop2=0.5/Mig=0
Bot=1000/Pop2=0.01/Mig=0
Bot=1000/Pop2=1/Mig=0

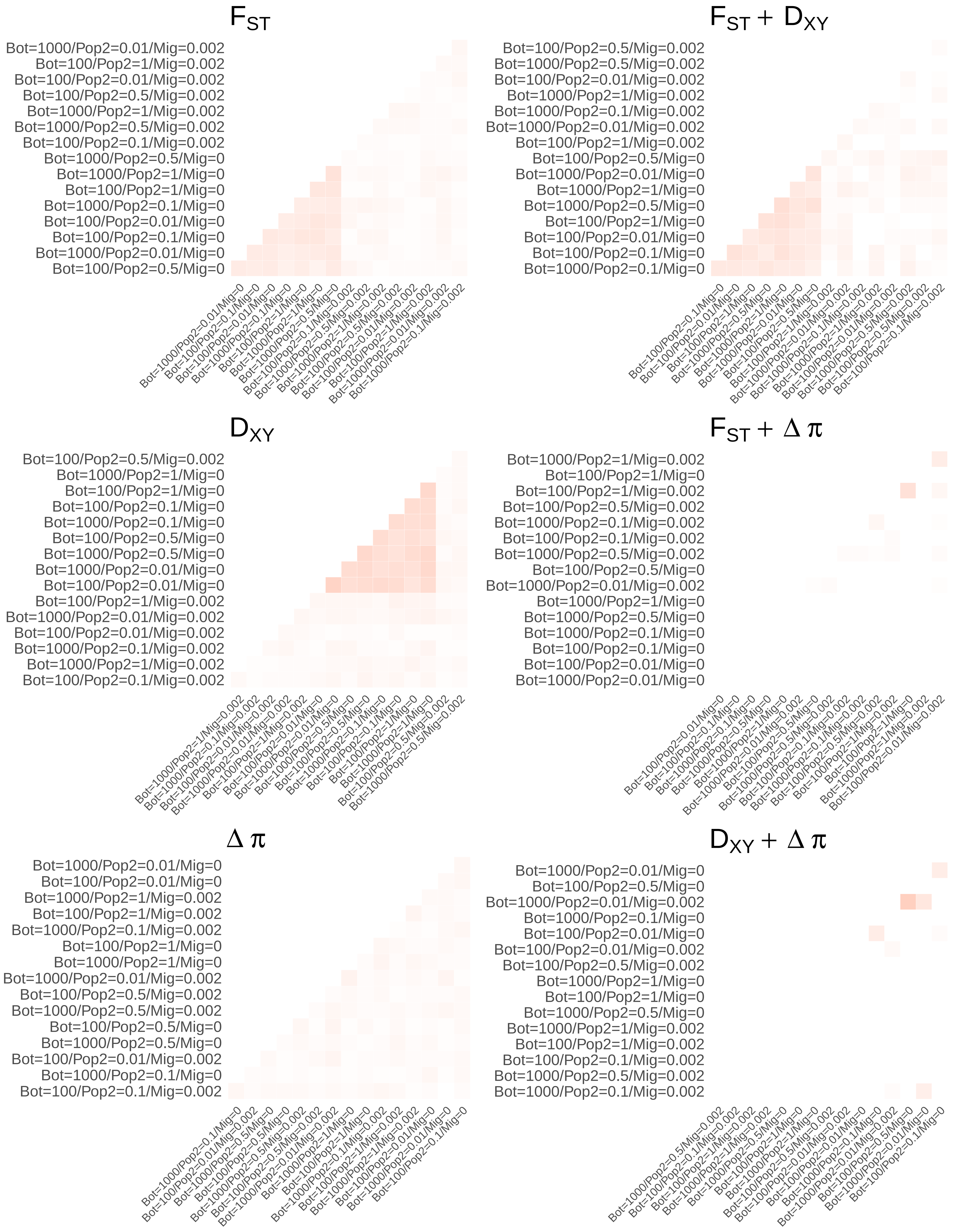
$F_{ST}$
Bot=1000/Pop2=0.01/Mig=0.002
Bot=100/Pop2=1/Mig=0.002
Bot=100/Pop2=0.01/Mig=0.002
Bot=100/Pop2=0.5/Mig=0.002
Bot=1000/Pop2=1/Mig=0.002
Bot=1000/Pop2=0.5/Mig=0.002
Bot=100/Pop2=0.1/Mig=0.002
Bot=1000/Pop2=0.5/Mig=0
Bot=1000/Pop2=1/Mig=0
Bot=100/Pop2=1/Mig=0
Bot=1000/Pop2=0.1/Mig=0
Bot=100/Pop2=0.01/Mig=0
Bot=100/Pop2=0.1/Mig=0
Bot=1000/Pop2=0.01/Mig=0
Bot=100/Pop2=0.5/Mig=0
$F_{ST}$ + $D_{XY}$
Bot=100/Pop2=0.5/Mig=0.002
Bot=1000/Pop2=0.5/Mig=0.002
Bot=100/Pop2=0.01/Mig=0.002
Bot=1000/Pop2=1/Mig=0.002
Bot=1000/Pop2=0.1/Mig=0.002
Bot=1000/Pop2=0.01/Mig=0.002
Bot=100/Pop2=1/Mig=0.002
Bot=100/Pop2=0.5/Mig=0
Bot=1000/Pop2=0.01/Mig=0
Bot=1000/Pop2=1/Mig=0
Bot=1000/Pop2=0.5/Mig=0
Bot=100/Pop2=1/Mig=0
Bot=100/Pop2=0.01/Mig=0
Bot=100/Pop2=0.1/Mig=0
Bot=1000/Pop2=0.1/Mig=0
$D_{XY}$
Bot=100/Pop2=0.5/Mig=0.002
Bot=1000/Pop2=1/Mig=0
Bot=100/Pop2=1/Mig=0
Bot=100/Pop2=0.1/Mig=0
Bot=1000/Pop2=0.1/Mig=0
Bot=100/Pop2=0.5/Mig=0
Bot=1000/Pop2=0.5/Mig=0
Bot=1000/Pop2=0.01/Mig=0
Bot=100/Pop2=0.01/Mig=0
Bot=100/Pop2=1/Mig=0.002
Bot=1000/Pop2=0.01/Mig=0.002
Bot=100/Pop2=0.01/Mig=0.002
Bot=1000/Pop2=0.1/Mig=0.002
Bot=1000/Pop2=1/Mig=0.002
Bot=100/Pop2=0.1/Mig=0.002
$F_{ST}$ + $\Delta\pi$
Bot=1000/Pop2=1/Mig=0.002
Bot=100/Pop2=1/Mig=0
Bot=100/Pop2=1/Mig=0.002
Bot=100/Pop2=0.5/Mig=0.002
Bot=1000/Pop2=0.1/Mig=0.002
Bot=100/Pop2=0.1/Mig=0.002
Bot=1000/Pop2=0.5/Mig=0.002
Bot=100/Pop2=0.5/Mig=0
Bot=1000/Pop2=0.01/Mig=0.002
Bot=1000/Pop2=1/Mig=0
Bot=1000/Pop2=0.5/Mig=0
Bot=1000/Pop2=0.1/Mig=0
Bot=100/Pop2=0.1/Mig=0
Bot=100/Pop2=0.01/Mig=0
Bot=1000/Pop2=0.01/Mig=0
$\Delta\pi$
Bot=1000/Pop2=0.01/Mig=0
Bot=100/Pop2=0.01/Mig=0
Bot=1000/Pop2=1/Mig=0.002
Bot=100/Pop2=1/Mig=0.002
Bot=1000/Pop2=0.1/Mig=0.002
Bot=100/Pop2=1/Mig=0
Bot=1000/Pop2=1/Mig=0
Bot=1000/Pop2=0.01/Mig=0.002
Bot=100/Pop2=0.5/Mig=0.002
Bot=1000/Pop2=0.5/Mig=0.002
Bot=100/Pop2=0.5/Mig=0
Bot=1000/Pop2=0.5/Mig=0
Bot=100/Pop2=0.01/Mig=0.002
Bot=1000/Pop2=0.1/Mig=0
Bot=100/Pop2=0.1/Mig=0.002
$D_{XY}$ + $\Delta\pi$
Bot=1000/Pop2=0.01/Mig=0
Bot=100/Pop2=0.5/Mig=0
Bot=1000/Pop2=0.01/Mig=0.002
Bot=1000/Pop2=0.1/Mig=0
Bot=100/Pop2=0.01/Mig=0
Bot=100/Pop2=0.01/Mig=0.002
Bot=100/Pop2=0.5/Mig=0.002
Bot=1000/Pop2=1/Mig=0
Bot=100/Pop2=1/Mig=0
Bot=1000/Pop2=0.5/Mig=0
Bot=1000/Pop2=1/Mig=0.002
Bot=100/Pop2=1/Mig=0.002
Bot=100/Pop2=0.1/Mig=0.002
Bot=1000/Pop2=0.5/Mig=0.002
Bot=1000/Pop2=0.1/Mig=0.002

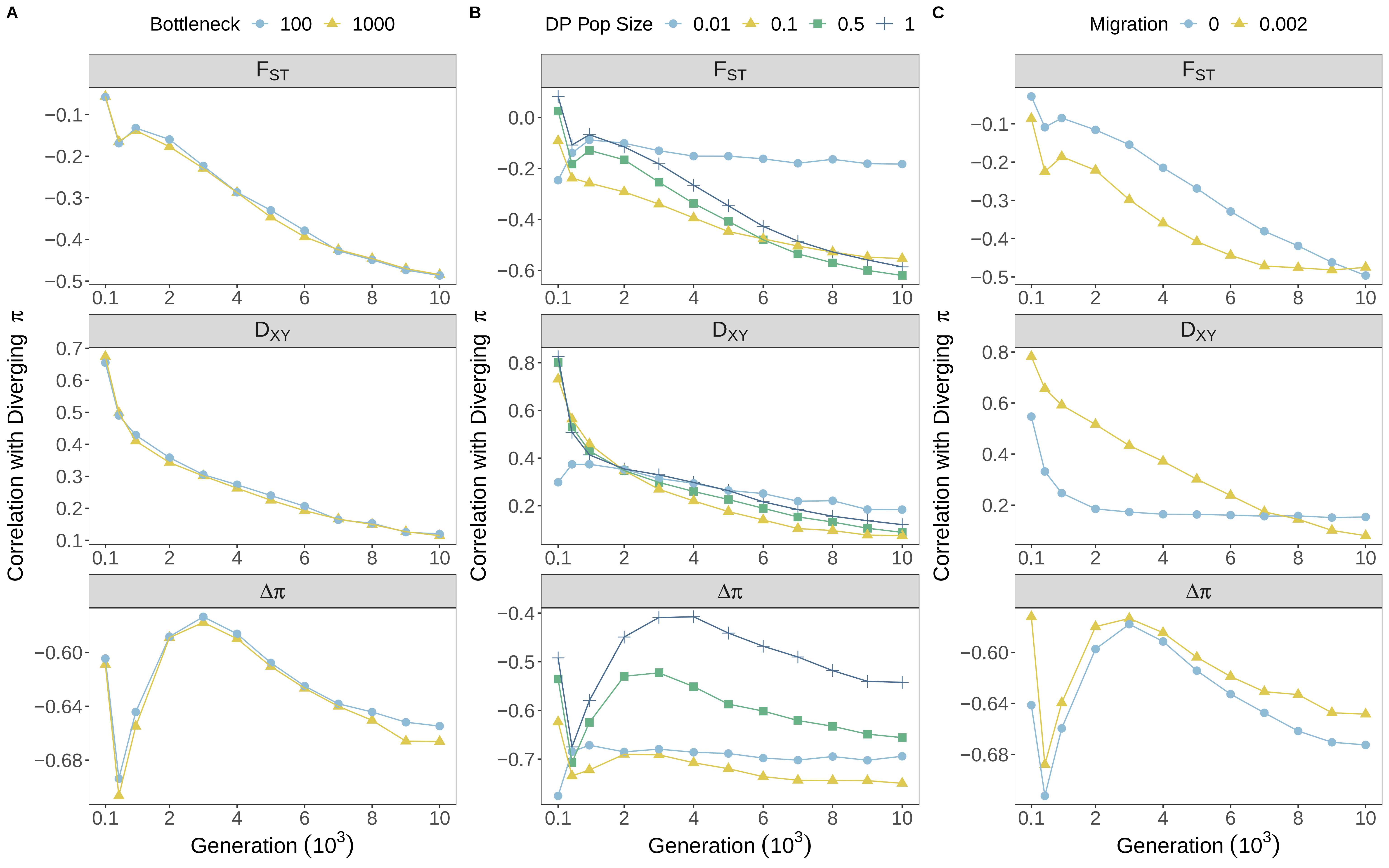

A
Bottleneck 100 1000
B
DP Pop Size 0.01 0.1 0.5 1
C
Migration 0 0.002
F_ST
D_XY
Δπ
Correlation with Diverging π
Generation (10^3)
0.1
2
4
6
8
10

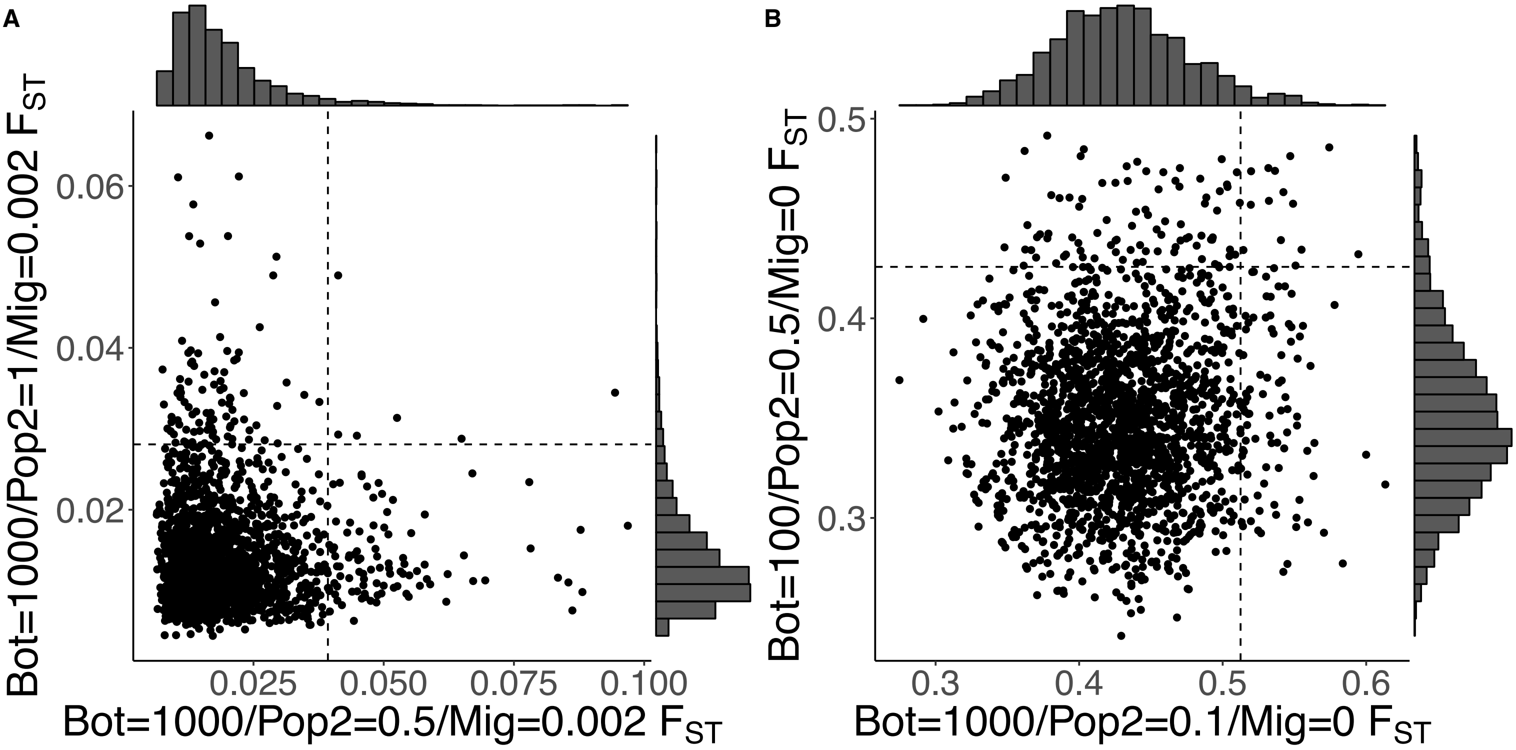

A
B
0.06
0.04
0.02
0.025
0.050
0.075
0.100
Bot=1000/Pop2=1/Mig=0.002 $F_{ST}$
Bot=1000/Pop2=0.5/Mig=0.002 $F_{ST}$
0.5
0.4
0.3
0.3
0.4
0.5
0.6
Bot=100/Pop2=0.5/Mig=0 $F_{ST}$
Bot=1000/Pop2=0.1/Mig=0 $F_{ST}$

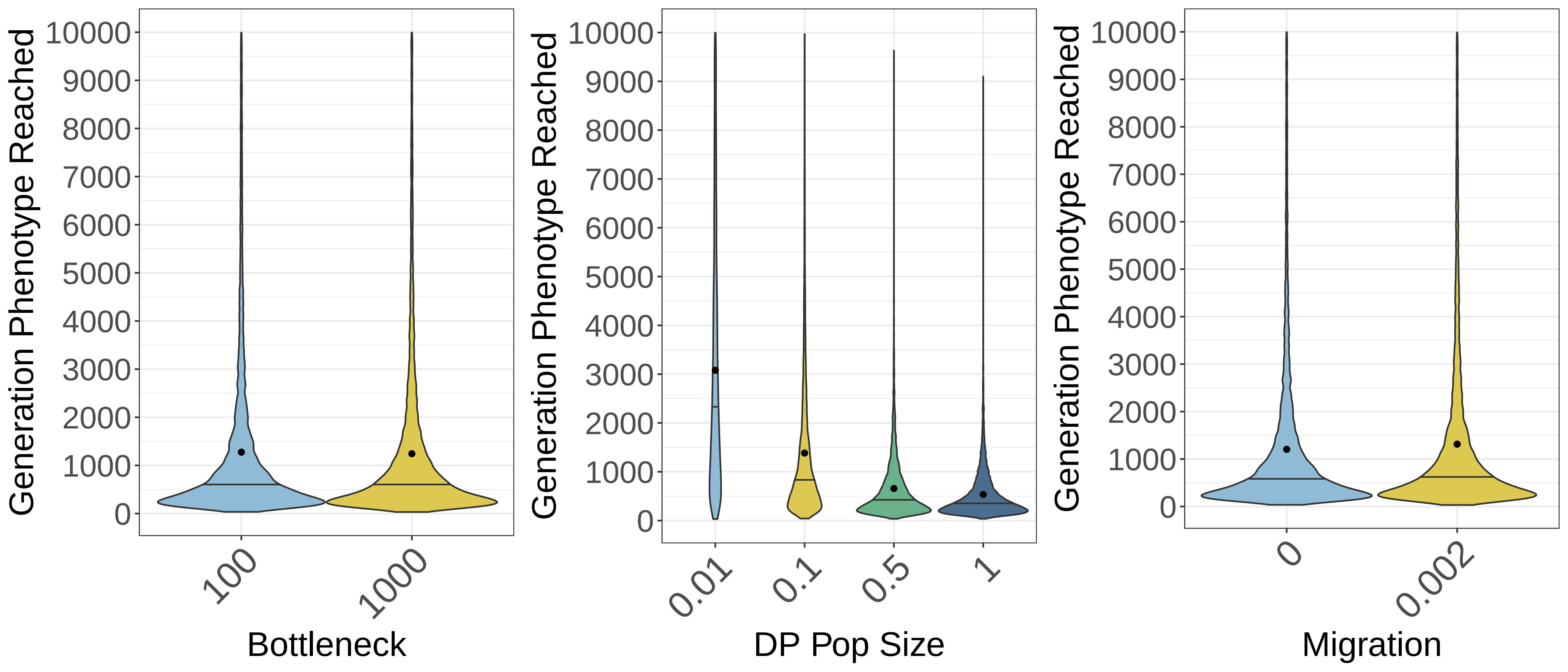

Generation Phenotype Reached
10000
9000
8000
7000
6000
5000
4000
3000
2000
1000
0
100
1000
Bottleneck
0.01
0.1
0.5
1
DP Pop Size
0
0.002
Migration

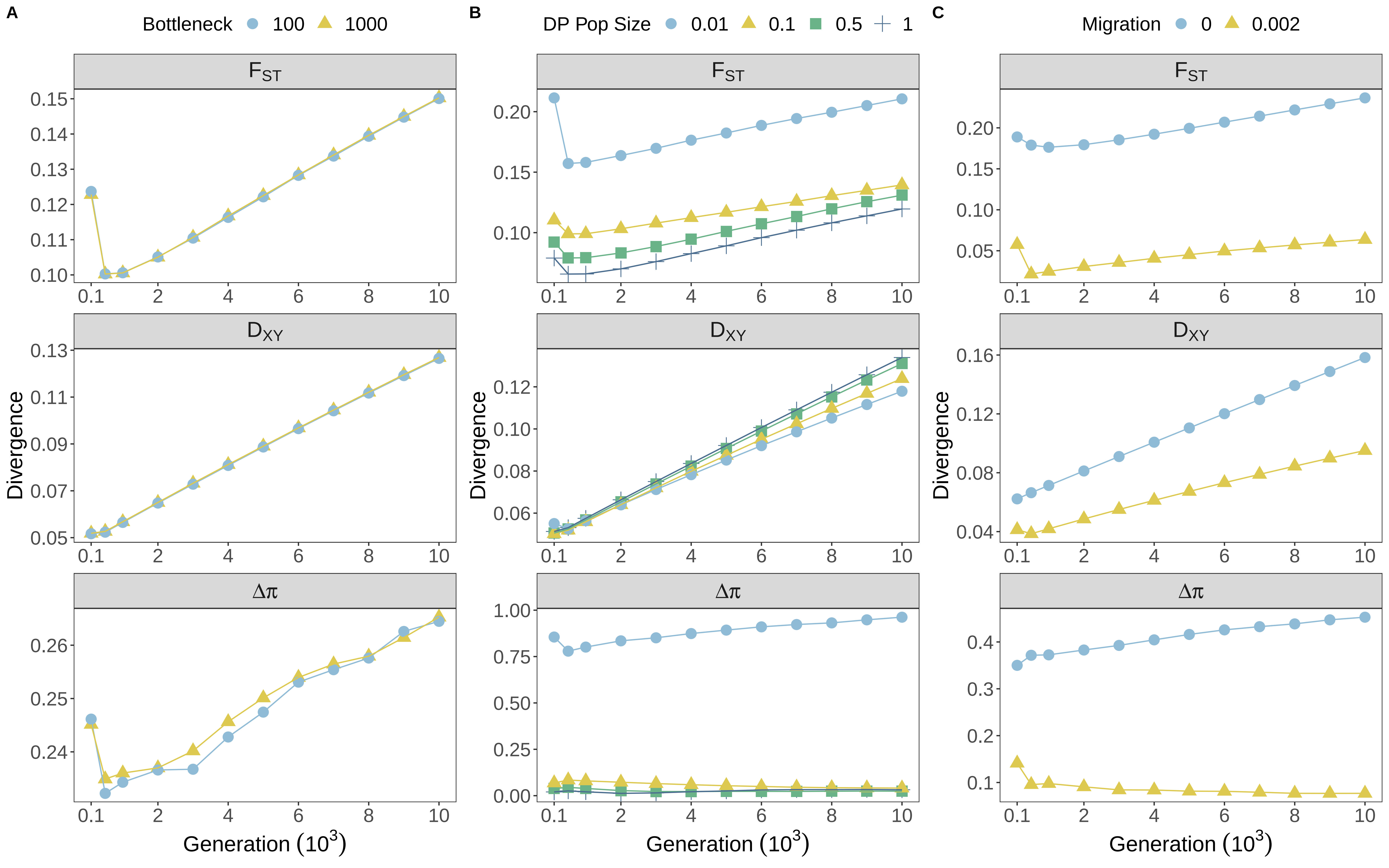

A
Bottleneck 100 1000
B
DP Pop Size 0.01 0.1 0.5 1
C
Migration 0 0.002
F_ST
D_XY
Δπ
Divergence
Generation (10^3)
0.1 2 4 6 8 10

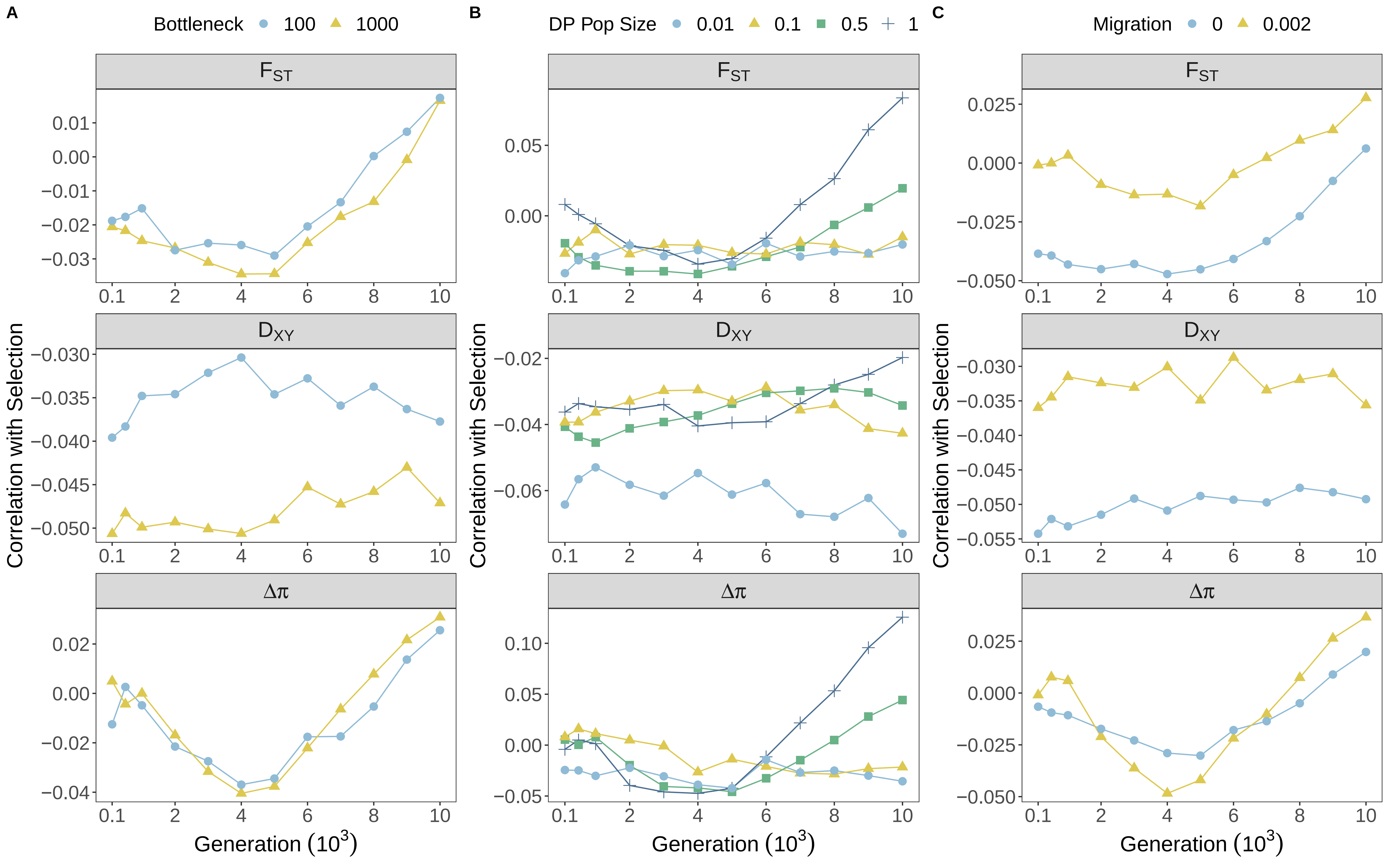
A
Bottleneck 100 1000
B
DP Pop Size 0.01 0.1 0.5 1
C
Migration 0 0.002
$F_{ST}$
$D_{XY}$
$\Delta\pi$
Correlation with Selection
Generation ($10^3$)

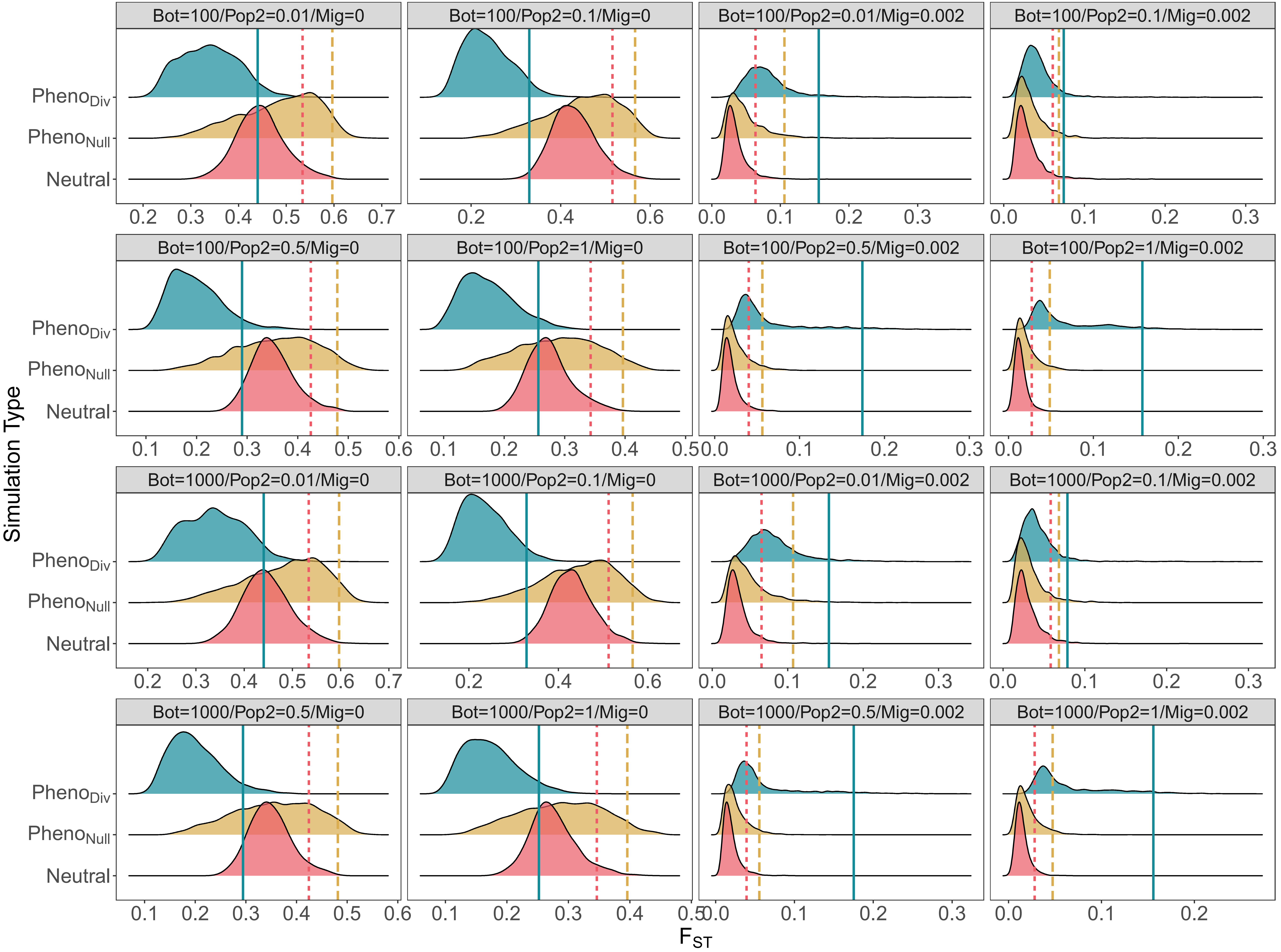

Bot=100/Pop2=0.01/Mig=0
Bot=100/Pop2=0.1/Mig=0
Bot=100/Pop2=0.01/Mig=0.002
Bot=100/Pop2=0.1/Mig=0.002
Bot=100/Pop2=0.5/Mig=0
Bot=100/Pop2=1/Mig=0
Bot=100/Pop2=0.5/Mig=0.002
Bot=100/Pop2=1/Mig=0.002
Bot=1000/Pop2=0.01/Mig=0
Bot=1000/Pop2=0.1/Mig=0
Bot=1000/Pop2=0.01/Mig=0.002
Bot=1000/Pop2=0.1/Mig=0.002
Bot=1000/Pop2=0.5/Mig=0
Bot=1000/Pop2=1/Mig=0
Bot=1000/Pop2=0.5/Mig=0.002
Bot=1000/Pop2=1/Mig=0.002
Simulation Type
$Pheno_{Div}$
$Pheno_{Null}$
Neutral
$F_{ST}$

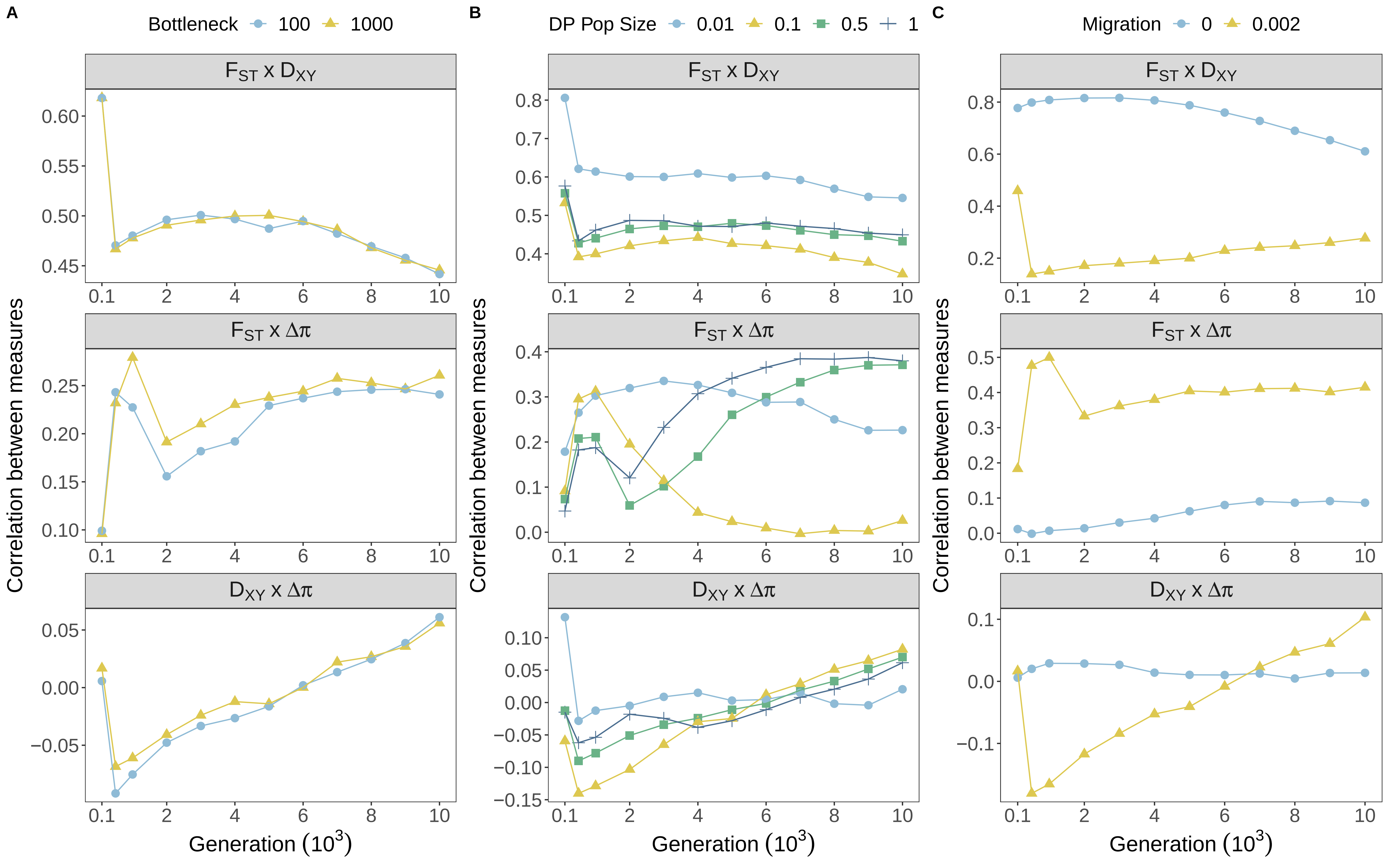

A
Bottleneck
100
1000
B
DP Pop Size
0.01
0.1
0.5
1
C
Migration
0
0.002
F_ST x D_XY
F_ST x Δπ
D_XY x Δπ
Correlation between measures
Generation (10^3)

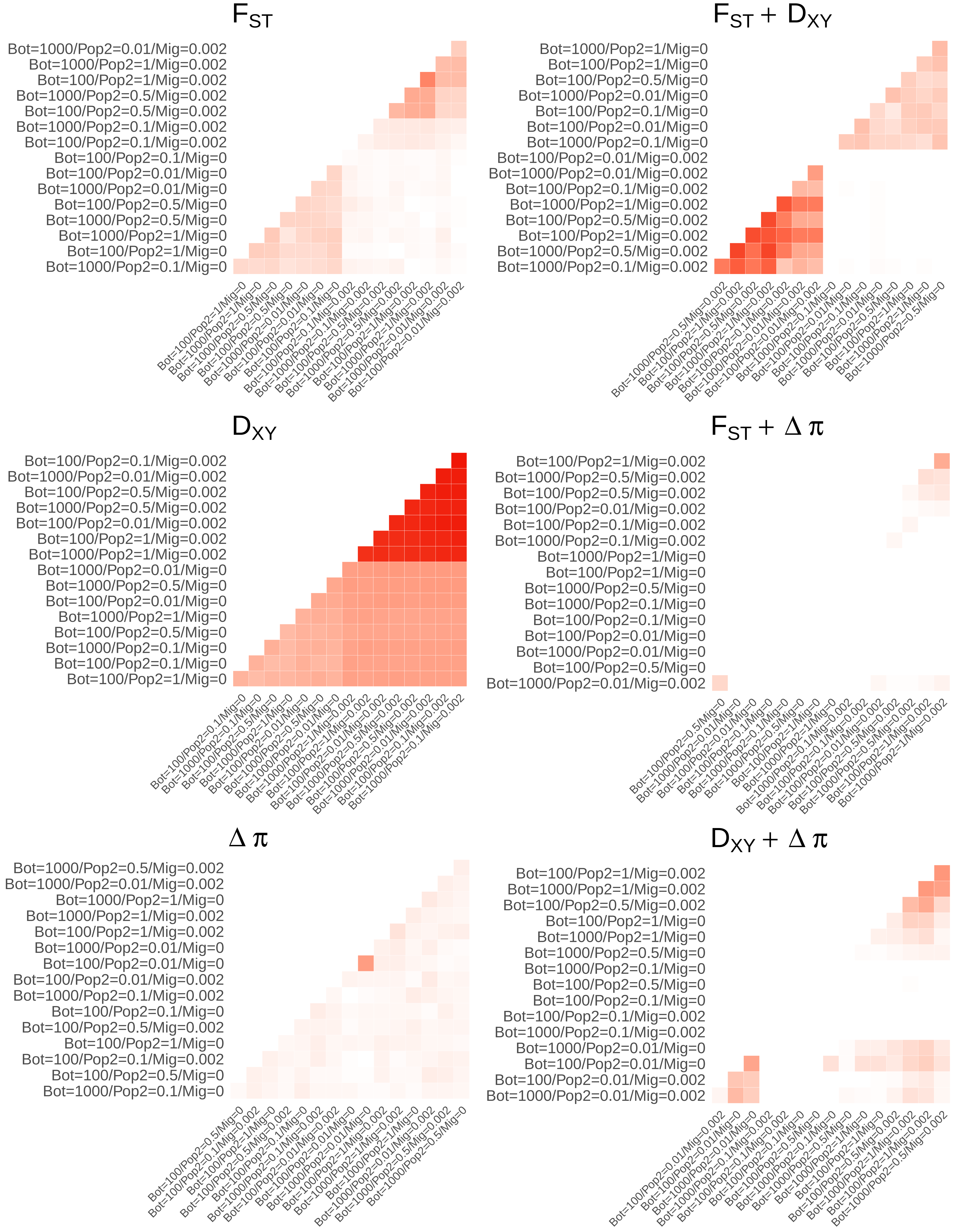
$F_{ST}$
Bot=1000/Pop2=0.01/Mig=0.002
Bot=1000/Pop2=1/Mig=0.002
Bot=100/Pop2=1/Mig=0.002
Bot=1000/Pop2=0.5/Mig=0.002
Bot=100/Pop2=0.5/Mig=0.002
Bot=1000/Pop2=0.1/Mig=0.002
Bot=100/Pop2=0.1/Mig=0.002
Bot=100/Pop2=0.1/Mig=0
Bot=100/Pop2=0.01/Mig=0
Bot=1000/Pop2=0.01/Mig=0
Bot=100/Pop2=0.5/Mig=0
Bot=1000/Pop2=0.5/Mig=0
Bot=1000/Pop2=1/Mig=0
Bot=100/Pop2=1/Mig=0
Bot=1000/Pop2=0.1/Mig=0
$F_{ST} + D_{XY}$
Bot=1000/Pop2=1/Mig=0
Bot=100/Pop2=1/Mig=0
Bot=100/Pop2=0.5/Mig=0
Bot=1000/Pop2=0.01/Mig=0
Bot=100/Pop2=0.1/Mig=0
Bot=100/Pop2=0.01/Mig=0
Bot=1000/Pop2=0.1/Mig=0
Bot=100/Pop2=0.01/Mig=0.002
Bot=1000/Pop2=0.01/Mig=0.002
Bot=100/Pop2=0.1/Mig=0.002
Bot=1000/Pop2=1/Mig=0.002
Bot=100/Pop2=0.5/Mig=0.002
Bot=100/Pop2=1/Mig=0.002
Bot=1000/Pop2=0.5/Mig=0.002
Bot=1000/Pop2=0.1/Mig=0.002
$D_{XY}$
Bot=100/Pop2=0.1/Mig=0.002
Bot=1000/Pop2=0.01/Mig=0.002
Bot=100/Pop2=0.5/Mig=0.002
Bot=1000/Pop2=0.5/Mig=0.002
Bot=100/Pop2=0.01/Mig=0.002
Bot=100/Pop2=1/Mig=0.002
Bot=1000/Pop2=1/Mig=0.002
Bot=1000/Pop2=0.01/Mig=0
Bot=1000/Pop2=0.5/Mig=0
Bot=100/Pop2=0.01/Mig=0
Bot=1000/Pop2=1/Mig=0
Bot=100/Pop2=0.5/Mig=0
Bot=1000/Pop2=0.1/Mig=0
Bot=100/Pop2=0.1/Mig=0
Bot=100/Pop2=1/Mig=0
$F_{ST} + \Delta\pi$
Bot=100/Pop2=1/Mig=0.002
Bot=1000/Pop2=0.5/Mig=0.002
Bot=100/Pop2=0.5/Mig=0.002
Bot=100/Pop2=0.01/Mig=0.002
Bot=100/Pop2=0.1/Mig=0.002
Bot=1000/Pop2=0.1/Mig=0.002
Bot=1000/Pop2=1/Mig=0
Bot=100/Pop2=1/Mig=0
Bot=1000/Pop2=0.5/Mig=0
Bot=1000/Pop2=0.1/Mig=0
Bot=100/Pop2=0.1/Mig=0
Bot=100/Pop2=0.01/Mig=0
Bot=1000/Pop2=0.01/Mig=0
Bot=100/Pop2=0.5/Mig=0
Bot=1000/Pop2=0.01/Mig=0.002
$\Delta\pi$
Bot=1000/Pop2=0.5/Mig=0.002
Bot=1000/Pop2=0.01/Mig=0.002
Bot=1000/Pop2=1/Mig=0
Bot=1000/Pop2=1/Mig=0.002
Bot=100/Pop2=1/Mig=0.002
Bot=1000/Pop2=0.01/Mig=0
Bot=100/Pop2=0.01/Mig=0
Bot=100/Pop2=0.01/Mig=0.002
Bot=1000/Pop2=0.1/Mig=0.002
Bot=100/Pop2=0.1/Mig=0
Bot=100/Pop2=0.5/Mig=0.002
Bot=100/Pop2=1/Mig=0
Bot=100/Pop2=0.1/Mig=0.002
Bot=100/Pop2=0.5/Mig=0
Bot=1000/Pop2=0.1/Mig=0
$D_{XY} + \Delta\pi$
Bot=100/Pop2=1/Mig=0.002
Bot=1000/Pop2=1/Mig=0.002
Bot=100/Pop2=0.5/Mig=0.002
Bot=100/Pop2=1/Mig=0
Bot=1000/Pop2=1/Mig=0
Bot=1000/Pop2=0.5/Mig=0
Bot=1000/Pop2=0.1/Mig=0
Bot=100/Pop2=0.5/Mig=0
Bot=100/Pop2=0.1/Mig=0
Bot=100/Pop2=0.1/Mig=0.002
Bot=1000/Pop2=0.1/Mig=0.002
Bot=1000/Pop2=0.01/Mig=0
Bot=100/Pop2=0.01/Mig=0
Bot=100/Pop2=0.01/Mig=0.002
Bot=1000/Pop2=0.01/Mig=0.002

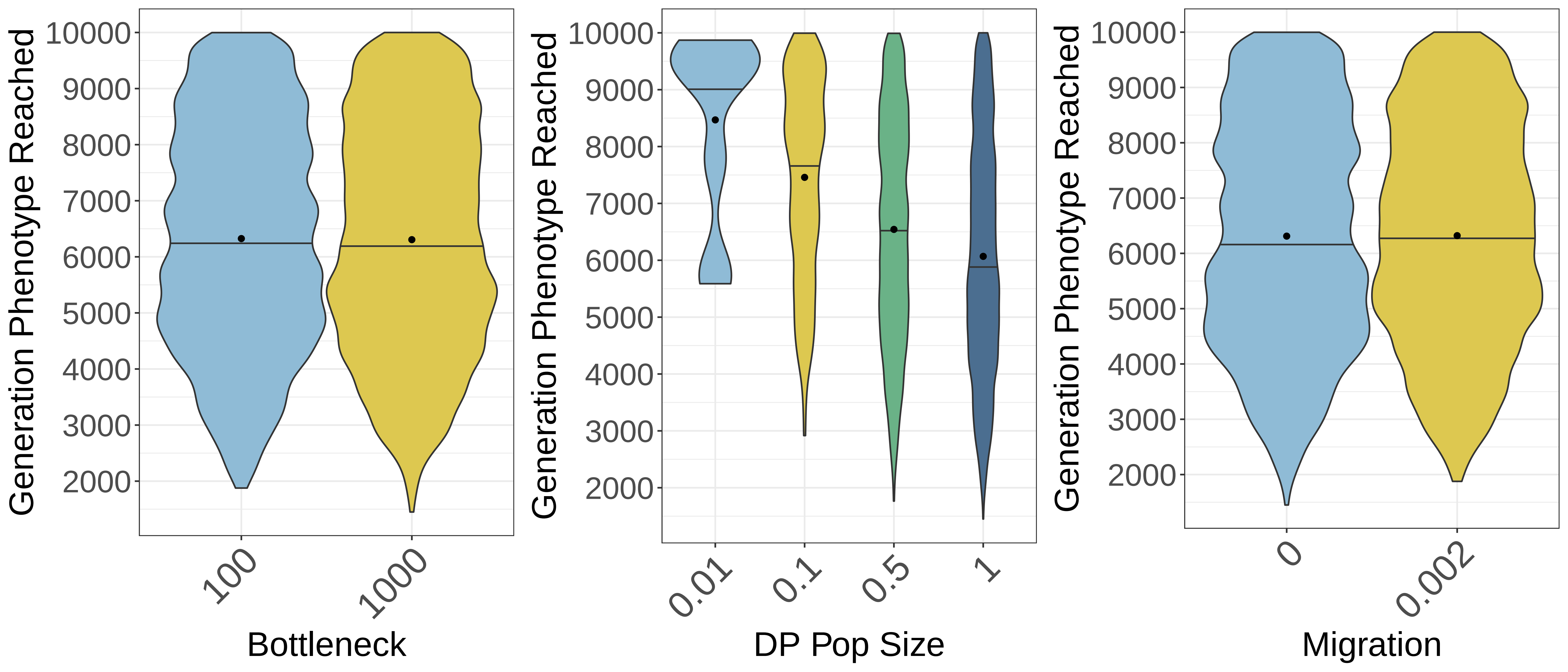

Generation Phenotype Reached
10000
9000
8000
7000
6000
5000
4000
3000
2000
100
1000
Bottleneck
0.01
0.1
0.5
1
DP Pop Size
0
0.002
Migration

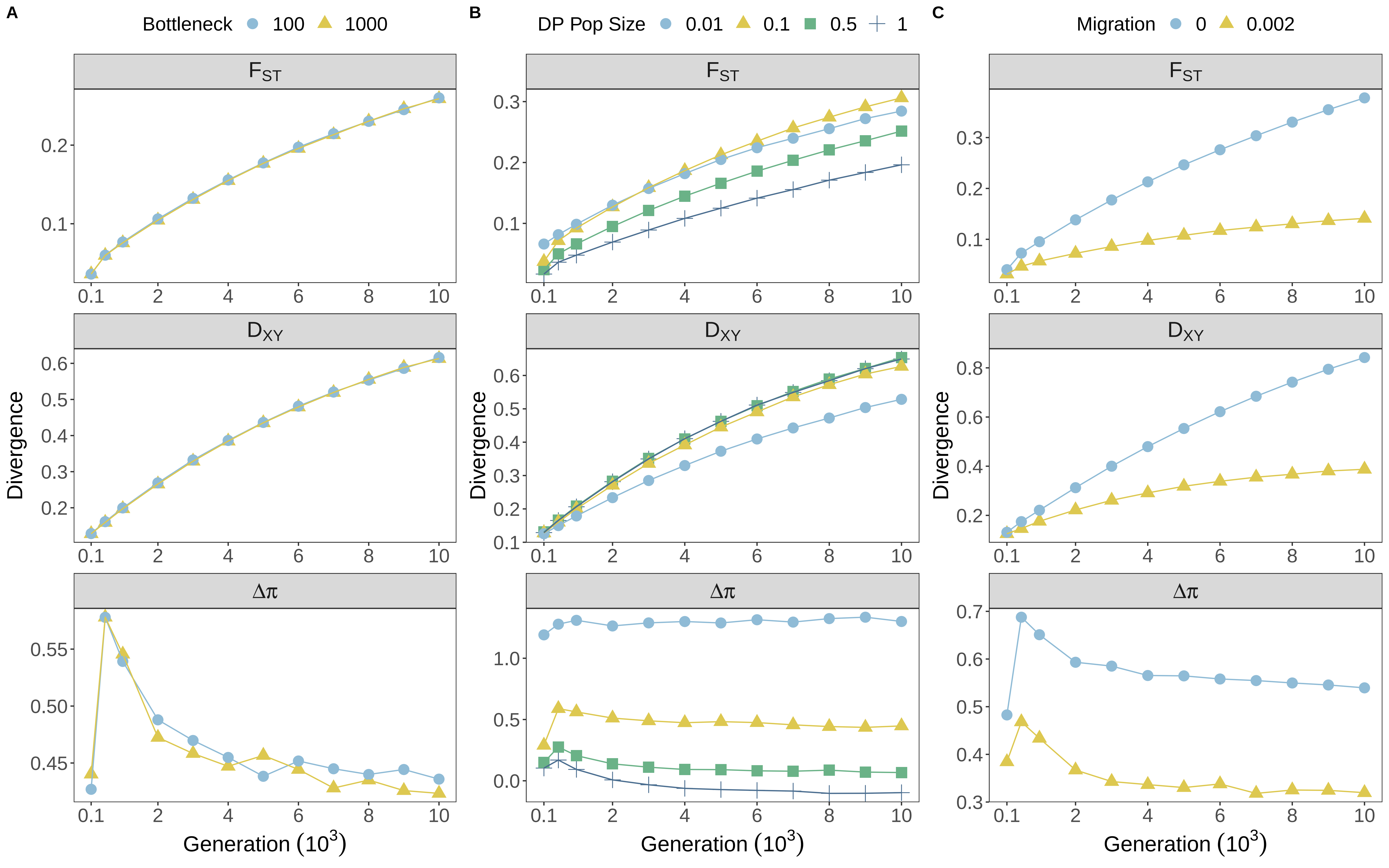

A
Bottleneck 100 1000
B
DP Pop Size 0.01 0.1 0.5 1
C
Migration 0 0.002
$F_{ST}$
$D_{XY}$
$\Delta\pi$
Divergence
Generation ($10^3$)

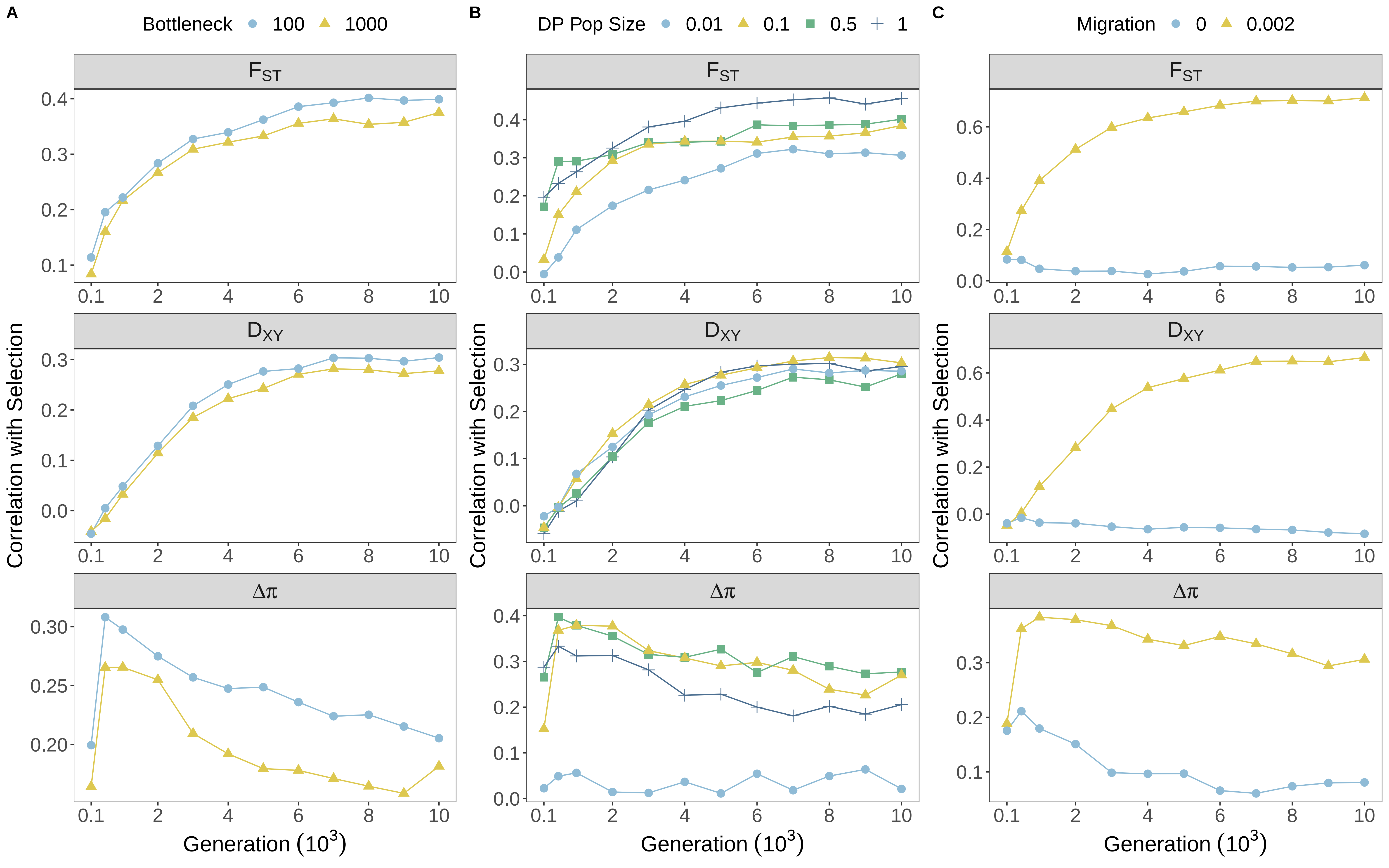

A
Bottleneck 100 1000
B
DP Pop Size 0.01 0.1 0.5 1
C
Migration 0 0.002
$F_{ST}$
$D_{XY}$
$\Delta\pi$
Correlation with Selection
Generation ($10^3$)

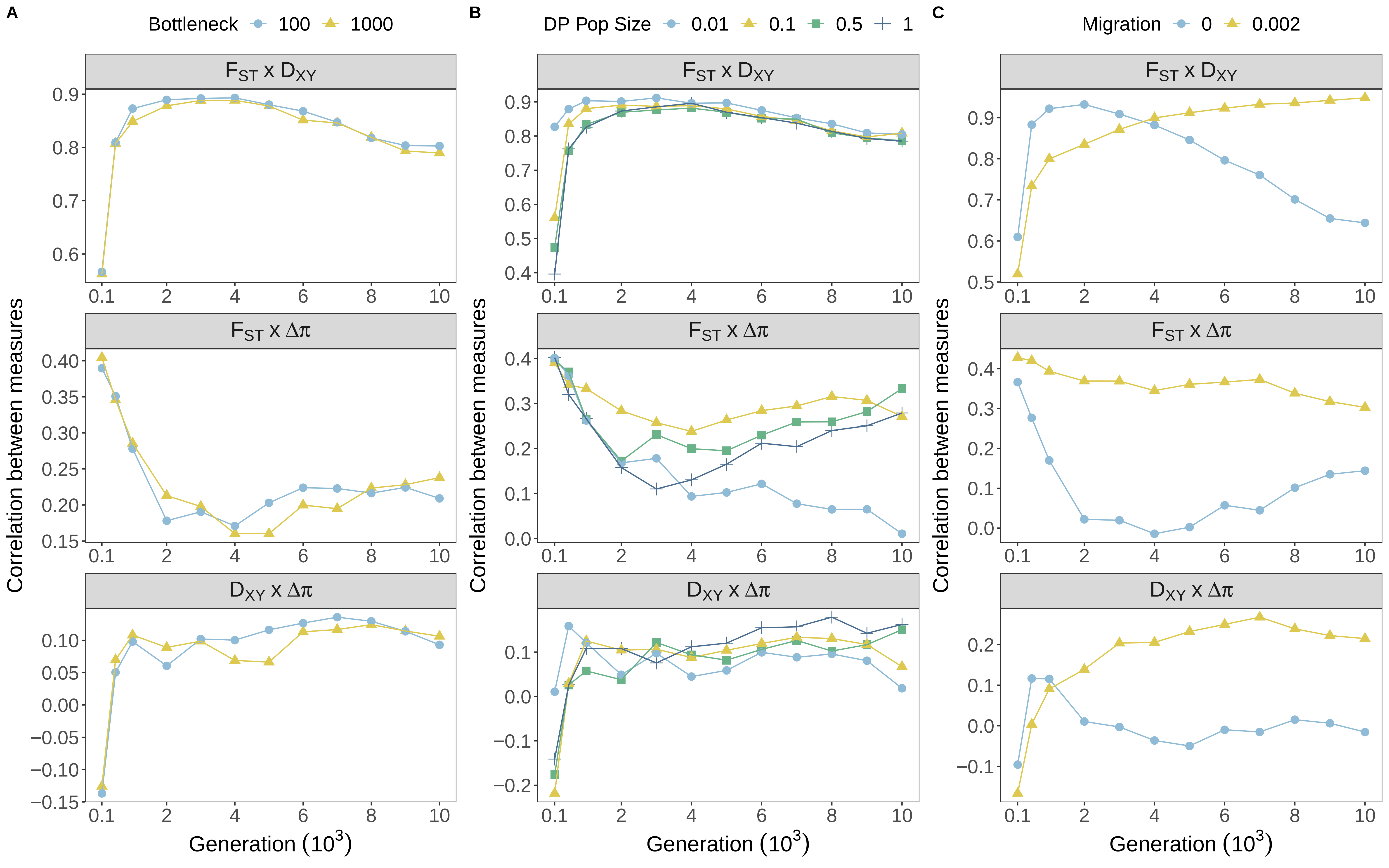

A
Bottleneck 100 1000
B
DP Pop Size 0.01 0.1 0.5 1
C
Migration 0 0.002
$F_{ST}$ x $D_{XY}$
$F_{ST}$ x Δπ
$D_{XY}$ x Δπ
Correlation between measures
Generation ($10^3$)
