## Supplementary material for "Contingent Convergence: The ability to detect convergent genomic evolution is dependent on population size and migration": Supp Figure Legends

**Figure S30:** The proportional overlap of outliers above the 95% quantile, averaged across 100 downsampled datasets consisting of 95% Neutral and 5% $Pheno_{Null}$ data for each of the 16 demographic treatments after 500 generations. Axis orderings were determined through hierarchical clustering. Heatmaps are shown for single measures of $F_{ST}$, $D_{XY}$ and $\Delta\pi$ in the first column, and combined measures in the second column. Heatmaps are coloured according to a common scale of 0 to 1. Treatments are labelled with founding bottleneck (Bot), DP population size (Pop2), and migration (Mig) values.

**Figure S31:** The proportional overlap of outliers above the 95% quantile, averaged across 100 downsampled datasets consisting of 95% Neutral and 5% $Pheno_{Null}$ data for each of the 16 demographic treatments after 3000 generations. Axis orderings were determined through hierarchical clustering. Heatmaps are shown for single measures of $F_{ST}$, $D_{XY}$ and $\Delta\pi$ in the first column, and combined measures in the second column. Heatmaps are coloured according to a common scale of 0 to 1. Treatments are labelled with founding bottleneck (Bot), DP population size (Pop2), and migration (Mig) values.

**Figure S37:** Effects of demographic treatments on relationships between measures of divergence and $DP_{\pi}$ across all sampling generations for $\mu = 4.89e^{-6}$ simulations. Point colour and shape denote treatment groups for founding bottlenecks (**A**), protracted bottlenecks (**B**) and migration (**C**). Each point represents correlation coefficients calculated across all genes within individual treatments groups, averaged within treatment levels, and averaged over 20 iterations.

**Figure S38:** The effect of migration on distributions of $F_{ST}$ and their random overlap. Data are plotted for $F_{ST}$ calculated over the 20 iterations of 100 genomic windows under neutrality. Shown are the overlap between two treatments with (**A**) and without (**B**) migration. In each figure, 95% quantiles are plotted for each distribution as dashed lines. Distributions for x and y data are plotted in axis margins.

**Figure S39:** The effect of demographic treatments on the generation DP reached its phenotypic optimum (trait value > 9). Violins denote distributions calculated over all iterations of genome windows, organised according to founding bottlenecks, DP size, and migration. Means of distributions are marked as points, and medians are highlighted as lines through violins.

**Figure S45:** The effect of demographic treatments on the generation DP reached its phenotypic optimum (trait value > 9) for $\mu_\sigma$ = 0.1 simulations. Violins denote distributions calculated over all iterations of genome windows, organised according to founding bottlenecks, DP size, and migration. Means of distributions are marked as points, and medians are highlighted as lines through violins.

**Figure S46:** Effects of demographic treatments on measures of genetic divergence across all sampling generations for simulations without expansions (Total N = 1000). Point colour and shape denote treatment groups for founding bottlenecks (**A**), protracted bottlenecks (**B**) and migration (**C**). Each point represents values of divergence averaged across all genes within individual treatments groups, averaged within treatment levels, and averaged over 20 iterations.

**Figure S47:** Effects of demographic treatments on the relationship between selection and measures of genetic divergence across all sampling generations for simulations without expansions (Total N = 1000). Point colour and shape denote treatment groups for founding bottlenecks (**A**), protracted bottlenecks (**B**) and migration (**C**). Each point represents correlation coefficients calculated across all genes within individual treatments groups, averaged within treatment levels, and averaged over 20 iterations.

**Figure S48:** Effects of demographic treatments on the relationship between selection and measures of genetic divergence across all sampling generations for simulations without expansions (Total N = 1000). Point colour and shape denote treatment groups for founding bottlenecks (**A**), protracted bottlenecks (**B**) and migration (**C**). Each point represents correlation coefficients calculated across all genes within individual treatments groups, averaged within treatment levels, and averaged over 20 iterations.
