## Supporting Information for "Contingent Convergence: The ability to detect convergent genomic evolution is dependent on population size and migration"

### Supporting information, Whiting and Fraser 2019 – Modifying simulation parameters

#### Reducing $\mu_\sigma$ from 1.0 to 0.1

In terms of absolute divergence (Figure S40), compared with larger mutation effect sizes, the effects of demography were largely similar. Thus, DP size and migration modify measures in the same way regardless of mutation effect size. What we do see however is variation in the shape of divergence curves through time. Notably, both $F_{ST}$ and $\Delta\pi$ are elevated to begin with. This may be because with smaller effect sizes, burn-in populations harbour more genetic variation, such that when populations split there are more rapid allele frequency changes and notable divergence by 100 generations. This allele change may occur rapidly because of the replacement of the phenotypic optimum with the derived optimum, causing the more variable DP population to drift around phenotype space. Further, we no longer see peaks of $\Delta\pi$ that we attribute to selective sweeps occurring with large effect mutations. This agrees with the notion that hard sweeps are restricted to loci of large effect.

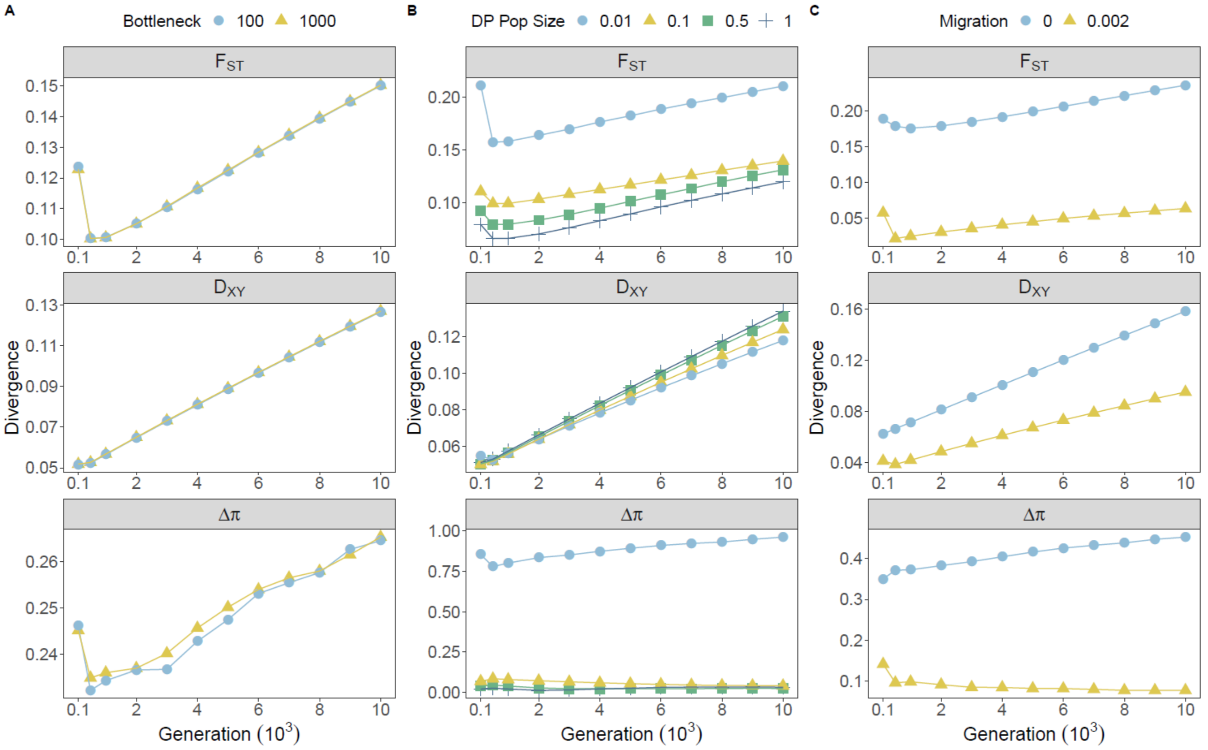


**Figure S40:** Effects of demographic treatments on measures of genetic divergence across all sampling generations for $\mu_\sigma$ = 0.1 simulations. Point colour and shape denote treatment groups for founding bottlenecks (**A**), protracted bottlenecks (**B**) and migration (**C**). Each point represents values of divergence averaged across all genes within individual treatments groups, averaged within treatment levels, and averaged over 20 iterations.

In contrast to simulations with larger effect, we fail to recover any real correlations with our selection parameter (*S*) (Figure S41). Correlations are so weak that effects of demography are negligible. This highlights that in our main results, correlations with selection are driven by interactions with large effect loci, rather than variation generally (although some variation will be linked to large effect loci).

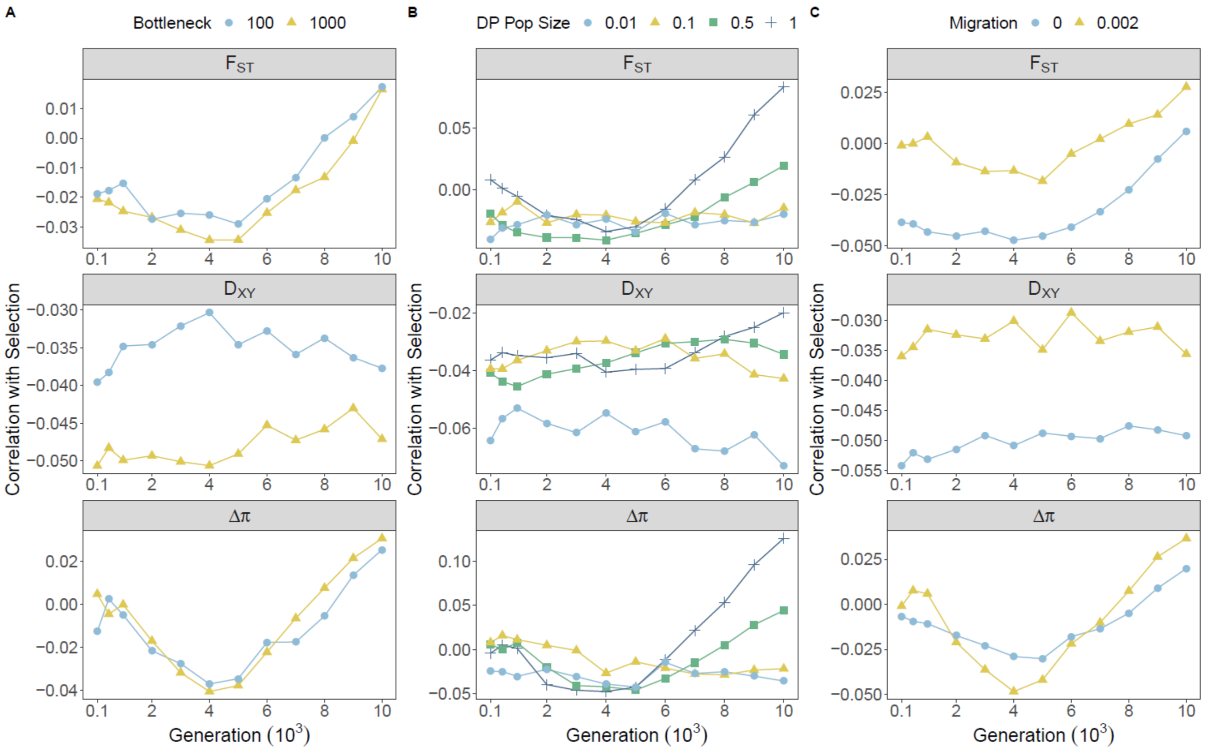


**Figure S41:** Effects of demographic treatments on the relationship between selection and measures of genetic divergence across all sampling generations for $\mu_\sigma$ = 0.1 simulations. Point colour and shape denote treatment groups for founding bottlenecks (**A**), protracted bottlenecks (**B**) and migration (**C**). Each point represents correlation coefficients calculated across all genes within individual treatments groups, averaged within treatment levels, and averaged over 20 iterations.

Despite weak correlations with selection, we still find that $Pheno_{Div}$ distributions are positively shifted relative to $Pheno_{Null}$ and Neutral simulations (Figure S42). This suggests that divergent selection is still able to modify divergence in the tail ends of distributions despite a weak influence over windows generally. Further, this effect is still predominantly driven by connectivity of AP and DP. Interestingly, without migration we find that divergent selection actually shifts $Pheno_{Div}$ distributions negatively.

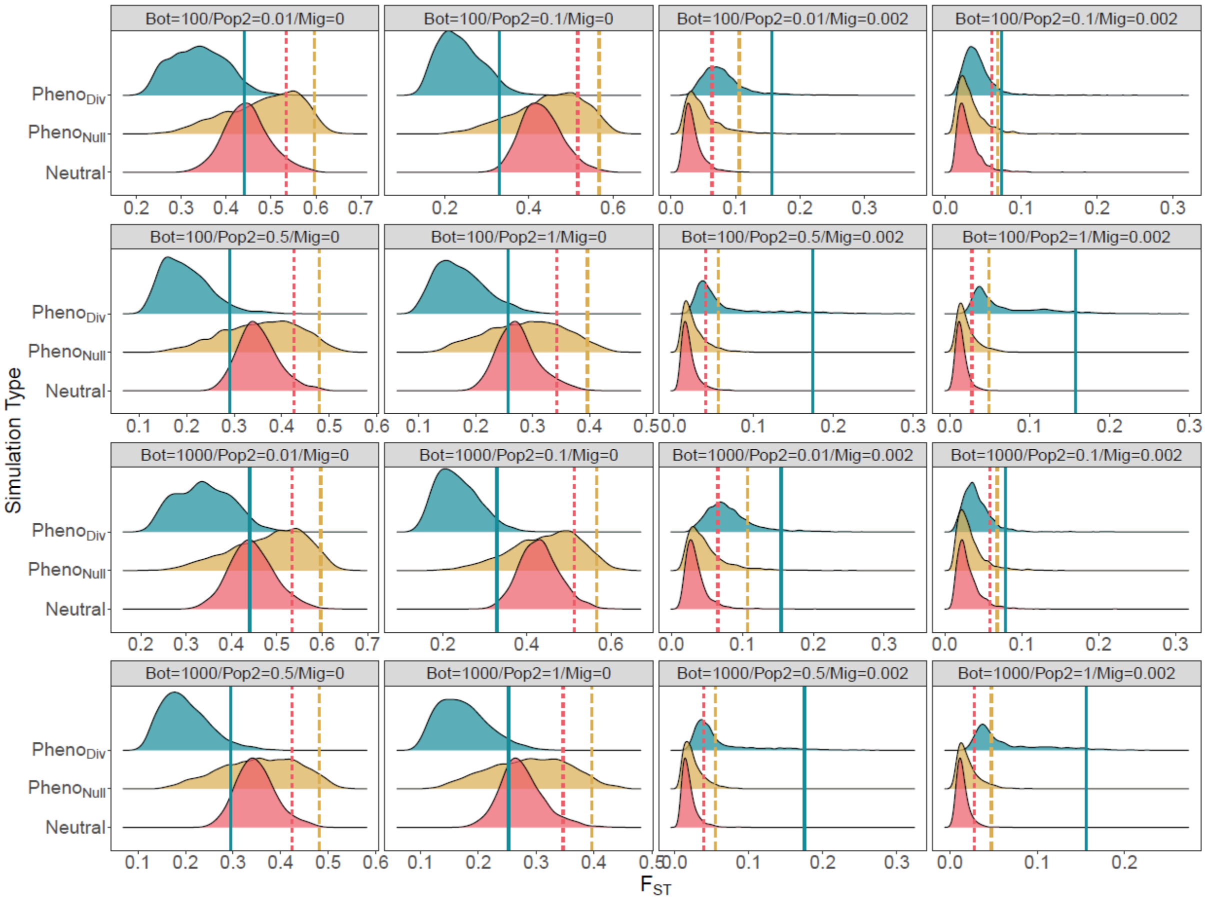


**Figure S42:** Distributions of $F_{ST}$ under each of the 16 unique demographic treatments under three selection regimes: $Pheno_{Div}$ (divergent selection), $Pheno_{Null}$ (stabilising selection) and Neutral, after 10,000 generations for $\mu_\sigma = 0.1$ simulations. Upper 5% quantiles are highlighted for each distribution, with linetype corresponding to selection: Solid = Divergent, Dashed = Stabilising, Dotted = Neutral. Each distribution represents data pooled from 20 iterations of 100 gene windows (N = 2000).

Correlations between measures (Figure S43), particularly $F_{ST}$ and $D_{XY}$ were similar to those observed for neutral simulations, consistent with a minimal influence of selection generally. Unlike for neutral simulations however, we failed to observe any emergence of negative correlations between $F_{ST}$ and $\Delta\pi$ that occur under neutrality. Demographic effects were broadly similar.

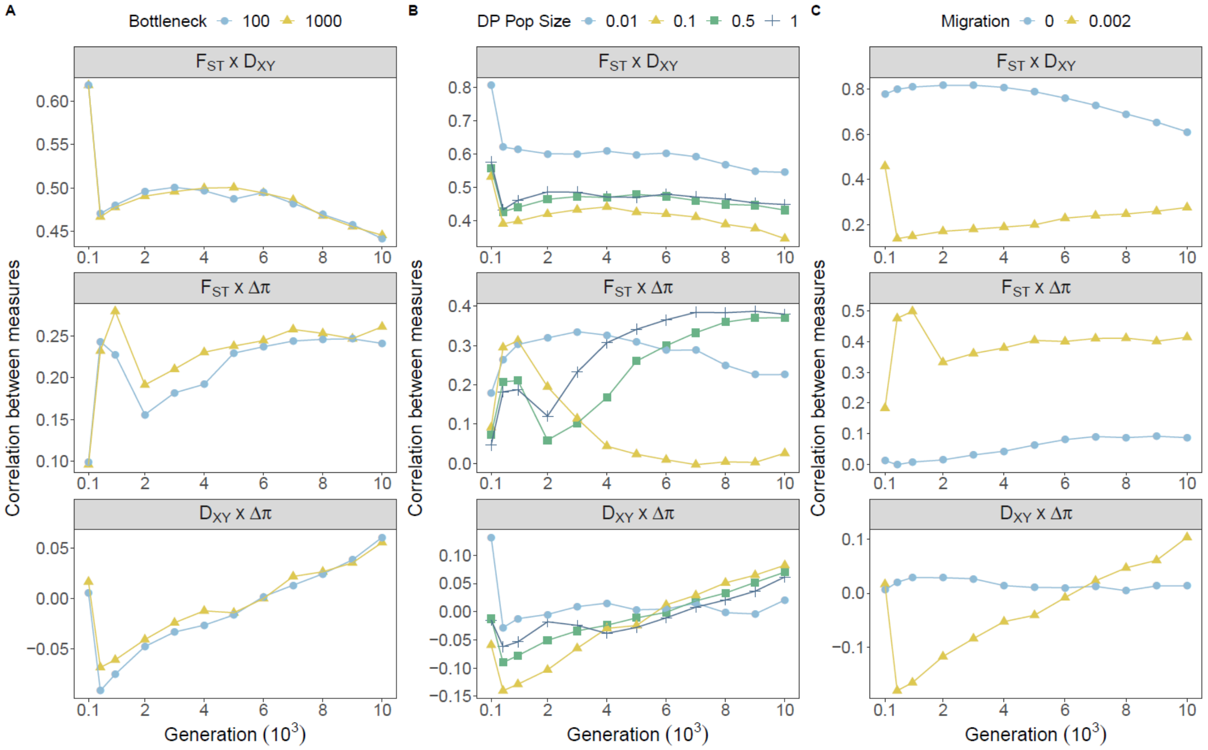


**Figure S43:** Effects of demographic treatments on the relationship between measures of genetic divergence across all sampling generations for $\mu_\sigma = 0.1$ simulations. Point colour and shape denote treatment groups for founding bottlenecks (**A**), protracted bottlenecks (**B**) and migration (**C**). Each point represents correlation coefficients calculated across all genes within individual treatments groups, averaged within treatment levels, and averaged over 20 iterations.

Even with reduced mutation effect size, overlapping outliers are still observed and still cluster according to migration treatment (Figure SXX). As for our main results, clusters of outliers for treatments with migration are still stronger for $F_{ST}$ , but are weaker in comparison with mutations of large effect. $D_{XY}$ in contrast shows large amounts of overlap across all demographic treatments, especially those that include migration. This is contrary to similarities between $F_{ST}$ and $D_{XY}$ that are observed under neutrality and with large effect loci.

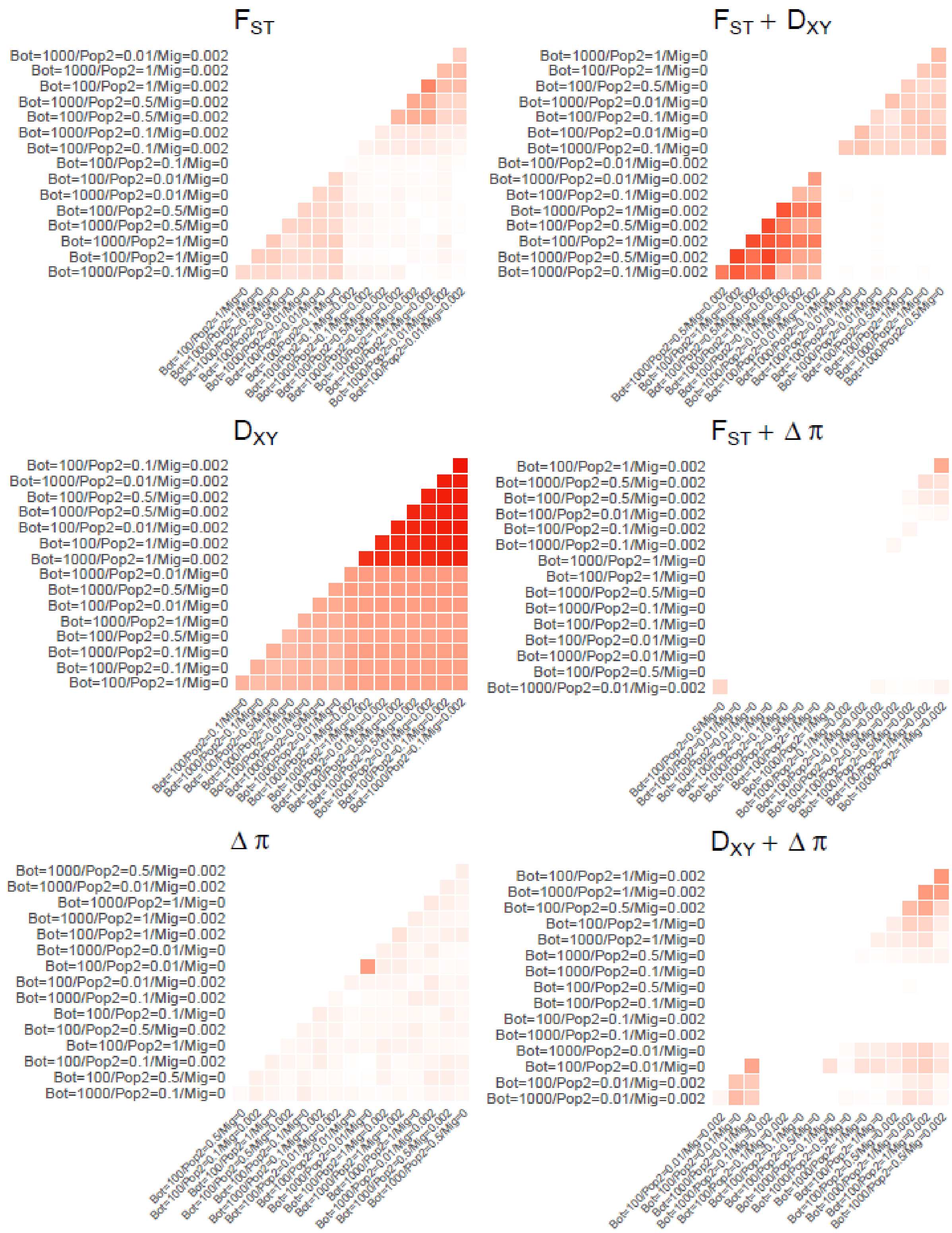


**Figure S44:** The proportional overlap of outliers above the 95% quantile, averaged across 100 downsampled datasets consisting of 95% Neutral and 5% $Pheno_{Div}$ data for each of the 16 demographic treatments after 10,000 generations for $\mu_\sigma = 0.1$ simulations. Axis orderings were determined through hierarchical clustering. Heatmaps are shown for single measures of $F_{ST}$, $D_{XY}$ and $\Delta\pi$ in the first column, and combined measures in the second column. Heatmaps are coloured according to a common scale of 0 to 1. Treatments are labelled with founding bottleneck (Bot), DP population size (Pop2), and migration (Mig) values.

Some of the discrepancies between large-effect, small-effect and neutral loci, may stem from temporal variation in models. Expectedly, it is apparent that without loci of larger effect we observe substantial variation surrounding time taken for DP to reach its phenotypic optimum, with a median generally of ~6000 generations and increasing up to ~9000 under the most extreme reductions in DP size (Figure S45). These results highlight the necessity of including loci of larger effect in our original simulations, given it is less realistic that population may survive several thousand generations away from their phenotypic optimum in nature.

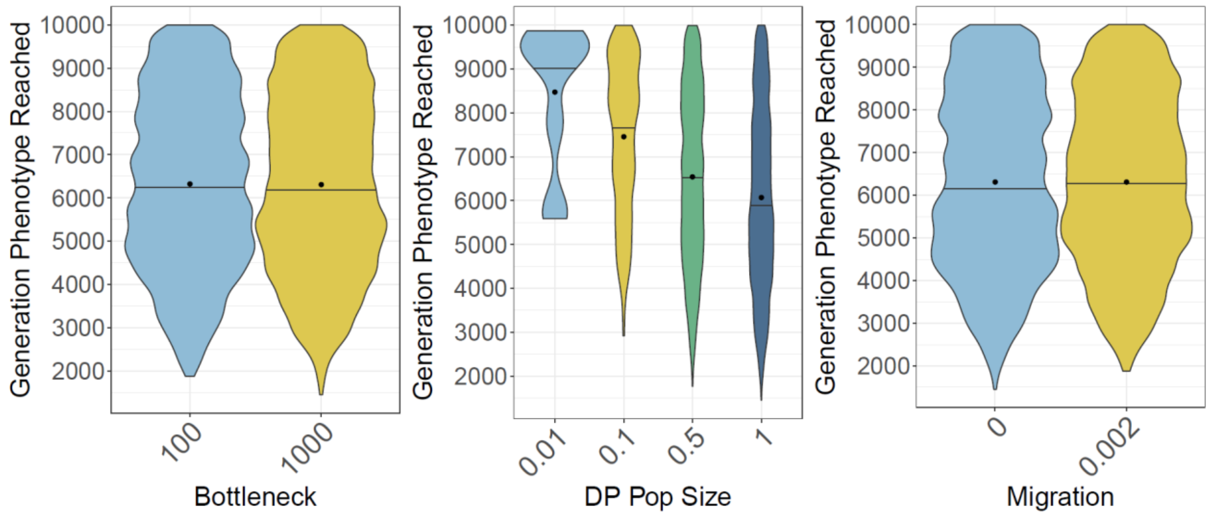


**Figure S45:** The effect of demographic treatments on the generation DP reached its phenotypic optimum (trait value > 9) for $\mu_\sigma = 0.1$ simulations. Violins denote distributions calculated over all iterations of genome windows, organised according to founding bottlenecks, DP size, and migration. Means of distributions are marked as points, and medians are highlighted as lines through violins.

#### Restricting the total N of individuals (AP + DP) to 1000

Given our simulations involve a burn-in population of 1000 individuals, which is then used to found AP (N = 1000) and DP (N = 10-1000), there is potential for a population expansion of the metapopulation. Population expansions may cause shifts in genetic variation through processes such as allele surfing. To investigate this potential we modified simulations to involve a burn-in population of 1000 individuals that is then sub-divided into AP and DP populations according to: AP/DP = 990/10, 900/100, 750/250, 500/500. The results for correlation analyses when running the simulations in this way are presented below (Figure S46-48), with little to differentiate between our main results.

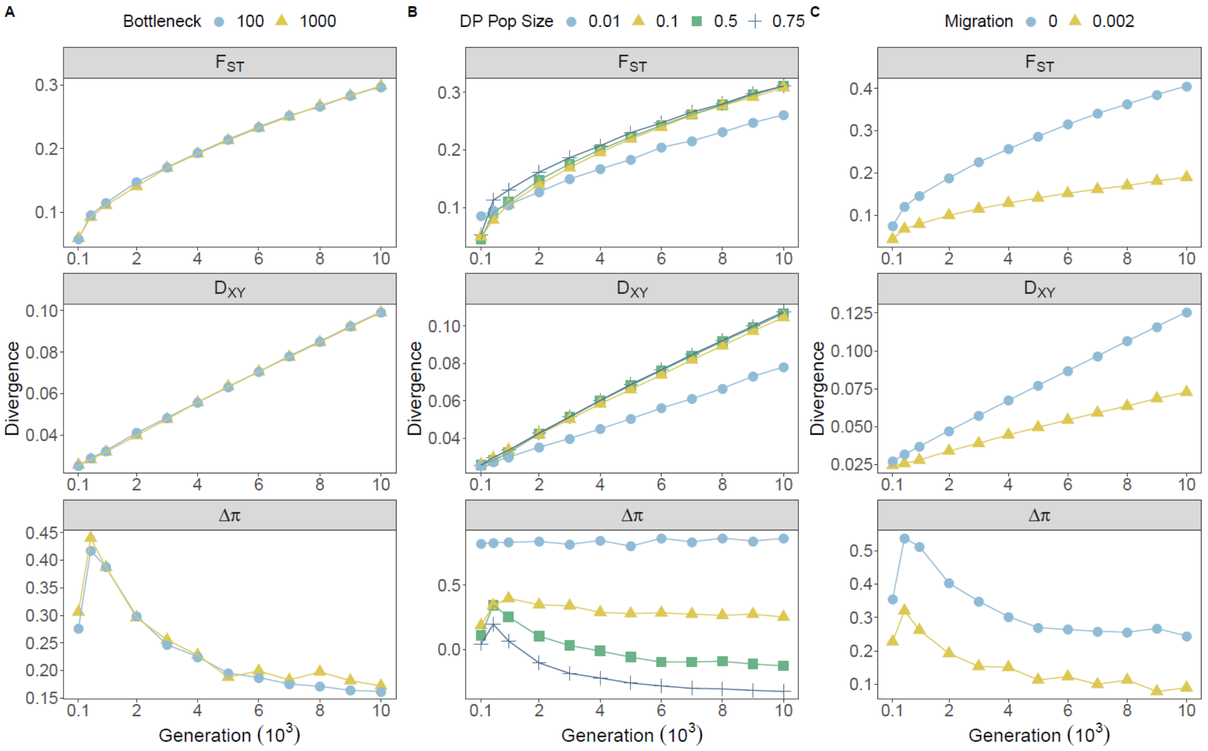


**Figure S46:** Effects of demographic treatments on measures of genetic divergence across all sampling generations for simulations without expansions (Total N = 1000). Point colour and shape denote treatment groups for founding bottlenecks (**A**), protracted bottlenecks (**B**) and migration (**C**). Each point represents values of divergence averaged across all genes within individual treatments groups, averaged within treatment levels, and averaged over 20 iterations.

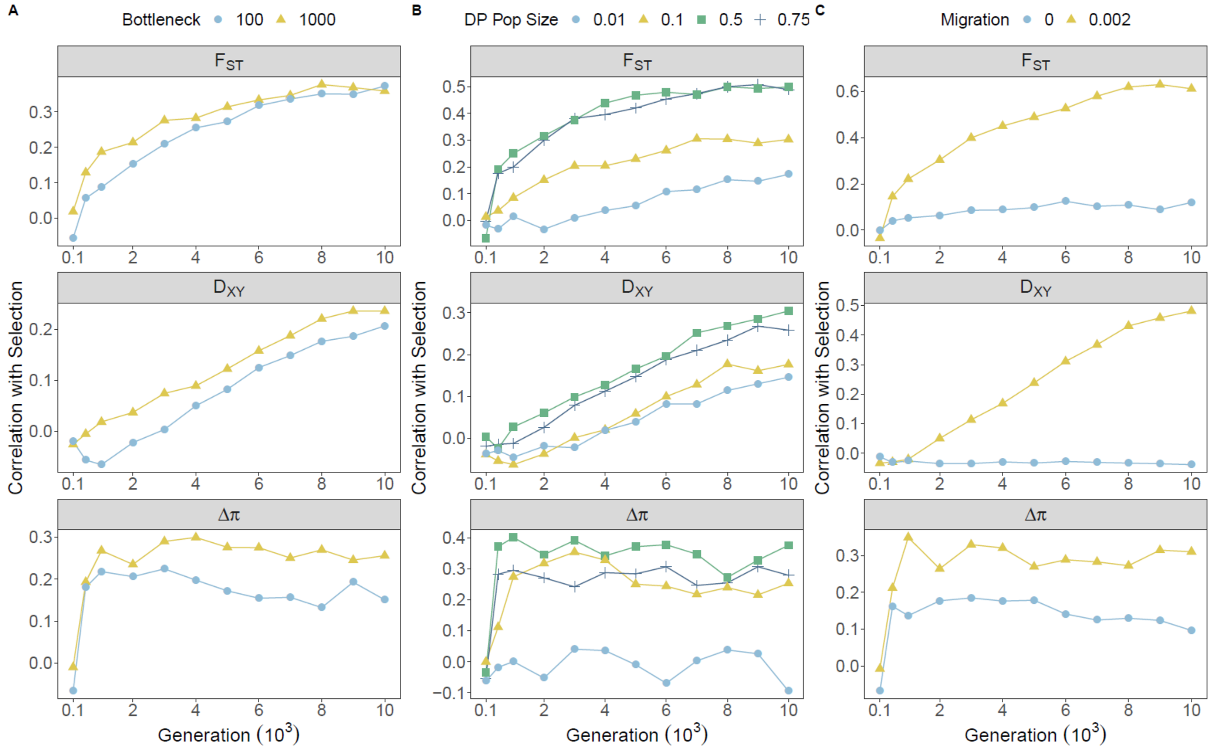


**Figure S47:** Effects of demographic treatments on the relationship between selection and measures of genetic divergence across all sampling generations for simulations without expansions (Total N = 1000). Point colour and shape denote treatment groups for founding bottlenecks (**A**), protracted bottlenecks (**B**) and migration (**C**). Each point represents correlation coefficients calculated across all genes within individual treatments groups, averaged within treatment levels, and averaged over 20 iterations.

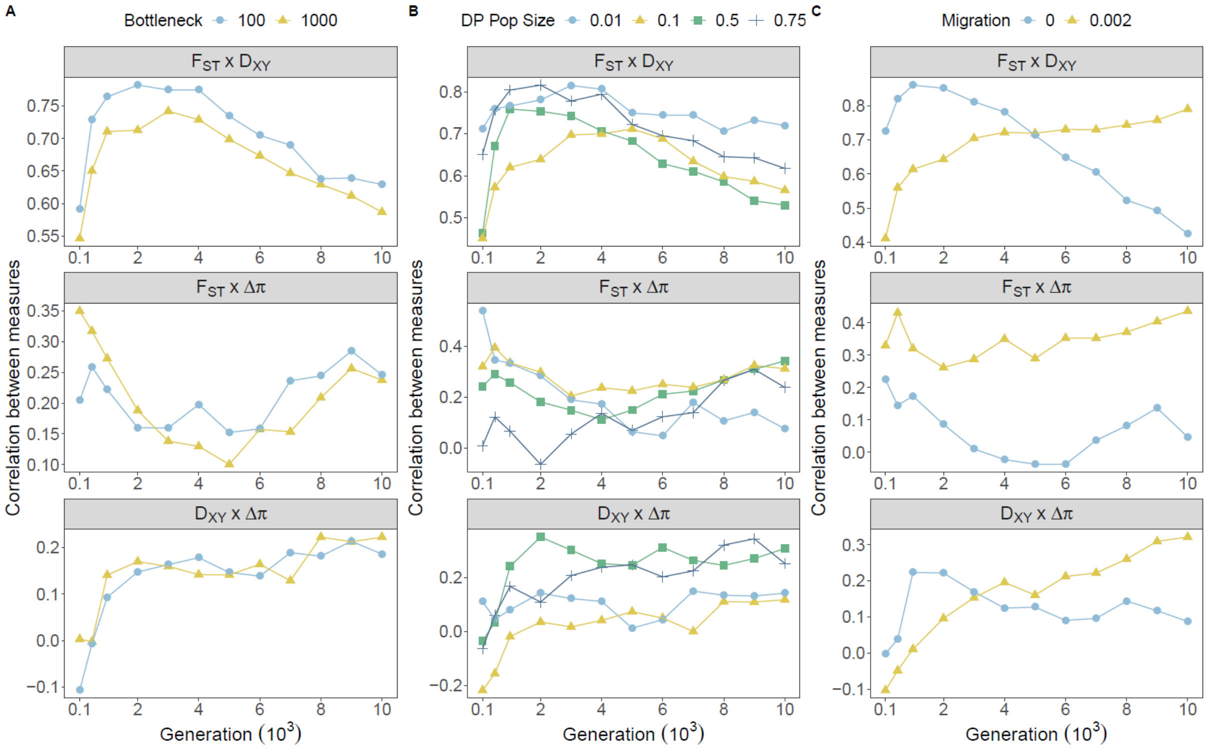


**Figure S48:** Effects of demographic treatments on the relationship between selection and measures of genetic divergence across all sampling generations for simulations without expansions (Total N = 1000). Point colour and shape denote treatment groups for founding bottlenecks (**A**), protracted bottlenecks (**B**) and migration (**C**). Each point represents correlation coefficients calculated across all genes within individual treatments groups, averaged within treatment levels, and averaged over 20 iterations.
